# Prenatal cannabis exposure affects human fetal neurodevelopment: an integrated multi-omics study

**DOI:** 10.64898/2026.07.15.738779

**Authors:** Guenther Kahlert, Xin Chen, Theo K. Bammler, James W. MacDonald, Lyndsey S. Benson, Ian A. Glass, Jennifer C. Dempsey, Dilip Kumar Singh, Bhagwat Prasad, Jashvant D. Unadkat

## Abstract

Prenatal cannabis use is on the rise, and observational studies suggest that such use results in neurodevelopmental deficits in the offspring. Because observational studies can be confounded by unaccounted factors, we studied the neurodevelopmental consequences, at the molecular level, of prenatal cannabis use. We applied an integrated multi-omics approach, combining transcriptomics and global proteomics, to first trimester (T1) and second trimester (T2) human fetal brains from pregnancies with and without documented maternal cannabis exposure and no use of drugs of abuse. Prenatal cannabis exposure produced minimal molecular effects in female T1 fetal brains but induced pronounced system-level disruption in male T2 fetal brains. These disrupted pathways have molecular signatures linked to neurodevelopmental and neuropsychiatric disorders, including autism spectrum disorder, schizophrenia-related pathology, and disorders of cortical connectivity, raising significant concerns of prenatal cannabis use.

## Introduction

Cannabis use during pregnancy has increased alongside expanding legalization and persistent perception of cannabis as a low-risk substance. This is despite public health warnings against prenatal cannabis use. About 7% of pregnant women use cannabis with the highest prevalence in the first trimester (10%) (*1*). The primary reason for use is to relieve stress or anxiety (81.5%), nausea/vomiting (77.8%), and pain (55.1%) (*2*).

Observational studies have linked prenatal cannabis exposure to adverse neurodevelopmental outcomes in offspring, including cognitive deficits, attention-related disorders, and increased autism-spectrum disorder risk (*3, 4*). Animal studies suggest that these neurodevelopmental effects of cannabis use are sex-dependent (*5*). Observational studies remain susceptible to uncontrolled confounding factors. For example, the Ontario (*6*) and ECHO (*7*) cohorts reached opposing conclusions regarding the association between prenatal cannabis exposure and autism spectrum disorder in the offspring. Because randomized clinical trials would be unethical, observational studies cannot establish definitive causal relationships between prenatal cannabis exposure and neurodevelopmental outcomes in the offspring.

Cannabis contains the intoxicating cannabinoid, Δ^9^-tetrahydrocannabinol (Δ^9^-THC). Δ^9^-THC is metabolized in humans to 11-OH-THC which is also intoxicating (*8*). Δ^9^-THC and 11-OH-THC exert their intoxicating effects through the cannabinoid receptor 1 (CB1), a key component of the endocannabinoid system (ECS) that is highly expressed during human cortical development (*9*). Endocannabinoid signaling functions as an intrinsic regulator of early brain development, coordinating neuronal differentiation, axon guidance, synaptic assembly, and circuit formation (*10, 11*). Through coupling to intracellular signaling pathways, CB1 activity regulates cytoskeletal dynamics, vesicle trafficking, neuronal polarity, and metabolic programs central to neuronal identity and connectivity (*12*). Experimental studies in animal models and *in vitro* human cell models show that exogenous cannabinoid exposure can perturb these processes relevant to neurodevelopment (*9, 13*). However, these findings are limited by species-specific differences in developmental timing and cortical organization, as well as exposure in these models to supra-pharmacological Δ^9^-THC concentrations. As a result, whether human-relevant fetal exposure to cannabinoids disrupts developmental programs in the human fetal brain remains unresolved.

To quantify human-relevant exposure to both Δ^9^-THC and 11-OH-THC, we first determined if these cannabinoids cross the perfused human placenta (*14*). Then, we quantified human fetal exposure to Δ^9^-THC and 11-OH-THC by measuring them in both the fetal circulation (at term) and in 1^st^ (T1) and 2^nd^ trimester (T2) fetal tissues, including the brain, after prenatal cannabis exposure. The fetal brain exposure of Δ^9^-THC, relative to maternal plasma exposure, was about 45–50% (*15*). Moreover, we developed a physiologically based pharmacokinetic model to predict for any dose and route of cannabis administration the fetal circulation and fetal tissue exposure to Δ^9^-THC and 11-OH-THC from 15 weeks of gestation to term (*15*). These findings established the pharmacological Δ^9^-THC and 11-OH-THC concentrations that the developing human fetal brain is exposed to during critical windows of neurodevelopment.

As a follow-up to the above studies, here we investigated if prenatal cannabis exposure causes molecular perturbations in the developing human fetal brain that provide insights into long-term sequelae to the offspring. To do so, we applied an integrated multi-omics approach, combining transcriptomics and global proteomics, and leveraged the above unique collection of T1 and T2 fetal brain samples from pregnancies with and without prenatal cannabis exposure and no use of other drugs of abuse (documented by urine toxicology).

## Results

The subjects enrolled were: 63.9 ± 10.7, 67.7 ± 9.6, 101.0 ± 16.3, and 99.4 ± 8.6 days (mean ± SD) of gestation for T1 controls, T1 cases, T2 controls and T2 cases, respectively (Table S1). They were representative of ethnicity and race in the Seattle metropolitan area. Cannabis was used mostly by inhalation, although 6 of 28 case subjects (∼21%) also reported combined use by inhalation and ingestion. Based on those who reported the quantity of last cannabis use, the estimated THC-equivalent dose (assuming 15% Δ^9^-THC by weight in cannabis plant material) was 183 ± 145 mg per use in T1 participants (n = 13 of 15) and 161 ± 138 mg in T2 participants (n = 7 of 12).

Cannabis use frequency ranged from 2–3 times per month to three or more times per day in T1, and from once per month or less to three or more times per day in T2 (Table S1). As cannabis use history was self-reported, these dose estimates should be interpreted with caution.

A total of 43 participant-derived samples were included for multi-omics analysis (Fig. 1; Table S1). Transcriptomics quantified 23,206 expressed genes (n=34), and DIA proteomics quantified 8,233 proteins (n=36). Twenty-seven samples (T1, n = 15; T2, n = 12) were profiled by both modalities. Fetal sex distribution differed between groups (Fig. 2A) with predominately females in T1 controls and predominately males in T2 cases.

**Figure 1.**
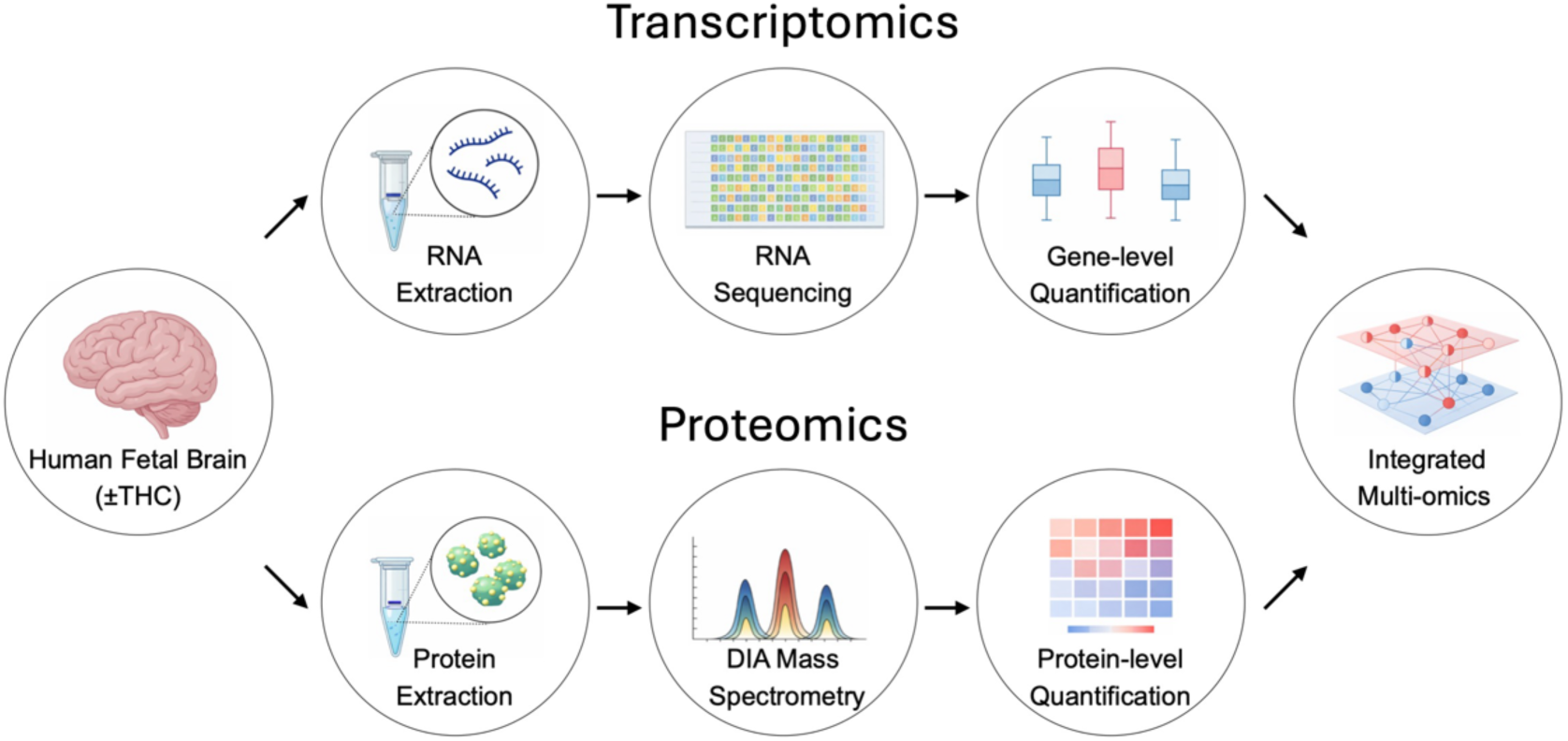
**Multi-omics workflow**. Schematic overview of the integrated multi-omics workflow applied to human fetal brain tissue. RNA sequencing was performed for gene-level quantification, and global proteomic profiling was performed using DIA mass spectrometry for protein-level quantification. Transcriptomic and proteomic datasets were subsequently integrated to enable multi-omics analysis.

**Figure 2.**
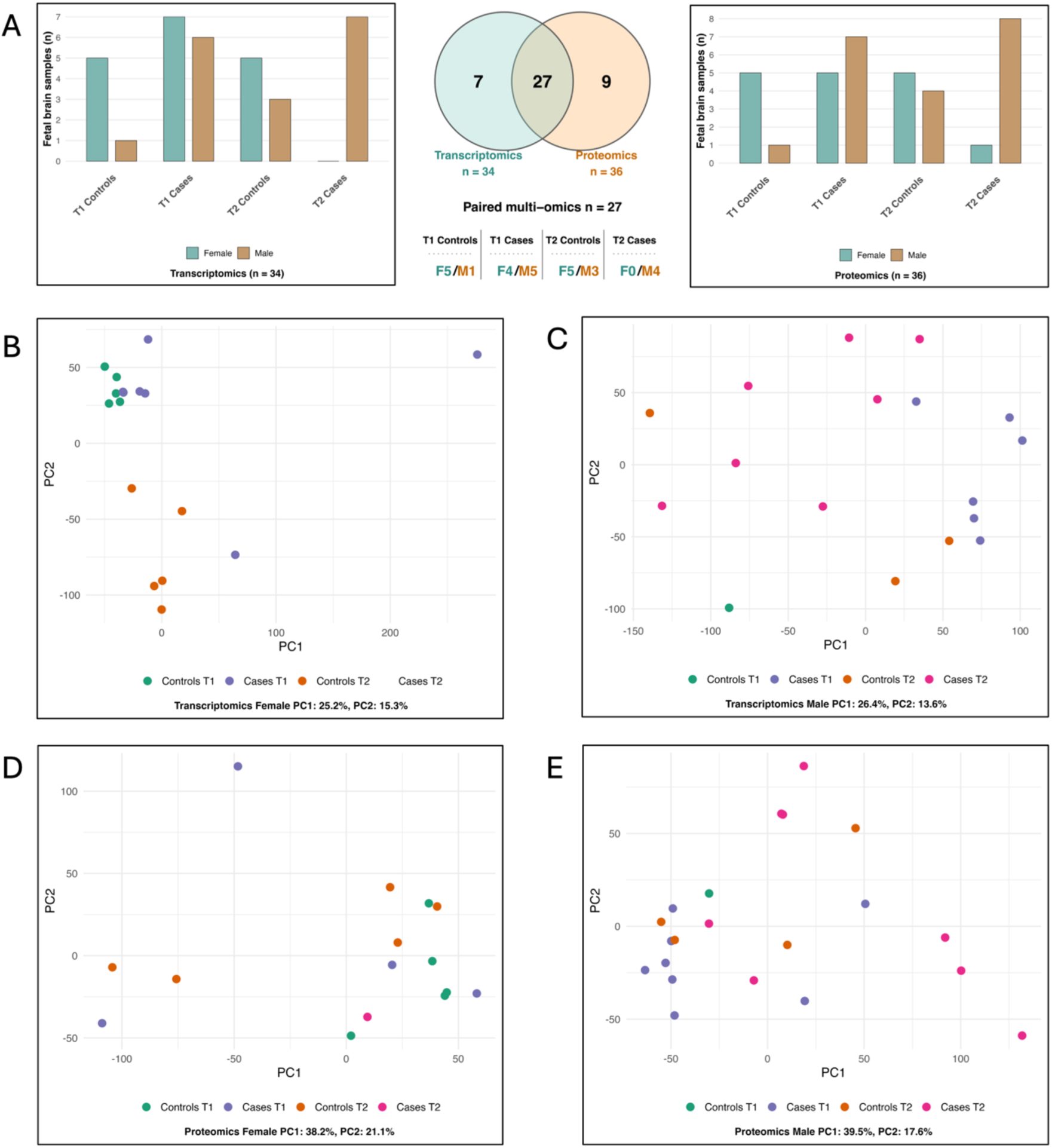
Cohort composition and global molecular structure of the fetal brain multi-omics dataset. (**A**) Composition of the fetal brain cohorts analyzed by transcriptomics and proteomics, stratified by trimester (T1 and T2), exposure status (cases and controls), and sex. Bar plots indicate the number of female and male samples included in each comparison. The central Venn diagram summarizes the overlap between transcriptomic and proteomic datasets and identifies the subset of paired multi-omics samples used for integrated analyses (n = 27), with the distribution of paired multi-omics samples across trimester, cannabis exposure group, and sex indicated below. F, female; M, male. (**B**) PCA of normalized female fetal brain transcriptomic data. Samples segregated primarily by trimester, with PC1 and PC2 explaining 25.2% and 15.3% of the variance, respectively. (**C**) PCA of normalized male fetal brain transcriptomic data.

Principal component analysis (PCA) of transcriptomic and proteomic datasets was performed in sex-independent (Fig. S1A–B) and sex-stratified manner (Fig. 2B–E). In both datasets, samples segregated primarily by trimester rather than cannabis exposure, identifying fetal brain development as the dominant source of variation. Given known sex-dependent regulation of the brain proteome (*16*), and reported sex-specific effects of cannabis exposure (*17*), subsequent analyses prioritized sex-stratified comparisons. The few RNAlater-stabilized samples did not segregate in proteomic PCA analyses.

Samples showed separation primarily by trimester and cannabis exposure status, with PC1 and PC2 explaining 26.4% and 13.6% of the variance, respectively. (**D**) PCA of normalized female fetal brain proteomic data. Samples showed broad dispersion without clear separation by trimester or cannabis exposure status, with PC1 and PC2 explaining 38.2% and 21.1% of the variance, respectively. (**E**) PCA of normalized male fetal brain proteomic data. Samples showed broad dispersion without clear separation by trimester or exposure status, with PC1 and PC2 explaining 39.5% and 17.6% of the variance, respectively. No obvious segregation by estimated THC dose or reported cannabis use frequency (Table S1) was observed in these PCA analyses (Fig. 2B–E).

### Prenatal cannabis exposure induced coordinated disruption of neuronal and metabolic systems in the T2 male fetal brains

A total of 481 differentially expressed genes (DEGs) and more than 1,400 differentially expressed proteins (DEPs) were identified in the T2 Cases vs. Controls comparison (Fig. S2–S3), indicating pronounced proteomic remodeling. Concordance analysis identified 80 significantly concordant transcript–protein pairs shared across both modalities (Fig. 3A). As discussed below, these DEGs and DEPs were enriched for regulators of cortical development, synaptic organization, and cellular metabolism. Interaction network analysis linked these alterations to disruption of synaptic structure, circuit specification, and cellular homeostasis in the developing male T2 fetal brains.

**Figure 3.**
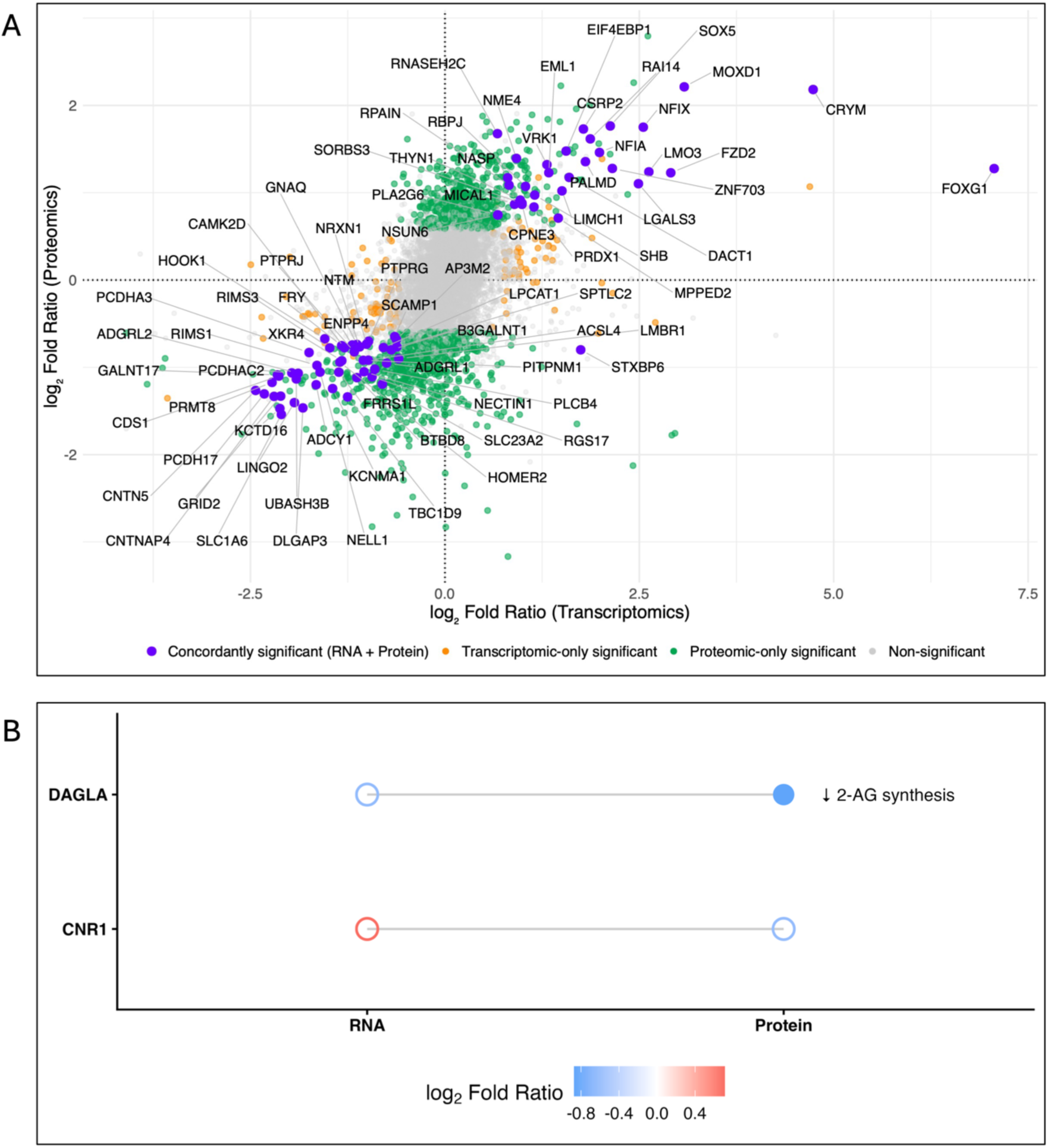
**Coordinated disruption of neuronal and metabolic systems in T2 male fetal brains following prenatal cannabis exposure**. (**A**) Multi-omics concordance of significant transcriptomic and proteomic changes in the male T2 Cases vs. Controls comparison. Points are colored by significance for both RNA and protein (blue), transcriptomic-only (orange), proteomic-only (green), or non-significant (gray). (**B**) Transcriptomic and proteomic expression changes of the endocannabinoid signaling components. CNR1 (CB1 receptor) expression was not significantly altered, whereas DAGLA exhibited reduced protein abundance, suggesting reduced endogenous 2-AG production. Color denotes log₂ fold ratio; filled and open circles indicate significant change (adj. P < 0.05) or no significant change. Lines connect RNA and protein data.

In the combined T2 cohort without sex stratification, transcriptomic analysis identified 468 DEGs (Fig. S4), whereas proteomic analysis identified no DEPs. The loss of DEPs in the combined cohort likely reflects attenuation of sex-specific proteomic signals after combining males and females rather than absence of molecular effects. This is consistent with evidence that human brain protein abundance is strongly sex-dependent (*16*).

### Effects on the endocannabinoid system

The CB1 receptor (CNR1) did not show a consistent cannabis exposure effect, with small, non-significant, and discordant RNA and protein changes across datasets (Fig. 3B). In contrast, the protein expression of diacylglycerol lipase α (DAGLA), the principal enzyme for 2-arachidonoylglycerol (2-AG) production, was significantly reduced, suggesting reduced CB1 receptor ligand availability. This reduction was not observed at the transcript level. Together, these findings indicate prenatal cannabis exposure to the exogenous CB1 ligand, Δ^9^-THC, disrupts endocannabinoid signaling despite preserved CB1 receptor expression.

### Concordant suppression of synaptic scaffold and vesicle machinery

Cannabis exposure in T2 male fetuses showed concordant reduction in core synaptic organization, reflected by coordinated downregulation of synaptic scaffold and vesicle-cycle components. This reduction was centered on concordantly reduced transcript– protein pairs, including NRXN1, a presynaptic adhesion molecule required for synapse formation, and presynaptic and postsynaptic scaffold-associated proteins RIMS1, RIMS3, DLGAP3, and HOMER2 (Figs. 3A and S5), suggesting altered synaptic architecture and organization (*18*). This reduction extended to presynaptic vesicle systems, with concordant downregulation of SCAMP1 and AP3M2 (Figs. 3A and S5). STXBP6 showed bidirectional discordance, with increased transcript abundance but reduced protein levels (Figs. 3A and 4A), whereas VAMP1 displayed proteomic downregulation without a corresponding transcriptomic signal (Fig. 4A). Proteomic downregulation further extended across vesicle fusion systems, including STX1B, VAMP1/2, UNC13A, and RAB3A, indicating impaired neurotransmitter release (*19*). Interaction network analysis identified transcriptomic and proteomic modules linked to glutamatergic synapse organization, vesicle fusion, synaptic signaling, and vesicle transport (Figs. S6–S7), consistent with impaired coupling between synaptic structure and neurotransmission.

**Figure 4.**
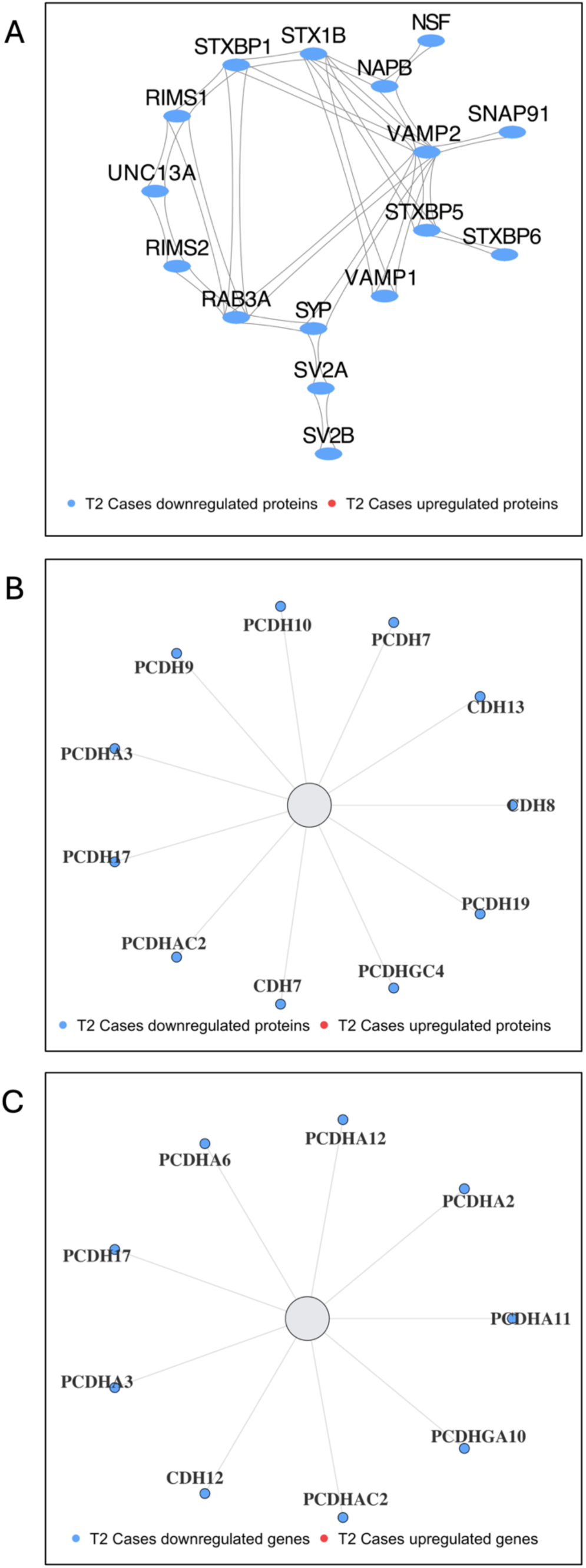
Proteomic and transcriptomic downregulation of synaptic vesicle and cell adhesion networks in T2 male fetal brains following prenatal cannabis exposure. (**A**) STRING network of significantly altered presynaptic vesicle fusion and release proteins in male T2 Cases vs. Controls. Nodes represent proteins meeting differential abundance criteria (adjusted P < 0.05, |log₂FR| ≥ 0.58); edges indicate protein–protein interactions; colors denote the direction of significant change. (**B**) Proteomic and (**C**) transcriptomic representations of protocadherin and cadherin family members in male T2 Cases vs. Controls. Nodes represent gene transcripts or proteins grouped by family; colors denote the direction of significant change. Filled nodes indicate significant changes (adjusted P < 0.05, |log₂FR| ≥ 0.58).

### Disruption of neuronal adhesion systems and circuit specification

Cannabis exposure of T2 male fetuses induced concordant downregulation of neuronal adhesion systems, including contactin networks and protocadherin-mediated circuit specification pathways (Fig. 3A, Fig. S5). Multi-omics concordance analysis identified reduced CNTN5 and CNTNAP4 together with protocadherin family members including PCDH17, PCDHA3, and PCDHAC2, indicating impaired neuronal identity encoding and synaptic specificity (*20*). Additional reductions were observed in neuronal adhesion molecules including NECTIN1 and NTM. Broader transcriptomic and proteomic reductions extended across multiple protocadherin and cadherin family members (Fig. 4B–C).

Interaction network analysis identified transcriptomic and proteomic modules linked to transsynaptic adhesion systems and neuronal wiring, including latrophilin signaling components (Figs. S6–S7) (*21*). Consistent with prior Δ^9^-THC studies in human iPSC-derived neurons, these changes were associated with suppression of synaptic and glutamatergic signaling pathways (*22*), indicating impaired circuit refinement and neuronal network formation.

*System-level disruption of neuronal excitability and membrane transport capacity* Prenatal cannabis exposure in T2 male fetuses induced concordant reduction of neuronal signaling systems, including GRID2 and SLC1A6 (EAAT4) (Figs. 3A and S5), indicating impaired glutamatergic signaling and synaptic responsiveness (*23*). Reduced transcript– protein abundance of KCNMA1 indicates altered membrane excitability. Downregulation of ADCY1, PLCB4, and GNAQ (Figs. 3A and S5) further indicates impaired GPCR- and Ca²⁺-dependent signaling, consistent with reduced neuronal excitability and signal integration in cannabis-exposed T2 male fetal brains. Interaction network analysis identified transcriptomic and proteomic modules linked to glutamatergic synapse organization and GPCR/Ca²⁺ signaling (Figs. S6–7), supporting disrupted neuronal excitability and intracellular signal coupling.

### Effects on transporters

Concordant transcript–protein reduction of SLC1A6 (EAAT4) and SLC23A2 (SVCT2) indicates impaired glutamatergic transport and intracellular ascorbate homeostasis (Figs. 3A, S5, and S8). Reduction in protein further extended across neurotransmitter, ion, lipid, amino acid, and metal transport systems, including SLC1A1 (EAAT3), SLC6A1 (GAT1), SLC32A1 (VGAT), and multiple SLC9, SLC27, SLC38, and SLC30 family members (Fig. S8). At the RNA level, selected transporters, including SLC17A7 (VGLUT1), SLC26A4 (Pendrin), and SLCO4C1 (OATP4C1), showed increased expression without corresponding change in protein, indicating a partial compensatory transcriptional response. Reduction in protein extended to ATP-binding cassette transporters, including ABCC1 (MRP1), ABCB8, ABCA5, and ABCA3 (Fig. S8). In contrast, ABCC12 (MRP9) showed transcript-level downregulation without change in protein. Together, these changes indicate broad reduction of membrane transport capacity.

**Figure 5.**
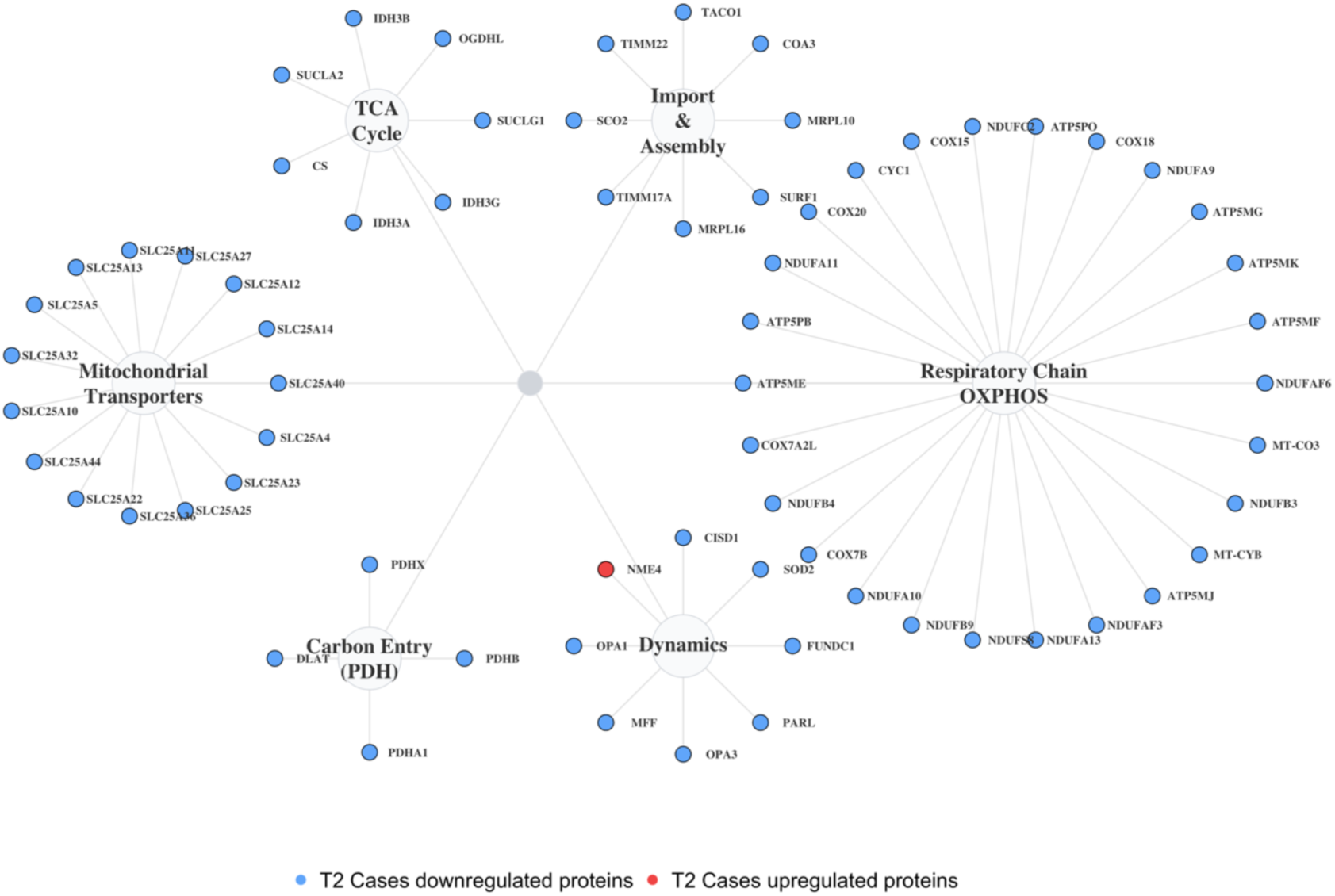
Widespread proteomic disruption of mitochondrial functional networks in T2 male fetal brains following prenatal cannabis exposure. Functional network representation of mitochondrial systems showing coordinated proteomic downregulation in male T2 Cases vs. Controls fetal brains. Nodes represent proteins meeting differential abundance criteria (adjusted P < 0.05, |log₂FR| ≥ 0.58) grouped into mitochondrial functional modules, including respiratory chain/OXPHOS, mitochondrial transporters, import and assembly, carbon entry (PDH), TCA cycle, and dynamics/quality control. Edges indicate curated functional relationships. Node color denotes the direction of significant change.

### Loss of membrane lipid homeostasis and intracellular trafficking integrity

Concordant reduction of lipid metabolism and intracellular trafficking systems (e.g., ACSL4, LPCAT1, SPTLC2, and CDS1) was observed in cannabis-exposed T2 male fetal brains. In contrast, PLA2G6 was increased, consistent with enhanced phospholipid turnover (Figs. 3A and S5). Furthermore, reduction in protein expression included CYP46A1 and CYP4X1, suggesting altered cholesterol and lipid metabolic capacity (Fig. S9). HOOK1 and PITPNM1 were reduced, whereas MICAL1 and SORBS3 were increased, indicating impaired vesicle trafficking with concurrent cytoskeletal remodeling (Fig. 3A, Fig. S5). Interaction network analysis linked these alterations to phospholipid remodeling, membrane lipid metabolism, intracellular trafficking, and membrane organization (Figs. S6 and S7). Collectively, these findings indicate coordinated disruption of membrane lipid homeostasis, intracellular trafficking, and vesicle-associated transport systems in cannabis-exposed T2 male fetal brains.

### Global proteomic depletion of mitochondrial energy systems

Cannabis-exposed T2 male fetal brains showed coordinated proteomic reduction of mitochondrial energy systems (Fig. 5). Electron transport chain components from Complex I (NDUF family), Complex III (CYC1, MT-CYB), Complex IV (MT-CO3, COX family), and ATP synthase (ATP5 family) were broadly decreased, together with respiratory complex assembly factors including NDUFAF family members, SURF1, SCO2, and COA3. Proteins involved in mitochondrial import and translation, including TIMM machinery components, TACO1, and mitochondrial ribosomal proteins, were reduced, indicating impaired respiratory complex maintenance. This reduction further extended to mitochondrial transport and metabolic systems, including fifteen SLC25 transporters, pyruvate dehydrogenase complex components (PDHA1, PDHB, PDHX, DLAT), and multiple TCA cycle enzymes. In contrast, NME4, a cardiolipin-associated mitochondrial membrane protein, was the only mitochondrial protein increased. Proteomic interaction analysis linked these alterations to coherent mitochondrial functional modules without corresponding transcriptomic enrichment (Fig. S7).

### Activation of neuronal lineage specification programs

Prenatal cannabis exposure of T2 male fetal brains induced concordant transcriptomic– proteomic upregulation of neuronal lineage regulators including FOXG1, SOX5, NFIX, EML1, RBPJ, and ZNF703 (Figs. 3A and S5), indicating activation of neurodevelopmental and cortical specification programs (*24*). FOXG1-AS1 was similarly increased at the transcript level (Fig. S10). Increased protein abundance of BCL11B (CTIP2) and SATB2 further indicated concurrent activation of corticofugal and callosal neuronal identity programs during cortical maturation (Fig. S10). Several zinc finger transcription factors were altered at the transcriptomic level (Fig. S11A), with substantially broader upregulation observed at the proteomic level (Fig. S11B), while interaction network analysis identified upregulated modules linked to chromatin remodeling, transcriptional regulation, and RNA processing (Fig. S7).

*Pathway-level analysis reveals systems-level reprogramming in the male T2 fetal brain* Prenatal cannabinoid exposure induced coordinated pathway-level reprogramming in T2 male fetal brains, with proteomic enrichment showing suppression of synaptic and mitochondrial pathways alongside activation of DNA replication, chromatin remodeling, DNA repair, and RNA-processing programs (Figs. 6A–F). Transcriptomic enrichment further revealed suppression of neuronal signaling, cell-cell adhesion, phospholipid metabolism, and G protein-mediated signaling, together with activation of ribosomal, translational, SRP-dependent protein targeting, and EIF2AK4/GCN2-associated stress-response programs (Fig. S12 and S14A–B). Proteomic enrichment resolved these changes into a neuronal systems phenotype characterized by retrograde endocannabinoid signaling, depletion of synaptic vesicle cycling, neurotransmitter transport, glutamatergic and GABAergic synapses, neuroactive ligand–receptor interaction, oxidative phosphorylation, and vesicle-associated secretory functions (Figs. 6A, C, E).

**Figure 6.**
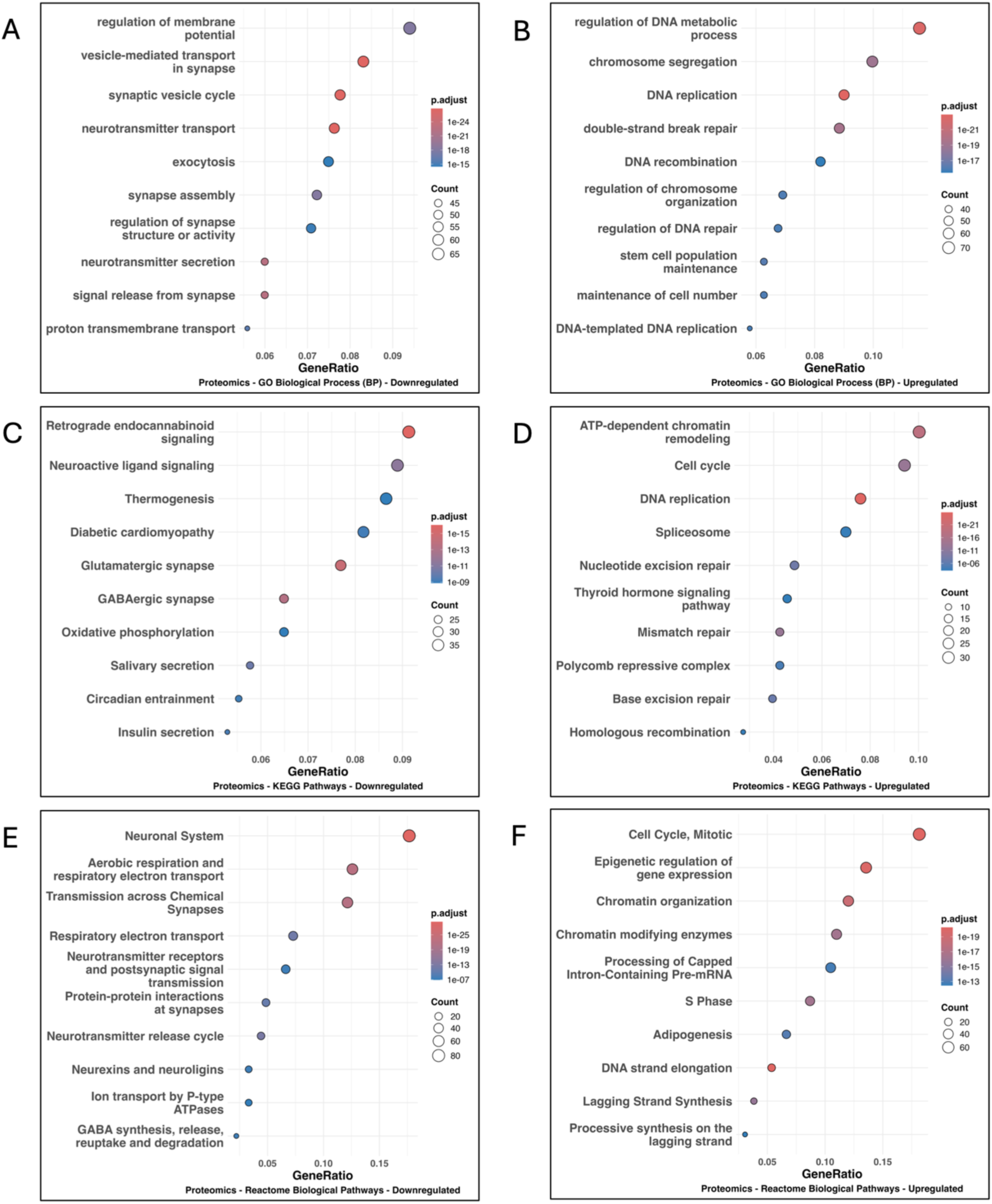
Functional enrichment of proteomic alterations following prenatal cannabis exposure of T2 male fetal brains. Gene Ontology (Biological Process) enrichment analysis of **(A)** downregulated and **(B)** upregulated proteins in the T2 Cases vs. Controls comparison. Dot plots show enriched terms ranked by gene ratio, with point size indicating protein count and color representing adjusted P value. Downregulated proteins are enriched for synapse-related processes, whereas upregulated proteins are enriched for DNA replication, chromosome organization, and DNA repair. KEGG pathway enrichment analysis of **(C)** downregulated and **(D)** upregulated proteins.

Downregulated pathways include synaptic signaling and metabolic processes, whereas upregulated pathways are enriched for cell cycle and DNA repair–associated programs. Reactome pathway enrichment analysis of **(E)** downregulated and **(F)** upregulated proteins. Downregulated pathways are enriched for neuronal system, synaptic transmission, and respiratory electron transport, whereas upregulated pathways are enriched for cell cycle progression, chromatin organization, and RNA processing.

Pathway activity inference using PROGENy identified activation of JAK-STAT, TNFα, and NFκB signaling together with suppression of PI3K-associated trophic signaling and altered steroid hormone pathway activity (Fig. 7). Reactome enrichment further revealed molecular-layer discordance, with suppression of neuronal pathways and translational enrichment at the transcriptomic level, including PLCβ-dependent signaling (Fig. S14C– D), whereas proteomic enrichment showed broad depletion of synaptic transmission, neurotransmitter release, receptor-mediated signaling, aerobic respiration, and electron transport chain pathways (Fig. 6E).

**Figure 7.**
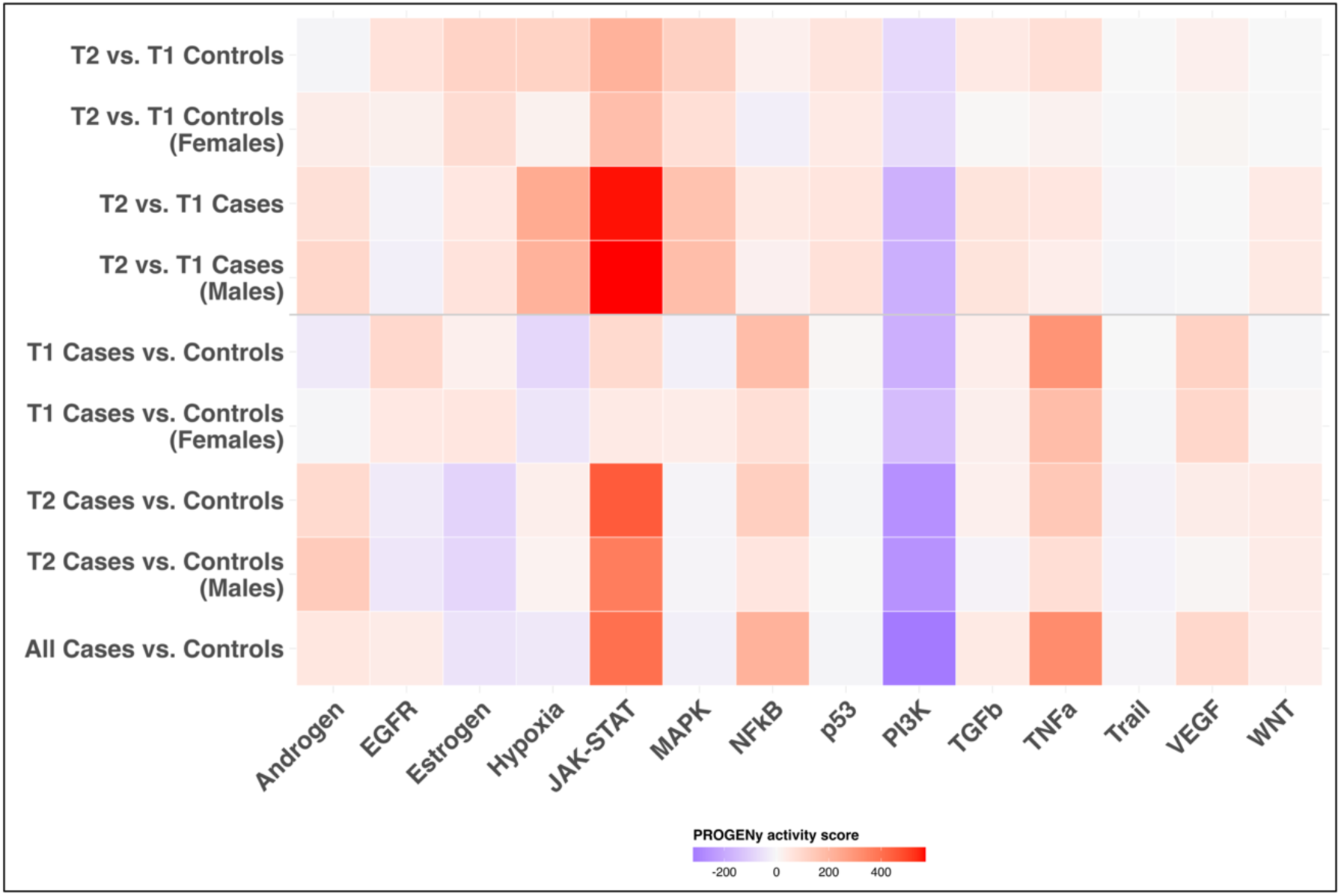
PROGENy pathway activity inference across fetal brain transcriptomes. Heat map shows inferred pathway activity scores for androgen, EGFR, estrogen, hypoxia, JAK-STAT, MAPK, NFκB, p53, PI3K, TGFβ, TNFα, Trail, VEGF, and WNT signaling. Warmer colors indicate increased inferred pathway activity, whereas cooler colors indicate decreased inferred pathway activity in the indicated comparison.

### Molecular effects on T1 fetal brains from prenatal cannabis exposure

T1 samples had insufficient male representation for statistically powered sex-stratified analysis; therefore, female-only and combined-cohort analyses were performed. Female-only analyses identified no significant DEGs and only a single DEP, NLRP2 (Fig. S15). In the combined sex cohort, approximately 30 genes reached statistical significance, including NLRC5, TNKS, PYCARD, SLC39A4, AGO4, and PBRM1 (Fig. S16), but these changes lacked coherent pathway organization and were not retained after sex stratification. Collectively, these findings indicate that prenatal cannabis exposure in T1 fetal brains did not induce a coordinated systems-level molecular response.

### Trajectory of gene and protein expression in fetal brain development from T1 to T2

#### In the absence of prenatal cannabis exposure (control brains)

Given the predominance of female samples in control T1 and T2 cohorts, analyses focused on this sex. At the transcriptomic level, female control brains exhibited a tightly coordinated developmental trajectory, with all significant genes upregulated in T2 relative to T1 (Fig. 8A). This program was characterized by increased expression of regulators of cortical specification and neuronal maturation (*25*), including TBR1 and EOMES (TBR2), key regulators of cortical neuron specification and laminar identity, marking progression from progenitor states toward differentiated neuronal populations. This development was accompanied by increased expression of SLC17A7 (VGLUT1), KCNJ4, and CHRM1, consistent with the emergence of glutamatergic neurotransmission, establishment of membrane excitability, and strengthening of cholinergic responsiveness. Upregulation of PPP1R1B further supports integration of synaptic input at the postsynaptic level, linking receptor activation to downstream signaling pathways and reinforcing functional circuit assembly. While most developmental transcriptomic changes were associated with forebrain specification and neuronal maturation programs, increased expression of SLN (sarcolipin) and NLRC5 suggests additional developmental remodeling beyond classical neuronal differentiation pathways.

**Figure 8.**
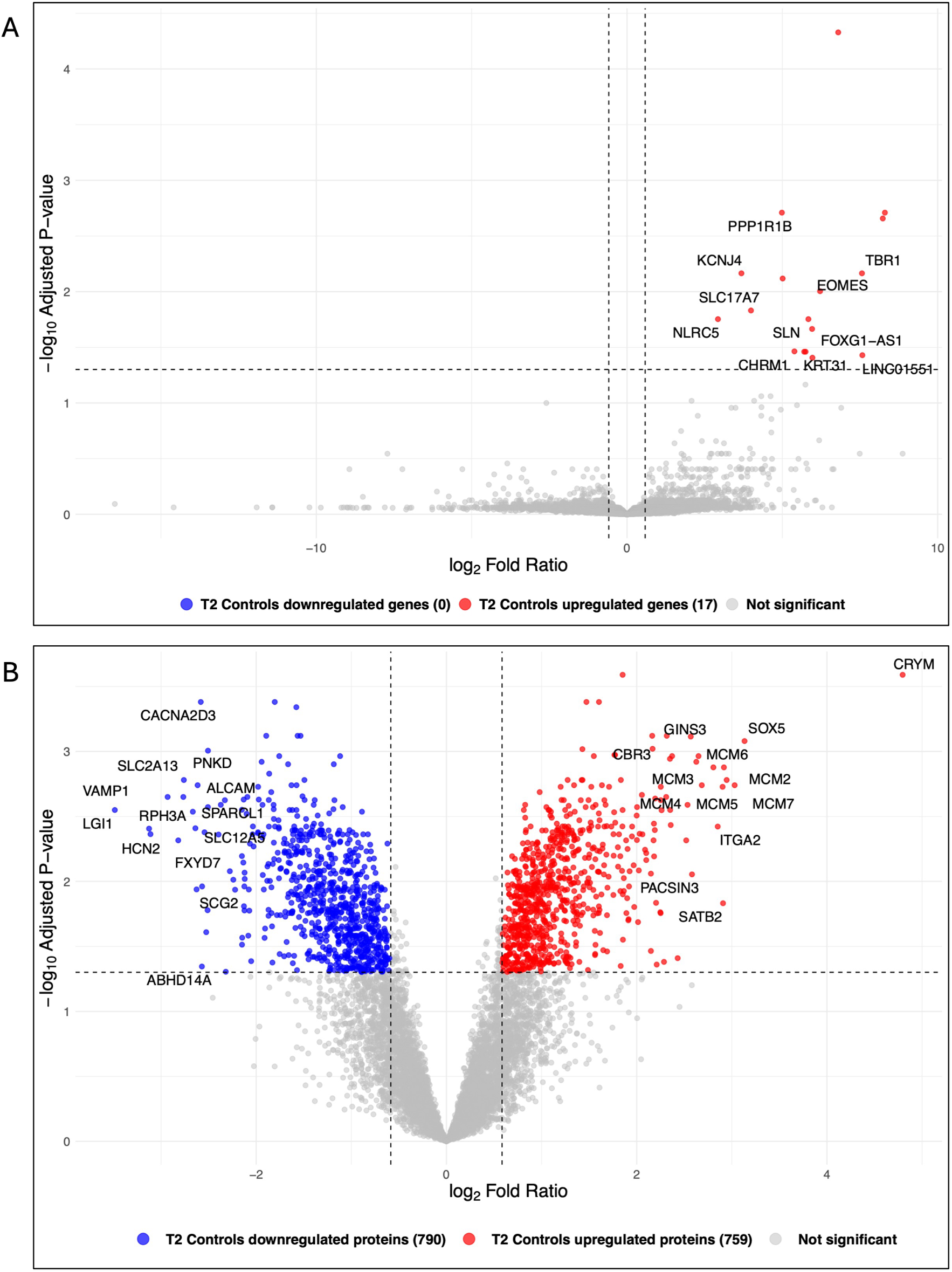
Sex-specific molecular trajectory of fetal brain development in female fetuses (T2 vs. T1 Controls). Transcriptomic **(A)** and proteomic **(B)** comparisons of T2 vs. T1 control brains showing gene products associated with development. Volcano plots showing quantified genes or proteins plotted by log₂ fold ratio and Benjamini-Hochberg adjusted P value. Dashed vertical lines indicate the fold-ratio threshold (|log₂FR| ≥ 0.58), and the horizontal dashed line indicates the adjusted P-value cutoff (adjusted P ≤ 0.05). In **A**, 17 DEGs were identified. Selected DEGs are labeled, and ENSG-only identifiers are omitted. All significant DEGs were upregulated in T2 relative to T1. In **B**, 1,549 DEPs were identified, with proteins both upregulated and downregulated in T2 relative to T1.

Consistent with this developmental program, Gene Ontology enrichment analysis identified processes related to cognition, learning and memory, ion transport, forebrain neuron differentiation, and cell fate specification (Fig. S17A), while KEGG analysis identified enrichment of the cholinergic synapse pathway (Fig. S17B). GSEA further revealed enrichment of translation-, RNA processing-, and cell cycle-associated pathways, together with relative depletion of neuronal system and oxidative phosphorylation pathways (Fig. S13).

At the proteomic level, female control samples showed a polarized developmental trajectory characterized by increased chromatin organization, transcriptional regulation, RNA processing, and cell cycle activity, alongside reduced neuronal functional systems (Fig. 8B). Proteins decreased in T2 included presynaptic vesicle and active-zone components (SNAP25, VAMP1/2, UNC13A/B, STX1B, RIMS1/2) together with calcium channel and postsynaptic signaling proteins (CACNA1A/B, CACNA2D1/2/3, CACNG2, GRIN1/2B), indicating reduced synaptic release and excitability-related functions (Fig. S18A, Table S2). Reduced abundance of the neurotransmitter transporters SLC17A6 (VGLUT2) and SLC32A1 (VGAT) further indicated remodeling of excitatory and inhibitory neurotransmission during fetal brain maturation (Fig. S18A; Fig. S23). In addition to neuronal and synaptic remodeling, developmental maturation was associated with a prominent DNA replication and genome maintenance module (Fig. S23), comprising MCM2–7, PCNA, RFC and RPA complexes, DNA polymerases, and DNA repair proteins, highlighting the extensive molecular restructuring accompanying human fetal brain development.

Consistent with this, Gene Ontology enrichment analysis identified synapse-related processes among downregulated proteins, including synaptic vesicle cycling, neurotransmitter transport, and regulation of synapse structure and activity, whereas upregulated proteins were enriched for RNA splicing, DNA replication, chromosome organization, and DNA repair (Fig. S18B-C). KEGG analysis similarly revealed depletion of neuronal signaling pathways, including synaptic vesicle cycling, neurotransmission, and axon guidance, alongside enrichment of chromatin- and cell-cycle-associated programs (Fig. S18D-E).

This coordinated reduction extended across neuronal systems governing adhesion, circuit specification, membrane transport, and mitochondrial metabolism (Fig. S23).

Downregulated proteins included solute carrier transporters, oxidative phosphorylation components, mitochondrial carriers, and V-ATPase complexes, indicating reduced transport and bioenergetic capacity. Proteomic interaction analysis linked these changes to synaptic vesicle fusion, presynaptic exocytosis, transsynaptic adhesion pathways including Ephrin–EPH receptor signaling (Fig. S23), SLIT–ROBO-mediated axon guidance (Fig. S18, Fig. S23), and ion channel-mediated excitability, whereas upregulated modules were associated with chromatin remodeling, transcriptional regulation, cell cycle progression, and RNA processing (Fig. S23).

#### In the presence of prenatal cannabis exposure (case brains)

Due to the predominance of male subjects in this category, analyses focused on this sex. Comparison of T1 to T2 male case samples revealed a strong molecular signal, with ∼670 DEGs and ∼2,400 DEPs (Fig. S19A–B). Transcriptomic analysis showed upregulation of neurodevelopmental regulators including TBR1, EOMES, FOXG1, and NEUROG1, consistent with progression of cortical maturation and neuronal differentiation. These changes largely overlapped the developmental trajectory observed in female controls and followed a shared directional axis of change at both the transcriptomic and proteomic levels (Figs. S20, S21A–B).

Transcriptomic enrichment identified reduced synaptic and phospholipid-associated pathways together with activation of DNA replication, cell cycle, and DNA repair programs (Fig. S22A–E). Proteomic enrichment similarly revealed suppression of synaptic, neurotransmission, endocannabinoid, and oxidative phosphorylation pathways alongside increased RNA processing, chromatin remodeling, DNA replication, and cell cycle activity (Fig. S24A–F). These pathway-level changes closely mirrored those observed in the T2 male cannabis exposure comparison.

## Discussion

This study provides for the first time a multi-omics analysis of the effects of prenatal cannabis exposure in the developing human fetal brain and links maternal cannabis use to molecular alterations associated with fetal neurodevelopment. Several study design features strengthened attribution of our observed changes to prenatal cannabis exposure. Integrated transcriptomic and proteomic profiling provided complementary, non-subjective, measures of molecular perturbation in cannabis-exposed fetal brains. Matched control cohorts enabled direct comparison with non-exposed fetal brains, while urine toxicology screening validated cannabis exposure in cases, the lack thereof in controls and excluded additional drugs of abuse in both. Inclusion of both T1 and T2 fetal brains enabled assessment of neurodevelopment in the absence and presence of prenatal cannabis exposure.

The ECS is a central regulator of fetal brain development, controlling neuronal generation, migration, axonal guidance, and synaptic assembly required for functional circuit formation through CB1 receptor signaling. These processes are tightly regulated in space and time during cortical development (*11, 12*). CB1 receptor is detectable in the human fetal brain by 19 weeks of gestation. Plus, spiral arteries supplying maternal blood to the placenta do not fully open until approximately 11–12 weeks of gestation (*26*). For these reasons, it is not surprising that fetal cannabis exposure and associated alterations in maternal–fetal blood flow are more prominent during T2 (*27, 28*). In accord with these observations, our data suggest increased vulnerability to prenatal cannabis exposure during T2, a period characterized by elevated metabolic demand and cortical maturation. Likewise, CB1-deficient mice exhibit abnormalities in cortical wiring and seizure susceptibility and are resistant to prenatal Δ^9^-THC-induced developmental alterations (*29*). DAGLα-deficient mice show loss of retrograde endocannabinoid signaling, impaired hippocampal synaptic plasticity, and reduced neurogenesis. These animal studies demonstrate that disruption of endocannabinoid signaling impairs neuronal differentiation.

Consistent with the above observations in animals, prenatal cannabis exposure in T2 male fetal brains induced coordinated systems-level reprogramming characterized by reduced synaptic and adhesion systems, disrupted membrane transport, metabolic homeostasis, and mitochondrial energy production, despite continued activation of developmental regulatory programs. These findings indicate that following prenatal cannabis exposure, developmental signaling persists while neuronal and metabolic functional systems are progressively compromised (Figs. 3-7 and S8).

In addition, in the developing T2 male cannabis-exposed brains, coordinated reduction of synaptic scaffold, vesicle-cycle, and protocadherin-mediated adhesion systems indicates impaired synapse formation, neurotransmission, and circuit organization (Figs. 3 and 4). Disruption of NRXN1-associated synaptic organization, SNARE-dependent neurotransmitter release machinery, and protocadherin-dependent neuronal identity systems have been broadly implicated in neurodevelopmental disorders, including autism spectrum disorder, schizophrenia, epilepsy, and other disorders of neuronal connectivity (*18, 20, 30*). Protocadherins, including PCDH10, play important roles in neuronal circuit formation, and prenatal Δ^9^-THC exposure in rodents disrupt neuronal differentiation and axonal development through CB1-dependent mechanisms (*31, 32*). PCDH10 has also been implicated in autism spectrum disorder through inherited genomic alterations identified in human genetic studies (*33*).

We also identified broad transporter dysregulation as a prominent feature of the cannabis-exposed fetal brain (Fig. 6 and S8), affecting neurotransmission, ion homeostasis, nutrient transport, and metabolic support. Downregulated transporters identified in T2 cannabis-exposed males, including SLC6A1 (GAT1), SLC32A1 (VGAT), SLC1A1 (EAAT3), SLC9A6 (NHE6), and SLC9A7 (NHE7), have been linked to epilepsy, autism spectrum disorder, intellectual disability, schizophrenia, and other neurodevelopmental disorders (*34*).

Mitochondrial dysfunction has been linked to multiple neurodevelopmental disorders (*35*). Consistent with this, cannabis-exposed T2 male fetal brains (Fig. 5) showed coordinated reduction of oxidative phosphorylation proteins, mitochondrial SLC25 transporters, and regulators of mitochondrial dynamics and quality control, including

OPA1, MFF, and PARL, indicating widespread impairment of mitochondrial function. Consistent with this, Δ^9^-THC exposure of human iPSC-derived neurons disrupts mitochondrial and oxidative phosphorylation pathways, indicating a conserved mitochondrial response to cannabinoid exposure (*22*). Likewise, prenatal Δ^9^-THC exposure reduces mitochondrial respiration in animal models (*36*).

Pathway-level analyses identified activation of JAK-STAT, TNFα, and NFκB signaling together with suppression of PI3K-associated trophic signaling (Fig. 7). JAK-STAT, TNFα, and NFκB pathways regulate immune responses and contribute to neural development and neuronal differentiation. PI3K-AKT-mTOR signaling is a central regulator of neuronal development, synaptogenesis, corticogenesis, cell migration, and axon guidance (*37*). In parallel, numerous transcription factors and zinc finger proteins were concordantly upregulated in cannabis-exposed T2 male brains (Fig. 3), including FOXG1 and SOX5. The latter are key regulators of forebrain development and neurogenesis, the disruption of which causes severe neurodevelopmental disorders including FOXG1 syndrome and Lamb–Shaffer syndrome (*38, 39*). Collectively, these findings indicate that prenatal cannabis exposure disrupts the balance between developmental, trophic, and inflammatory signaling during T2 male brain development.

Prenatal cannabis exposure did not produce a coordinated systems-level response in T1 female brains (Figs. S15 and S16). This may reflect lower sensitivity to cannabinoid perturbation during early gestation, as CB1 receptor expression and endocannabinoid signaling become increasingly integrated into neuronal differentiation, axon guidance, synaptogenesis, and circuit refinement during mid-gestational cortical development (*9, 10, 12, 40*). Δ^9^-THC was detectable but variable in a subset of T1 fetal brain samples (*15*), potentially reflecting limited early fetal exposure prior to complete opening of maternal spiral arteries at the end of the first trimester (*26*). Consistent with this, the strongest molecular effects of prenatal cannabis exposure were observed in our T2 cohort.

Female control T1-to-T2 maturation was characterized by coordinated remodeling of synaptic, transport, mitochondrial, and metabolic systems. Although transcriptomic analysis identified only a limited number of developmental changes (Fig. 8A), high-depth DIA proteomics revealed extensive developmental remodeling during the female T1-to-T2 transition, substantially expanding upon the developmental features previously resolved (as described below) by transcriptomic and lower-depth proteomic studies of the human fetal brain (Fig. 8B).

While developmental maturation of neuronal pathways has been described previously using transcriptomics (*41, 42*) and proteomics (*43, 44*), our proteomic analysis identified broader restructuring of membrane transport, mitochondrial carrier, V-ATPase, and metabolic systems (Fig. S23). Beyond established markers of cortical neuronal maturation, developmental transcriptomic changes included increased expression of NLRC5, a regulator of MHC class I gene expression, and SLN (Fig. 8A). Proteomic remodeling likewise encompassed a DNA replication and genome maintenance module containing MCM2–7, PCNA, RFC/RPA complexes, DNA polymerases, and DNA repair proteins (Fig. S23). Together, these findings broaden the developmental features identified during the T1-to-T2 transition beyond neuronal differentiation and synaptic maturation alone. Importantly, many of these same systems showed reduced abundance under prenatal cannabis exposure, suggesting that cannabinoid exposure may affect developmental remodeling of neuronal transport, metabolic support, and functional maturation.

Our study has several limitations, including limited male and female representation in T1 and T2, respectively, which constrained interpretation of sex-specific effects of prenatal cannabis exposure and fetal brain development. Our cannabis-exposed T1 and T2 cohorts were separated by ∼33 days, constraining our ability to map fetal brain development and the effect of cannabis exposure over a longer period. The small size in each cohort also limited the power of our analyses. Since we could enroll subjects only when they showed up in the clinic, cannabis exposure history relied on self-reports, preventing objective data on the dose of cannabis consumed and therefore assessment of dose-response relationships. Although urine toxicology distinguished cannabis users from non-users, prior cannabis use among controls and use of other substances among controls or cases cannot be excluded. While alcohol and nicotine exposure (Table S1) may contribute to the observed phenotype, these substances act through distinct developmental mechanisms (*45, 46*), whereas the molecular alterations identified here are most consistent with disruption of the endocannabinoid signaling. Although Δ^9^-THC and 11-OH-THC are the most likely mediators of the observed effects, contributions from other cannabis constituents cannot be excluded. Finally, limited tissue availability restricted analyses to whole-brain, and future studies examining specific brain regions may provide additional insight into region-specific effects of prenatal cannabis exposure.

In summary, this integrated multi-omics analysis provides for the first time direct molecular evidence that prenatal cannabis exposure is associated with disruption of synaptic, metabolic, and developmental systems in the T2 human fetal brain. Convergent transcriptomic, proteomic, and pathway-level alterations indicate that prenatal cannabis exposure during T2 shifts the developing male brain toward reduced synaptic capacity, impaired metabolic support, and increased cellular stress. These findings provide a plausible molecular basis for neurodevelopmental and behavioral deficits reported in children exposed to cannabis *in utero* (*47, 48*). Our findings raise significant concerns regarding prenatal cannabis exposure and identify molecular pathways and developmental regulators that warrant mechanistic and translational follow-up. Future studies can build upon our data to explore how these pathways contribute to adverse effects of prenatal cannabis exposure on human fetal neurodevelopment.

## Materials and Methods

### Study design

Research Objectives: This cross-sectional observational study used integrated transcriptomics and DIA proteomics to characterize molecular effects of prenatal cannabis exposure on the developing T1 and T2 human fetal brains.

<u>Research subjects</u>: This study was approved by the institutional review board of the University of Washington. All potential study participants received pregnancy options counseling and made an independent decision with their healthcare provider to terminate their pregnancy via a surgical procedure. The procedure used was either dilation and curettage in T1 or dilation and evacuation (usually a two-day procedure) in T2. Once consent for the pregnancy termination procedure was signed, and subjects had their routine dating ultrasound, they were screened for study eligibility. Screening for eligibility and subsequent consent for this research study was obtained by personnel not involved in consenting subjects for termination, providing termination care, or counseling.

Eligibility criteria included subjects age ≥18 years, seeking pregnancy termination between 8 and 24 weeks of gestation (by ultrasound), proficiency in reading and speaking English, use of cannabis by inhalation, ingestion or both during pregnancy, including within 2 days of termination and no use of other drugs of abuse during pregnancy (see below). Henceforth, these subjects are referred to as cases. Control subjects were eligible for the study if they met the above criteria and had not used cannabis or other drugs of abuse during pregnancy. If eligible, subjects consented to the collection of fetal tissue for research and were asked to complete an enrollment survey which included demographic information, history of their reproductive health, as well as substance and cannabis use during pregnancy. Following completion of the survey, the subjects were asked to donate a urine sample to conduct a toxicology screen to confirm cannabis use (cases) or lack thereof (controls), as well as lack of use of drugs of abuse (all subjects). All enrolled subjects received a $40 gift card as remuneration for completing the survey.

Gestational age of the fetus was classified based on fetal foot length and expressed as post-conception age. Trimester 1 (T1) was defined as ≤84 days post-conception and Trimester 2 (T2) as >84 days. The collected fetal tissues (including the brain) were stored at 4°C prior to transportation (on the same day) to the University of Washington where they were separated by dissection into individual tissues, flash-frozen at −80 °C and stored at −80 °C until processing. A subset of samples was stabilized overnight in RNAlater at 4 °C prior to freezing at −80 °C. Due to the small size of the fetal brain, regional dissection was not performed, and RNA and protein were extracted from whole tissue (Fig. 1 and Table S1).

<u>Experimental design:</u> Because this was an exploratory first-in-kind human fetal brain multi-omics study, no prospective power calculation, interim analyses, or stopping rules were defined. Subjects were recruited with a goal to attain about the same number of controls and cases (irrespective of fetal sex) over the duration of funding. Recruitment was completed in July 2024.

The primary endpoints were to compare the following transcriptomic and proteomic changes in the fetal brains associated with prenatal cannabis exposure: T1 cases vs. controls, T2 cases vs. controls, pooled cases vs. controls. Secondary endpoints were to determine transcriptomic and proteomic changes due to fetal brain development, i.e., T1 controls vs. T2 controls, and T2 cases vs. T1 cases.

<u>Data inclusion/exclusion</u>: Samples were used for transcriptomics or proteomics analysis only if the urine toxicology test confirmed the self-reported information. Urine toxicology tests screened for cannabinoids, alcohol, acetaminophen, amphetamines or methamphetamines, barbiturates, benzodiazepines, cocaine, fentanyl, methadone, opiates, phencyclidine, and tricyclic antidepressants. Except for the following, detection of any of these compounds was exclusionary: a) detection of benzodiazepines, fentanyl, or opiates was permitted when consistent with medications administered as part of the clinical procedure (dilation and evacuation); b) use of alcohol, acetaminophen, tobacco (not tested), and tricyclic antidepressants was not exclusionary. Samples failing quality-control criteria for RNA quality (RIN number < 4), and where there was insufficient volume (some tissues were used up for cannabinoid quantification) were excluded.

Replicates: Technical replicates of RNA sequencing or proteomics (LC-MS) analyses were not performed.

<u>Transcriptomics:</u> Total RNA was extracted from 30–50 mg of frozen fetal brain tissue (n=34) using the PureLink™ RNA Mini Kit (Invitrogen™, Carlsbad, CA, USA) according to the manufacturer’s instructions. Tissue was homogenized in lysis buffer using Combitips™ Advanced Tips (Eppendorf, Enfield, CT, USA), RNA was purified using silica spin columns with sequential wash steps, and RNA was eluted in RNase-free water. Purified RNA was stored at −80°C until downstream processing.

Library preparation and sequencing: RNA integrity was assessed by Novogene (Sacramento, CA, USA), and samples with an RNA integrity number (RIN) of at least 4 (n=34) were retained for transcriptomic analysis (Fig. 1, Table S1). Poly(A) mRNA was enriched using poly-T oligo-attached magnetic beads, followed by fragmentation and cDNA synthesis. Libraries were constructed as non-directional and sequenced in paired-end mode with 150-bp reads, generating FASTQ files.

Read alignment and quantification: RNA-seq reads were aligned to the *Homo sapiens* reference genome (GRCh38) using the HISAT2 aligner. Gene-level read counts were generated from aligned BAM files using featureCounts from the Bioconductor Rsubread package with Ensembl gene annotations. Count matrices were imported into R for downstream transcriptomic analyses.

Gene filtering: Prior to differential expression analysis, genes with low expression were filtered to reduce noise. Genes with at least 15 total counts across all samples and 10 counts in the smallest comparison group were retained. From 60,173 genes, a total of 23,206 genes passed filtering.

<u>Proteomics:</u> Tissue pieces (11-428 mg) were washed with Cell Wash Buffer and homogenized in 0.6 mL Thermo Scientific™ N-PER™ Neuronal Protein Extraction Reagent (Catalog number 87792) supplemented with Thermo Scientific™ 1X Halt™ Protease and Phosphatase Inhibitor (Catalog number 78440). Homogenization was performed using ceramic beads with four 15-second cycles, with 30-second incubations on ice between cycles. Homogenates were agitated (350 rpm, 10 minutes) and debris pelleted by centrifugation (3,000 × g, 15 minutes, 4 °C). Protein concentration in the tissue lysate was determined using a bicinchoninic acid (BCA) assay. Lysate was diluted to 1 mg/mL in extraction reagent supplemented with protease inhibitors.

Fifty µg of lysate protein was adjusted to 200 µL with LC-MS–grade water and subjected to chloroform–methanol precipitation by sequential addition of methanol (800 µL), chloroform (200 µL), and water (600 µL) with vortexing after each addition (*49*). Phase separation was induced by centrifugation at 19,090 × g for 1 min. The aqueous phase was discarded. Methanol (600 µL) was added, and proteins were pelleted by centrifugation (19,090 × g, 5 min). The air-dried protein pellet was resuspended in 100 mM ammonium bicarbonate (50 µL), and 20 µg protein was used for digestion. Proteins were reduced with 5 mM tris (2-carboxyethyl) phosphine (TCEP; Thermo Fisher Scientific, Rockford, IL, USA) for 25 min at room temperature, alkylated with 14 mM iodoacetamide (Thermo Fisher Scientific, Rockford, IL, USA) for 30 min in the dark, quenched with 10 mM

TCEP for 15 min, and digested overnight with trypsin (Thermo Fisher Scientific, Rockford, IL, USA) at 37 °C. Digestion was quenched with formic acid (pH <3). Following centrifugation (25,000 × g, 10 min, 4 °C), the peptide-containing supernatant was transferred to autosampler vials for LC–MS analysis.

LC–MS data acquisition: Analytical columns were prepared in-house from fused silica capillary tubing (75 µm ID × 363 µm OD). Capillaries were pulled using a P-2000 laser puller (Sutter Instruments, CA, USA). Columns were slurry-packed with Reprosil-Pur 120 C18-AQ resin (5 µm) to a length of 15 cm and coupled to a Vanquish Neo UHPLC system for LC–MS analysis (Thermo Fisher Scientific, Waltham, MA, USA).

Peptide samples were analyzed by nano LC-MS/MS on a Vanquish Neo UHPLC system coupled to an Orbitrap Astral mass spectrometer (Thermo Fisher Scientific, Waltham, MA, USA). Peptides were loaded in trap-and-elute mode onto a 15 cm × 75 µm analytical column at 0.4 µL/min. Mobile phases consisted of water (A) and 80% acetonitrile (B). Formic acid (0.1%, v/v) was added to both mobile phases as a pH modifier. Peptides were separated using a 30-min gradient from 1% to 45% B followed by wash and re-equilibration (total run time, 36 min). Data were acquired in positive-ion DIA mode. The DIA acquisition strategy was adapted from Guzman et al. (2024) (*50*). Full MS1 scans were acquired in the Orbitrap over m/z 380–980 at a resolution of 240,000 with an AGC target of 5 × 10⁵ and a maximum injection time of 3 milliseconds. DIA MS2 scans were acquired on the Astral detector using sequential non-overlapping 2 m/z isolation windows across m/z 380–980. Fragment ions were generated by higher-energy collisional dissociation (HCD; 30% normalized collision energy) and detected over m/z 200–1500 with an AGC target of 2 × 10⁴ and a maximum injection time of 2.5 milliseconds.

Raw DIA files were processed in DIA-NN (v2.2.0) using a two-pass library-assisted workflow with a predicted *Homo sapiens* UniProtKB spectral library. MS1 mass accuracy was set to 5 ppm, and MS2 mass accuracy was set to 10 ppm. Searches used trypsin specificity (≤2 missed cleavages), peptide lengths of 7–30 amino acids, precursor charge states of 2–4, fixed carbamidomethylation, and variable methionine oxidation.

Protein inference and quantification were performed at the gene level using high-precision mode with interference correction and match-between-runs. Identifications were filtered at 1% FDR at the precursor and protein-group levels.

### Statistical Analysis

<u>Differential expression analysis of the RNAseq data:</u> Differential expression was performed using limma-voom. Counts were transformed to log2 counts per million (logCPM), and weighted linear models were fitted using voom-derived observational weights to account for the mean–variance relationship. The design matrix included cannabis exposure and trimester as primary factors, with fetal sex and gestational age as covariates. Data quality and global structure were assessed using multidimensional scaling (MDS). Sample-specific weights were applied to down-weight poorly fitting samples rather than exclude them. Statistical significance was assessed using empirical Bayes moderation with Benjamini–Hochberg correction. Genes were designated as differentially expressed genes (DEGs) at an adjusted P value ≤0.05. DEGs with an absolute log_2_ fold ratio (|log_2_ FR|) ≥ 0.58, corresponding to a 1.5-fold difference in either direction, were considered biologically significant.

<u>Protein filtering and normalization:</u> Protein-level quantitative matrices exported from DIA-NN were analyzed in R using Bioconductor. Proteins passing DIA-NN false discovery rate filtering (1% protein-group level) and supported by at least two proteotypic peptides were retained. Data were log₂-transformed, and technical variability was assessed using pooled quality-control runs and exploratory diagnostics. Scale normalization was applied to remove residual technical effects while preserving biological variability. Data completeness was high, consistent with narrow-window DIA acquisition, and missing values were primarily limited to low-abundance proteins.

Proteins with >20% missing values were excluded. For sensitivity analyses, remaining missing values were imputed using the Random Forest–based missForest approach and compared with complete-case analyses to assess robustness. Differential protein abundance was assessed using limma-trend with empirical Bayes moderation. Models included gestational age and fetal sex as covariates, and sample-specific weights were estimated to down-weight outlier samples and account for heteroscedasticity. Statistical significance was assessed using Benjamini–Hochberg false discovery rate control.

Proteins were designated as differentially abundant proteins (DEPs) at an adjusted P value ≤0.05. DEPs with an absolute log_2_ fold ratio (|log_2_ FR|) ≥0.58, corresponding to a 1.5-fold difference in either direction, were considered biologically significant.

Complete-case and imputed analyses yielded concordant results, and primary biological conclusions were unaffected by imputation.

<u>Computational environment and reproducibility</u>: All statistical analyses were conducted in R (version 4.5.2) using Bioconductor packages. Gene set enrichment analysis (GSEA) was performed using fgsea (fgseaMultilevel) on pre-ranked gene lists, with genes ranked by signed −log10 (P value) and direction of change (log_2_FR). Gene sets were obtained from the MSigDB Hallmark (H) and Reactome (C2:CP:REACTOME) collections.

Enrichment scores were normalized for gene set size, and significance was assessed using adaptive multilevel split Monte Carlo estimation with Benjamini–Hochberg correction. Pathways with adjusted P < 0.10 were considered enriched and visualized using normalized enrichment scores (NES) with significance encoded as −log_10_ adjusted P values. Network analyses were performed using STRINGdb and Cytoscape.

## Funding

This study was funded by NIH P01 DA032507 (to JDU). IAG was supported by a National Institute for Childhood Health and Disease Award R24HD000836.

## Author Contributions

J.D.U. designed and supervised the research. L.S.B., J.C.D. and I.A.G. provided resources to collect the samples and/or collected samples. X.C. performed RNA isolation and RNA-seq sample preparation. D.K.S. and B.P. contributed proteomics methodology and performed protein extraction. G.K. performed proteomic sample preparation and DIA-MS data acquisition. T.K.B. and J.W.M. performed bioinformatic and statistical analyses. G.K. analyzed and integrated the transcriptomic and proteomic data, conducted pathway analyses, prepared figures and tables, and wrote the manuscript. All authors interpreted the data, revised the manuscript, and approved the final version.

## Competing Interest

J.D.U. has consulted and continues to consult for numerous pharmaceutical companies, none of which are associated with sale of cannabis products. He also owns stocks in Precision Quantomics Inc., a company focused on providing proteomics services. Bhagwat Prasad is co-founder of Precision Quantomics Inc. and recipient of research funding from AbbVie, Boehringer Ingelheim, Bristol-Myers Squibb, Genentech, Generation Bio, Gilead Sciences, Johnson & Johnson, Merck, Novartis, and Takeda.

## Data, code and material availability

All data associated with this study are included in the paper or the Supplementary Materials. RNA-seq data are deposited in GEO/SRA under accession GSE336661, and proteomics data are deposited in the PRIDE repository under accession PXD080835. The datasets will be made publicly available upon publication.

## Acknowledgments

This work was supported by National Institutes of Health [Grant P01 DA032507, J.D.U.; Grant R24HD000836, IAG]. We thank Edward J. Kelly (University of Washington) for scientific discussions and interpretation of this work at earlier stages of this research and Lucinda A. Cort (University of Colorado School of Medicine) for her contribution to fetal tissue collection. We thank Aditya R. Kumar (University of Washington) for help with collection of samples.

## Notes

### Competing Interest Statement

The authors have declared no competing interest.

