## Supplementary Figures and Tables for "Prenatal cannabis exposure affects human fetal neurodevelopment: an integrated multi-omics study"

### Supplemental table 1. Demographic and cannabis exposure characteristics of the fetal brain cohorts

| Subject ID | Trimester | Postconception age (weeks) | Fetal sex | Assay modality | Maternal race | Maternal ethnicity | Group | Cannabis use route | Cannabis frequency | Amount at last sitting | Tobacco use last month |
|---|---|---|---|---|---|---|---|---|---|---|---|
| 29111 | 1 | 7.6 | F | Transcriptomic / Proteomic | Black or African American | Non-Hispanic | Control | | | | NO |
| 90001 | 1 | 10.3 | F | Transcriptomic / Proteomic | White | Non-Hispanic | Control | | | | NO |
| 90007 | 1 | 10.9 | M | Transcriptomic / Proteomic | NA† | Hispanic | Control | | | | NO |
| 90009 | 1 | 8.1 | F | Transcriptomic / Proteomic | Black or African American | Non-Hispanic | Control | | | | NO |
| 90028 | 1 | 7.6 | F | Transcriptomic / Proteomic | Asian | Non-Hispanic | Control | | | | NO |
| 90034 | 1 | 10.3 | F | Transcriptomic / Proteomic | White | Non-Hispanic | Control | | | | NO |
| 28711 | 1 | 11.4 | M | Transcriptomic / Proteomic | White | Non-Hispanic | Case | inhalation / ingestion | 1-2 times per week | 0.3-0.5 grams/marijuana plant | NO |
| 28928 | 1 | 9.6 | M | Transcriptomic / Proteomic | Black or African American | Non-Hispanic | Case | inhalation | 2-3 times per month | 3 grams marijuana plant | YES |
| 28972 | 1 | 10.3 | M | Proteomic | White | Non-Hispanic | Case | inhalation / ingestion | three or more times per day | 1/2 gram marijuana plant | YES |
| 29045 | 1 | 10.6 | F | Proteomic | White | Non-Hispanic | Case | inhalation | three or more times per day | 0.10 grams marijuana plant | NO |
| 29050 | 1 | 8.1 | M | Proteomic | Black or African American | Non-Hispanic | Case | inhalation | three or more times per day | 1 gram marijuana plant | NO |
| 29110 | 1 | 11.7 | F | Transcriptomic | NA | Hispanic | Case | inhalation / ingestion | twice per day | 2 grams marijuana plant | NO |
| 29141 | 1 | 9.6 | M | Transcriptomic / Proteomic | Asian | Non-Hispanic | Case | inhalation | three or more times per day | 1 gram marijuana plant | NO |
| 90004 | 1 | 10.6 | F | Transcriptomic / Proteomic | White | Non-Hispanic | Case | inhalation / ingestion | three or more times per day | 1 gram marijuana plant | NO |
| 90006 | 1 | 8.4 | F | Transcriptomic / Proteomic | White | Non-Hispanic | Case | ingestion | once per day | 100 mg THC | YES |
| 90020 | 1 | 10.6 | F | Transcriptomic / Proteomic | White | Non-Hispanic | Case | inhalation / ingestion | three or more times per day | 2 mg marijuana plant/100 mg THC | NO |
| 90024 | 1 | 10.4 | M | Transcriptomic / Proteomic | White | Non-Hispanic | Case | inhalation | twice per day | 3 grams marijuana plant | NO |
| 90025 | 1 | 8.1 | M | Transcriptomic | Asian | Non-Hispanic | Case | NR‡ | NR | NR | NR |
| 90030 | 1 | 11.1 | M | Transcriptomic / Proteomic | Black or African American | Non-Hispanic | Case | inhalation | three or more times per day | 2 grams marijuana plant | YES |
| 90032 | 1 | 7.7 | F | Transcriptomic | Asian | Non-Hispanic | Case | inhalation | three or more times per day | not sure | YES |
| 90033 | 1 | 8.9 | F | Transcriptomic | American Indian / Alaska Native | Hispanic | Case | inhalation | 1-2 times per week | 0.5 grams | NO |
| 90035 | 1 | 7.6 | F | Transcriptomic / Proteomic | NA | Hispanic | Case | NR | NR | NR | NR |
| 28726 | 2 | 17.9 | F | Transcriptomic / Proteomic | Native Hawaiian / Pacific Islander | Non-Hispanic | Control | | | | NO |
| 28973 | 2 | 18.9 | F | Transcriptomic / Proteomic | Asian | Non-Hispanic | Control | | | | NO |
| 29035 | 2 | 13.7 | F | Transcriptomic / Proteomic | Asian | Non-Hispanic | Control | | | | NO |
| 29058 | 2 | 12.7 | M | Proteomic | NA | Hispanic | Control | | | | NO |
| 90002 | 2 | 13.0 | F | Transcriptomic / Proteomic | White | Non-Hispanic | Control | | | | NO |
| 90013 | 2 | 12.3 | M | Transcriptomic / Proteomic | NA | Hispanic | Control | | | | NO |
| 90019 | 2 | 14.0 | F | Transcriptomic / Proteomic | White | Non-Hispanic | Control | | | | YES |
| 90031 | 2 | 14.0 | M | Transcriptomic / Proteomic | Black or African American | Non-Hispanic | Control | | | | NO |
| 90037 | 2 | 13.4 | M | Transcriptomic / Proteomic | NA | Hispanic | Control | | | | NO |
| 28891 | 2 | 14.0 | M | Transcriptomic | Black or African American | Non-Hispanic | Case | inhalation / other | once per month or less | NR | YES |
| 28894 | 2 | 12.4 | M | Proteomic | White | Non-Hispanic | Case | inhalation | three or more times per day | 1-2 grams plant | YES |
| 28983 | 2 | 15.4 | M | Proteomic | White | Non-Hispanic | Case | inhalation / other | three or more times per day | NR | YES |
| 28998 | 2 | 13.0 | M | Proteomic | Black or African American | Non-Hispanic | Case | inhalation | three or more times per day | 1 gram marijuana plant | NO |
| 29069 | 2 | 15.7 | M | Transcriptomic | Black or African American | Non-Hispanic | Case | inhalation | once per day | 0.5 grams/plant | NO |
| 90000 | 2 | 14.7 | F | Proteomic | White | Non-Hispanic | Case | inhalation / ingestion | twice per day | 1 gram marijuana plant | NO |
| 90014 | 2 | 15.0 | M | Transcriptomic / Proteomic | White | Hispanic | Case | inhalation | three or more times per day | not sure | NO |
| 90016 | 2 | 14.4 | M | Transcriptomic / Proteomic | Native Hawaiian / Pacific Islander | Non-Hispanic | Case | inhalation | twice per day | NR | NO |
| 90018 | 2 | 14.7 | M | Transcriptomic / Proteomic | White | Non-Hispanic | Case | inhalation | three or more times per day | 0.5 mg THC oil | NO |
| 90022 | 2 | 15.7 | M | Proteomic | White | Non-Hispanic | Case | inhalation | three or more times per day | 3 grams marijuana plant | YES |
| 90023 | 2 | 12.4 | M | Transcriptomic / Proteomic | Black or African American | Non-Hispanic | Case | inhalation | 2-3 times per month | 0.5 marijuana plant | NO |
| 90038 | 2 | 13.0 | M | Transcriptomic | NA | Hispanic | Case | inhalation | three or more times per day | NR | YES |

† NA, not available. ‡ NR, not reported. THC-equivalent dose was estimated assuming 15% Δ9-THC by weight in cannabis plant material. Based on self-reported amount consumed at last sitting, the estimated THC-equivalent dose was 183 ± 145 mg per use in T1 participants (n = 13 of 15) and 161 ± 138 mg per use in T2 participants (n = 7 of 12).

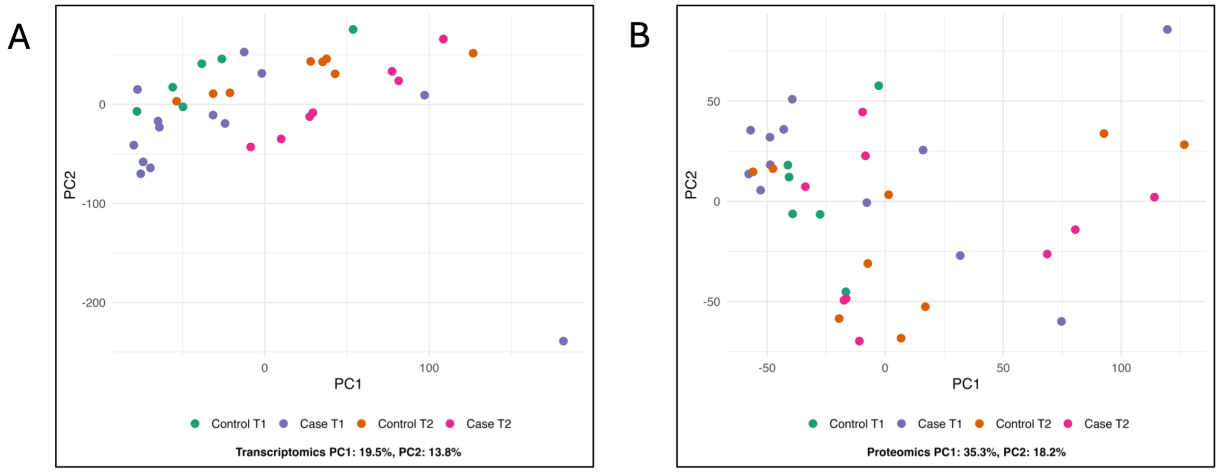


**Figure S1. Principal component analysis of fetal brain transcriptomic and proteomic data.** (**A**) Normalized transcriptomic data segregated primarily by trimester, with PC1 and PC2 explaining 19.5% and 13.8% of the variance, respectively. (**B**) Normalized proteomic data showed substantial overlap between trimesters though it remained a major source of variation, with PC1 and PC2 explaining 35.3% and 18.2% of the variance, respectively.

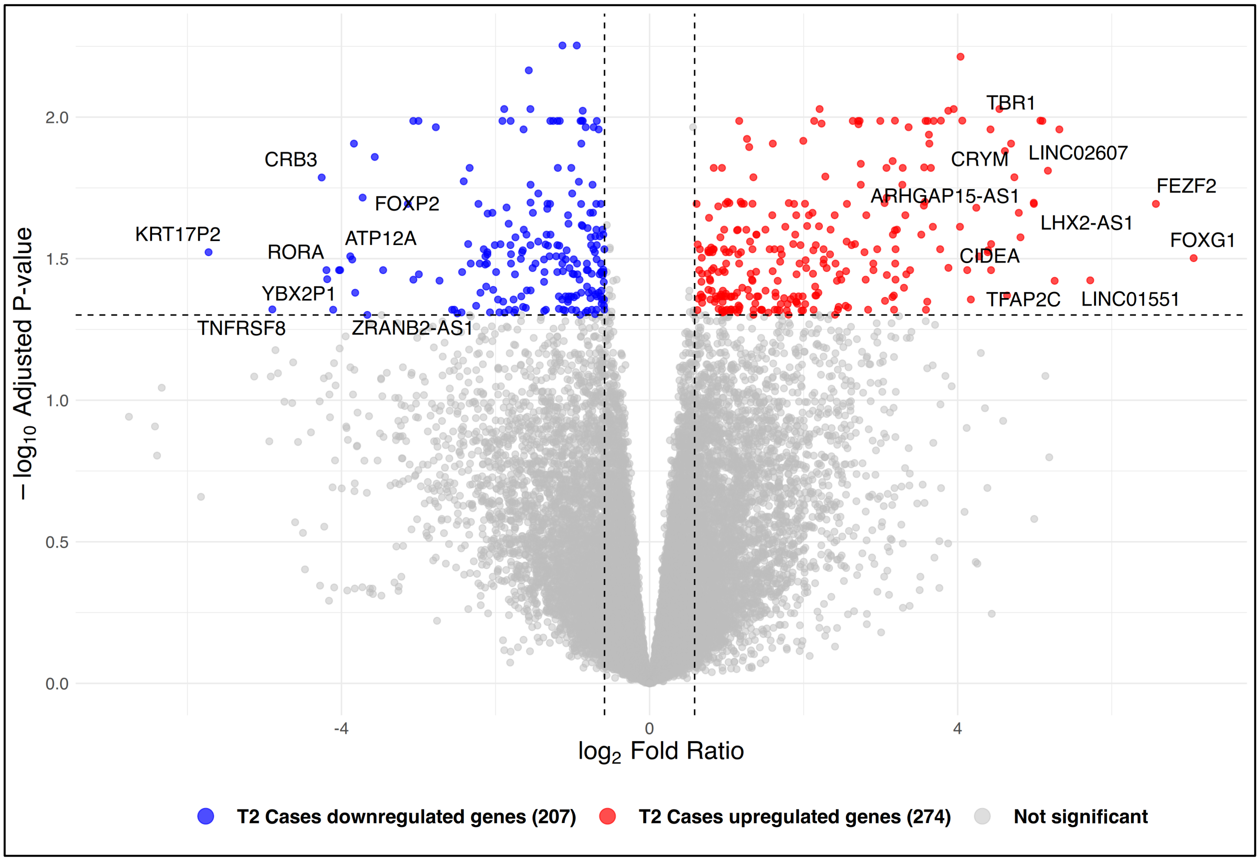


**Figure S2. Transcriptomic alterations in T2 male fetal brains associated with prenatal cannabis exposure.** Transcriptomic comparison of T2 Cases vs. Controls showing genes associated with prenatal cannabis exposure. Volcano plot showing quantified genes plotted by $\log_2$ fold ratio and Benjamini-Hochberg adjusted P value. Dashed vertical lines indicate the fold-ratio threshold ($|\log_2 FR| \geq 0.58$), and the horizontal dashed line indicates the adjusted P-value cutoff (adjusted $P \leq 0.05$). A total of 481 DEGs were identified. Selected DEGs are labeled. Significantly downregulated and upregulated genes are shown in blue and red, respectively.

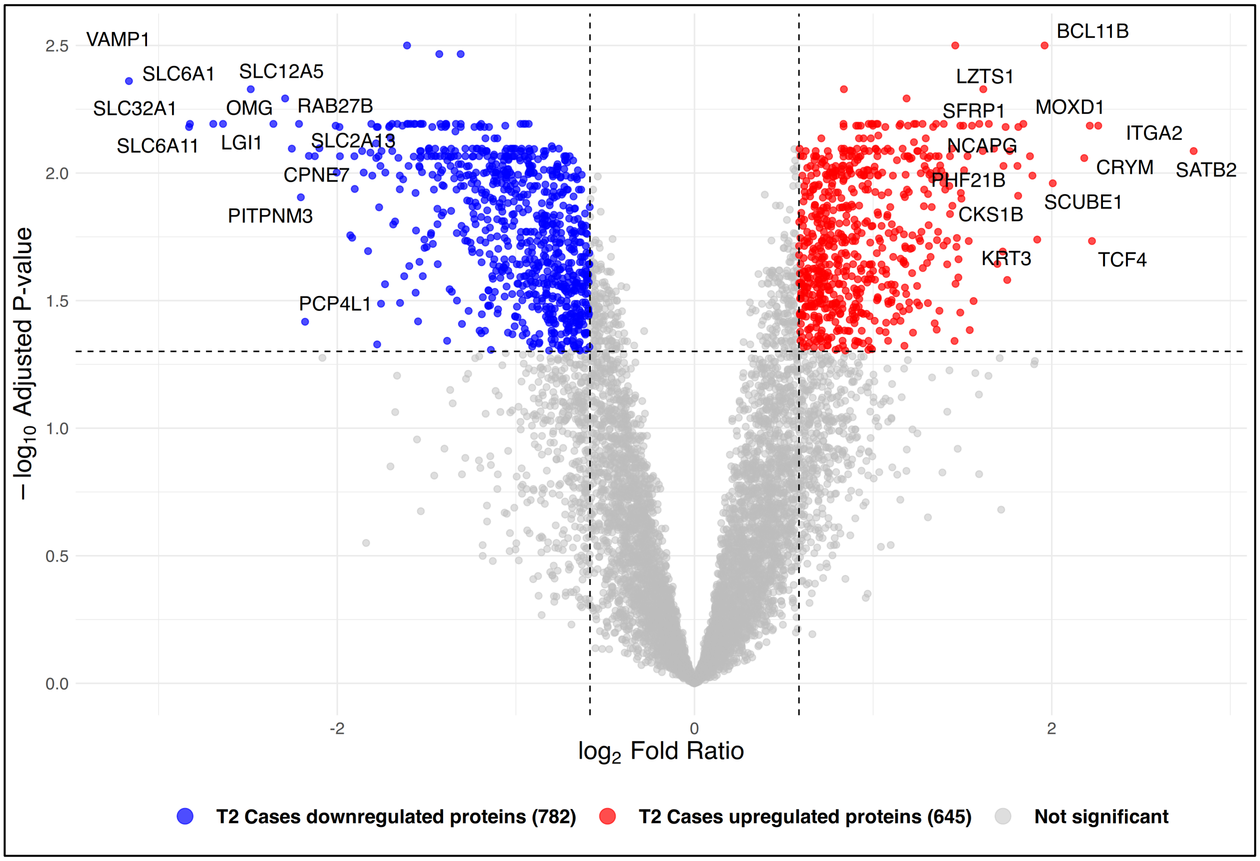


**Figure S3. Proteomic alterations in T2 male fetal brains associated with prenatal cannabis exposure.** Proteomic comparison of T2 Cases vs. Controls showing proteins associated with prenatal cannabis exposure. Volcano plot showing quantified proteins plotted by $\log_2$ fold ratio and Benjamini-Hochberg adjusted P value. Dashed vertical lines indicate the fold-ratio threshold ($|\log_2 FR| \geq 0.58$), and the horizontal dashed line indicates the adjusted P-value cutoff (adjusted P $\leq 0.05$). A total of >1,400 DEPs were identified. Significantly downregulated (blue) and upregulated (red) proteins are highlighted, with selected proteins labeled.

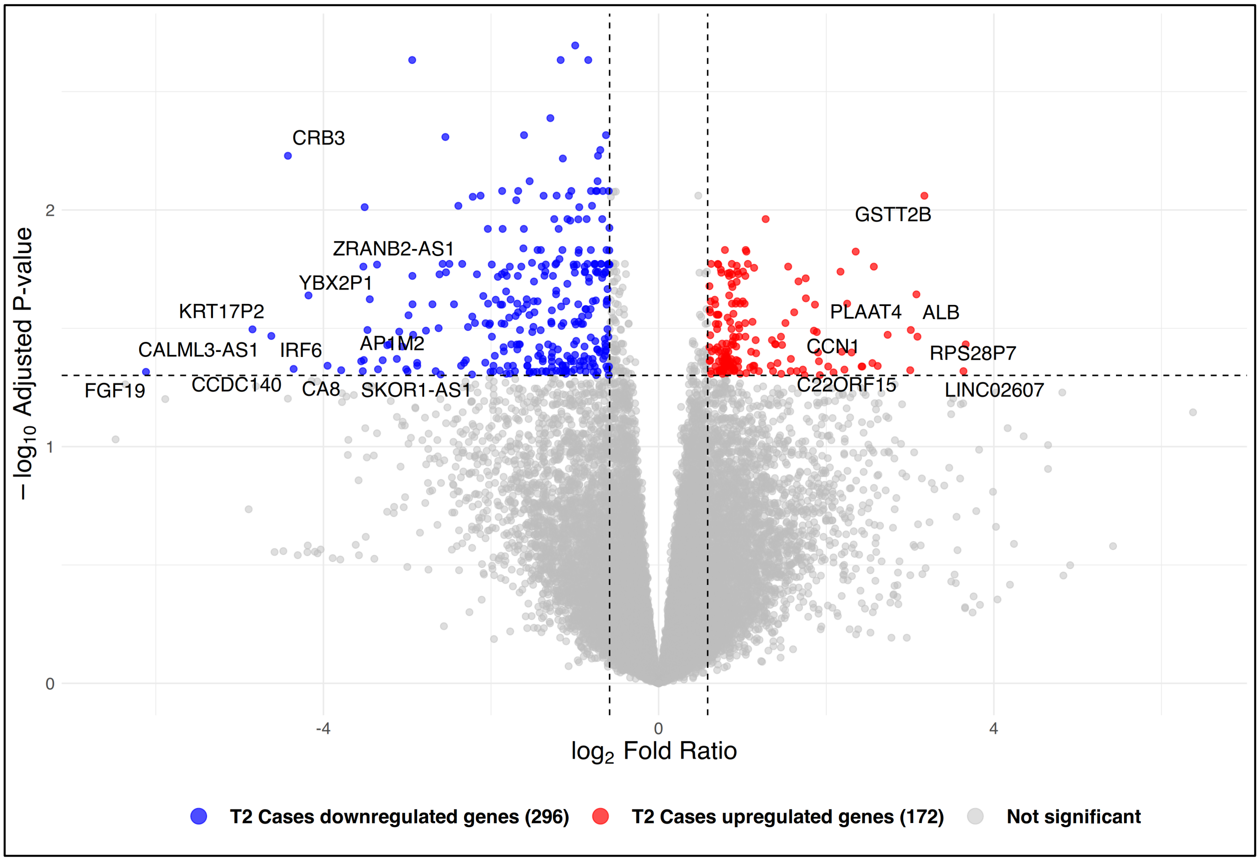


**Figure S4. Transcriptomic alterations in T2 fetal brains associated with prenatal cannabis exposure (males and females).** Transcriptomic comparison of T2 Cases vs. Controls showing genes associated with prenatal cannabis exposure. Volcano plot showing quantified genes plotted by $\log_2$ fold ratio and Benjamini-Hochberg adjusted P value. Dashed vertical lines indicate the fold-ratio threshold ($|\log_2 FR| \geq 0.58$), and the horizontal dashed line indicates the adjusted P-value cutoff (adjusted $P \leq 0.05$). A total of 468 DEGs were identified. Significantly downregulated (blue) and upregulated (red) genes are highlighted, with selected genes labeled.

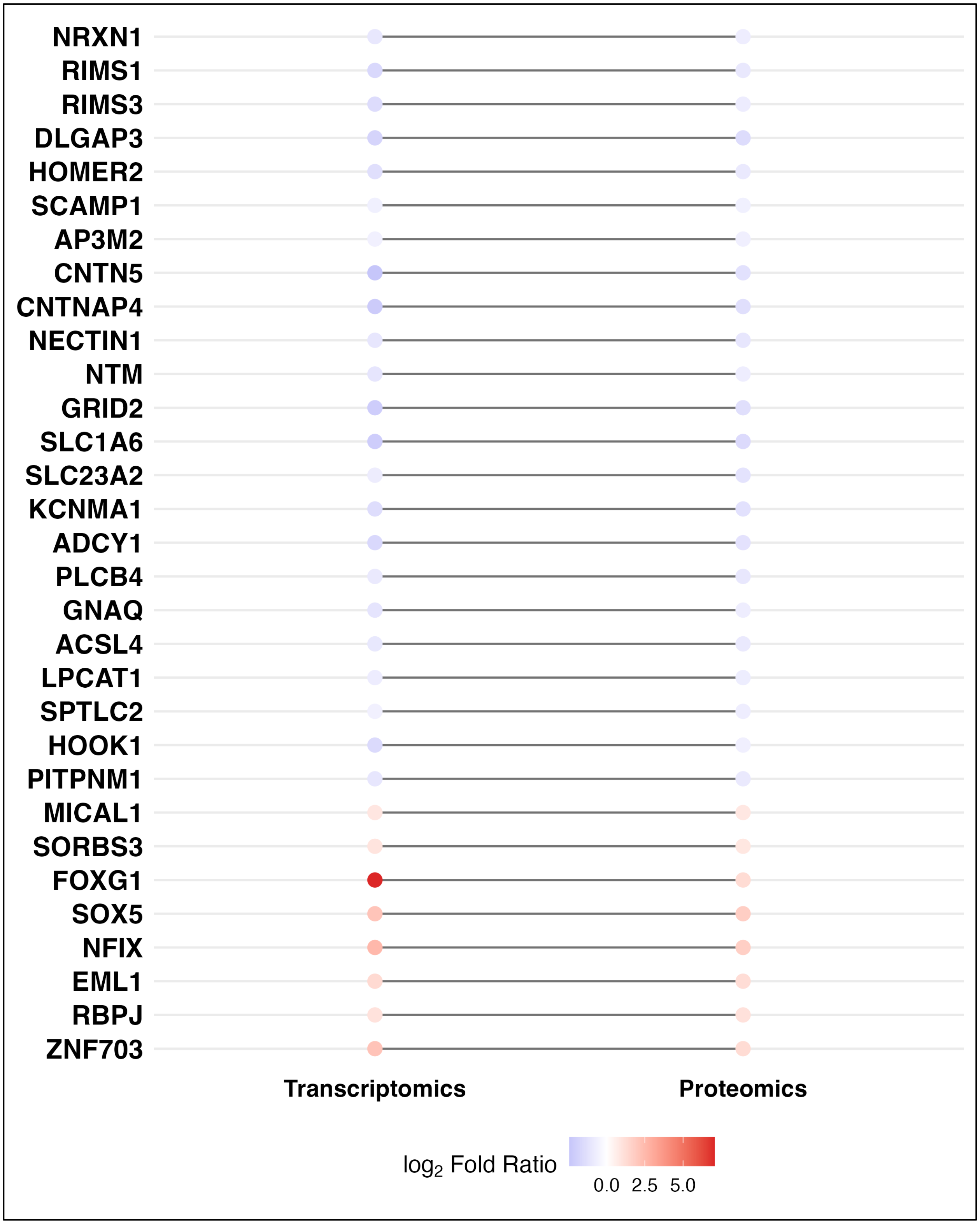


**Figure S5. Selected concordant transcriptomic and proteomic alterations in male T2 Cases vs. Controls.** Transcriptomic and proteomic expression changes for selected concordantly regulated genes and proteins that met our differential expression criteria (adjusted $P \leq 0.05$ and

$|log_2FR| \geq 0.58$) in both datasets. Blue indicates downregulation in T2 Cases, whereas red indicates upregulation in T2 Cases. Concordant reductions in T2 Cases were observed for synaptic scaffold and vesicle-associated components (NRXN1, RIMS1, RIMS3, DLGAP3, HOMER2, SCAMP1, AP3M2), neuronal adhesion systems (CNTN5, CNTNAP4, NECTIN1, NTM), excitatory signaling, glutamate/ascorbate transport, and membrane excitability regulators (GRID2, SLC1A6, SLC23A2, KCNMA1, ADCY1, PLCB4, GNAQ), lipid metabolism regulators (ACSL4, LPCAT1, SPTLC2), and intracellular trafficking proteins (HOOK1, PITPNM1). Concordant increases in T2 Cases were observed for cytoskeletal remodeling proteins (MICAL1, SORBS3) and developmental regulators (FOXG1, SOX5, NFIX, EML1, RBPJ, ZNF703). Lines connect the transcriptomics and proteomics data.

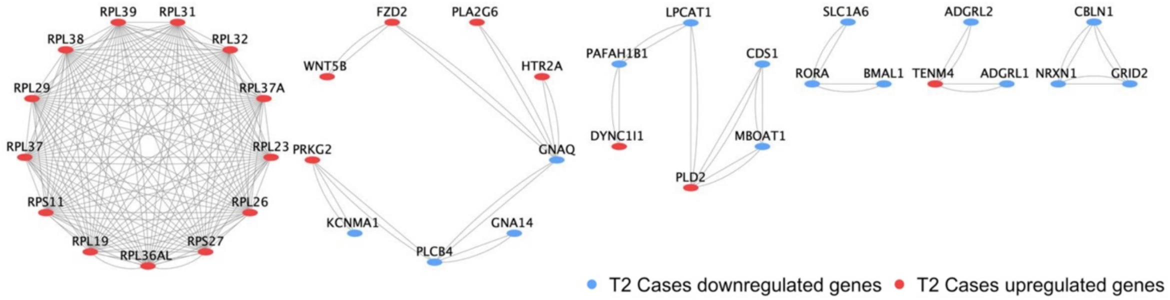


| Cluster | STRING Interaction Module Enriched For | Upregulated Genes | Downregulated Genes |
|---|---|---|---|
| A | Ribosomal Subunit Assembly and Cytosolic Translation | RPL19, RPL23, RPL26, RPL29, RPL31, RPL32, RPL36AL, RPL37, RPL37A, RPL38, RPL39, RPS11, RPS27 | |
| B | GPCR Signaling and Phospholipase Calcium Pathway | FZD2, HTR2A, PLA2G6, PRKG2, WNT5B | GNA14, GNAQ, KCNMA1, PLCB4 |
| C | Phospholipid Remodeling and Membrane Lipid Biosynthesis | DYNC1I1, PLD2 | CDS1, LPCAT1, MBOAT1, PAFAH1B1 |
| D | Circadian Transcriptional Regulation and Glutamate Transport | | BMAL1, RORA, SLC1A6 |
| E | Transsynaptic Adhesion Complexes and Neuronal Wiring | TENM4 | ADGRL1, ADGRL2 |
| F | Postsynaptic Receptor Scaffold and Glutamatergic Synapse Organization | | CBLN1, GRID2, NRXN1 |

**Figure S6. STRING-based transcriptomic interaction networks in male T2 Cases vs. Controls.** (A–F) STRING-derived transcriptomic interaction subnetworks constructed from significantly altered genes in male T2 Cases vs. Controls. Blue nodes indicate genes downregulated in T2 Cases, whereas red nodes indicate genes upregulated in T2 Cases. The table summarizes the major functional modules represented in each interaction cluster.

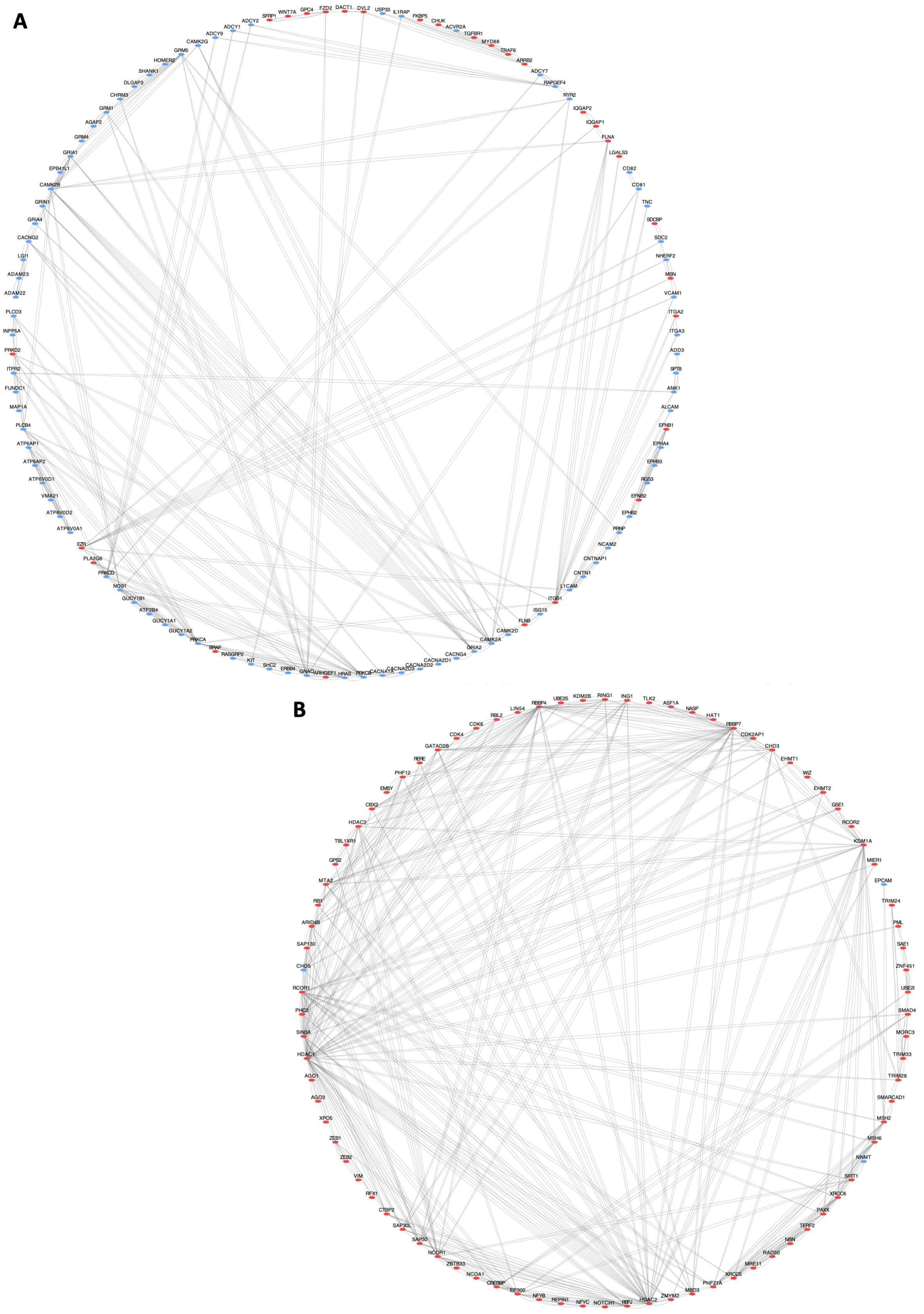
A
SFRP1
WNT7A
GPC4
FZD2
DACT1
DVL2
USP33
IL1RAP
FKBP5
CHUK
ACVR2A
TGFBR1
MYD88
TRAF6
ARRB2
ADCY7
RAPGEF4
RYR2
IQGAP2
IQGAP1
FLNA
LGALS3
CD82
CD81
TNC
SDCBP
SDC2
NHERF2
MSN
VCAM1
ITGA2
ITGA3
ADD3
SPTB
ANK1
ALCAM
EFNB1
EPHA4
EPHB3
RGS3
EFNB2
EPHB2
PRNP
NCAM2
CNTNAP1
CNTN1
L1CAM
ITGB1
ISG15
FLNB
CAMK2D
CAMK2A
GRIA2
CACNG4
CACNA2D1
CACNA2D2
CACNA2D3
CACNA1A
PRKCB
HRAS
ARHGEF1
GNAQ
ERBB4
SHC2
KIT
RASGRP2
BRAP
PRKCA
GUCY1A2
GUCY1A1
ATP2B4
GUCY1B1
NOS1
PRKCD
PLA2G6
EZR
ATP6V0A1
ATP6V0D2
VMA21
ATP6V0D1
ATP6AP2
ATP6AP1
PLCB4
MAP1A
FUNDC1
ITPR2
PRKD2
INPP5A
PLCD3
ADAM22
ADAM23
LGI1
CACNG2
GRIA4
GRIN1
CAMK2B
EPB41L1
GRIA1
GRM4
AGAP2
GRM1
CHRM3
DLGAP3
SHANK1
HOMER2
GRM5
CAMK2G
ADCY9
ADCY1
ADCY2
B
RBBP4
UBE2S
KDM2B
RING1
ING1
TLK2
ASF1A
NASP
HAT1
RBBP7
CDK2AP1
CHD3
EHMT1
WIZ
EHMT2
GSE1
RCOR2
KDM1A
MIER1
EPCAM
TRIM24
PML
SAE1
ZNF451
UBE2I
SMAD4
MORC3
TRIM33
TRIM28
SMARCAD1
MSH2
MSH6
NNMT
SIRT1
XRCC6
PAXX
TERF2
NBN
RAD50
MRE11
XRCC5
PHF21A
MBD3
ZMYM2
HDAC2
NOTCH1
NFYC
REPIN1
NFYB
EP300
CREBBP
NCOA1
ZBTB33
NCOR1
SAP30
SAP30L
CTBP2
RFX1
VIM
ZEB2
ZEB1
XPO5
AGO3
AGO1
HDAC1
SIN3A
PHC2
RCOR1
CHD5
SAP130
ARID4B
RB1
MTA2
GPS2
TBL1XR1
HDAC3
CBX2
EMSY
PHF12
RERE
GATAD2B
CDK4
CDK6
RBL2
LIN54

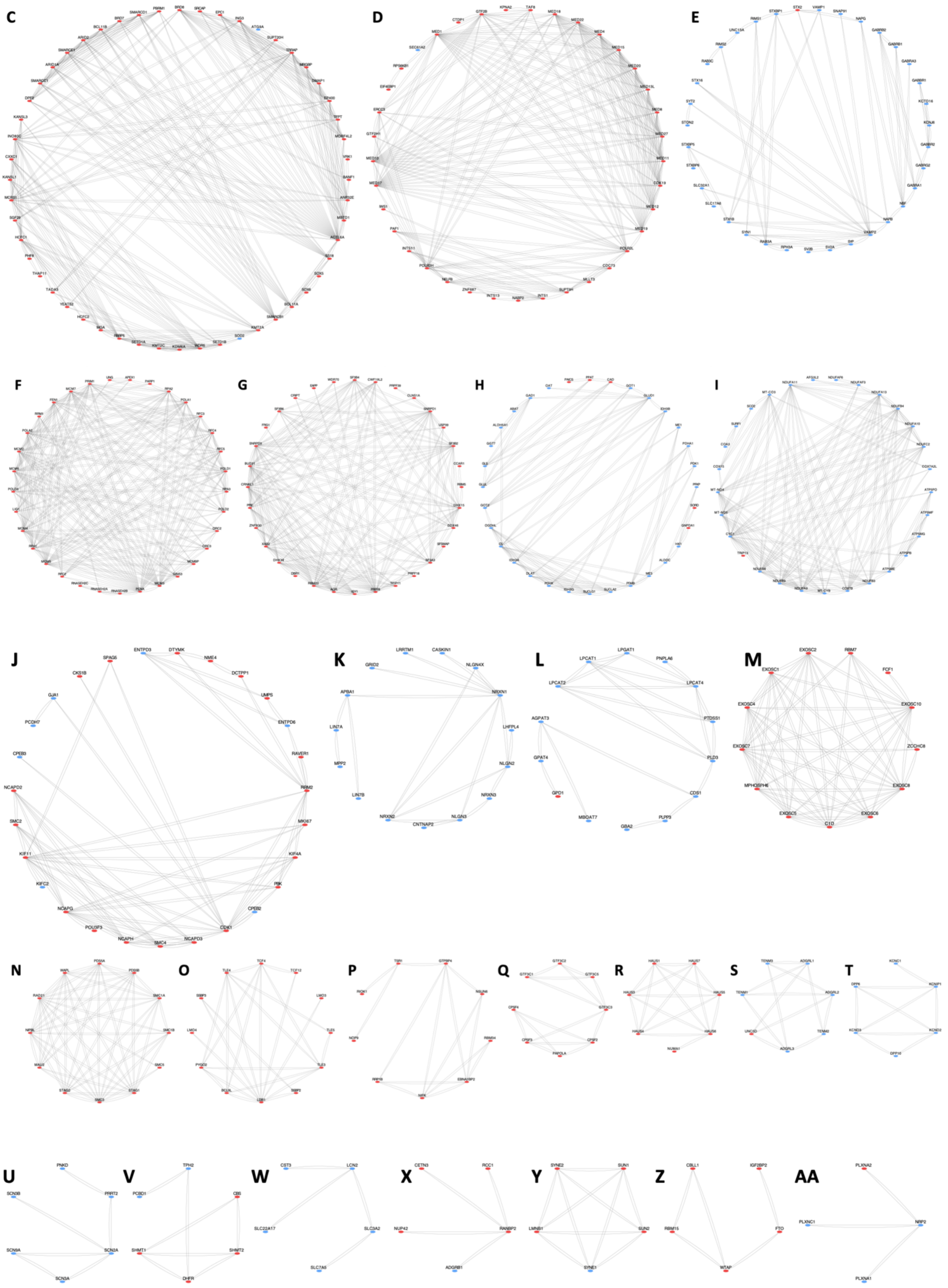
C
D
E
F
G
H
I
J
SPAG5
ENTPD3
DTYMK
NME4
CKS1B
DCTPP1
GJA1
UMPS
PCDH7
ENTPD6
CPEB3
RAVER1
NCAPD2
RRM2
SMC2
MKI67
KIF11
KIF4A
KIFC2
PBK
NCAPG
CPEB2
POU3F3
CDK1
NCAPH
SMC4
NCAPD3
K
LRRTM1
CASKIN1
GRID2
NLGN4X
APBA1
NRXN1
LIN7A
LHFPL4
MPP2
NLGN2
LIN7B
NRXN3
NRXN2
CNTNAP2
NLGN3
L
LPCAT1
LPGAT1
PNPLA6
LPCAT2
LPCAT4
AGPAT3
PTDSS1
GPAT4
PLD3
GPD1
CDS1
MBOAT7
GBA2
PLPP3
M
EXOSC2
RBM7
EXOSC1
FCF1
EXOSC4
EXOSC10
EXOSC7
ZCCHC8
MPHOSPH6
EXOSC8
EXOSC5
C1D
EXOSC6
N
O
P
Q
R
S
T
U
PNKD
SCN3B
PRRT2
SCN9A
SCN2A
SCN3A
V
TPH2
PCBD1
CBS
SHMT1
SHMT2
DHFR
W
CST3
LCN2
SLC22A17
SLC3A2
SLC7A5
X
CETN3
RCC1
NUP42
RANBP2
ADGRB1
Y
SYNE2
SUN1
LMNB1
SUN2
SYNE1
Z
CBLL1
IGF2BP2
RBM15
FTO
WTAP
AA
PLXNA2
PLXNC1
NRP2
PLXNA1

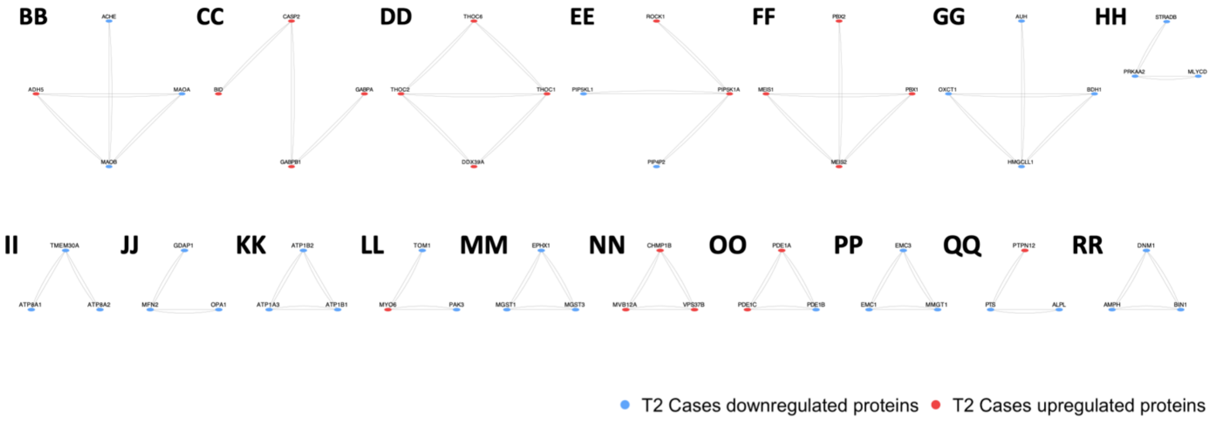


| Cluster | STRING Interaction Module Enriched For | Upregulated Proteins | Downregulated Proteins |
|---|---|---|---|
| A | Calcium-Dependent Signaling and Synaptic Plasticity Pathways | ARHGEF1, ARRB2, BRAP, CHUK, DACT1, DVL2, EFNB1, EFNB2, EZR, FKBP5, FLNA, FLNB, FZD2, GPC4, IQGAP1, IQGAP2, ITGA2, ITGB1, LGALS3, MSN, MYD88, PLA2G6, PRKD2, SDCBP, SFRP1, TGFBR1, TRAF6, WNT7A | ACVR2A, ADAM22, ADAM23, ADCY1, ADCY2, ADCY7, ADCY9, ADD3, AGAP2, ALCAM, ANK1, ATP2B4, ATP6AP1, ATP6AP2, ATP6V0A1, ATP6V0D1, ATP6V0D2, CACNA1A, CACNA2D1, CACNA2D2, CACNA2D3, CACNG2, CACNG4, CAMK2A, CAMK2B, CAMK2D, CAMK2G, CD81, CD82, CHRM3, CNTN1, CNTNAP1, DLGAP3, EPB41L1, EPHA4, EPHB2, EPHB3, ERBB4, FUNDC1, GNAQ, GRIA1, GRIA2, GRIA4, GRIN1, GRM1, GRM4, GRM5, GUCY1A1, GUCY1A2, GUCY1B1, HOMER2, HRAS, IL1RAP, INPP5A, ISG15, ITGA3, ITPR2, KIT, L1CAM, LGI1, MAP1A, NCAM2, NHERF2, NOS1, PLCB4, PLCD3, PRKCA, PRKCB, PRKCD, PRNP, RAPGEF4, RASGRP2, RGS3, RYR2, SDC2, SHANK1, SHC2, SPTB, TNC, USP33, VCAM1, VMA21 |
| AA | Plexin-Semaphorin Signaling and Axon Guidance | PLXNA2 | NRP2, PLXNA1, PLXNC1 |
| B | Chromatin Remodeling and Histone Deacetylation | AGO1, AGO3, ARID4B, ASF1A, CBX2, CDK2AP1, CDK4, CDK6, CHD3, CREBBP, CTBP2, EHMT1, EHMT2, EMSY, EP300, GATAD2B, GPS2, GSE1, HAT1, HDAC1, HDAC2, HDAC3, ING1, KDM1A, KDM2B, LIN54, MBD3, MIER1, MORC3, MRE11, MSH2, MSH6, MTA2, NASP, NBN, NCOA1, NCOR1, NFYB, NFYC, NOTCH1, PAXX, PHC2, PHF12, PHF21A, PML, RAD50, RB1, RBBP4, RBBP7, RBL2, RBPJ, RCOR1, RCOR2, REPIN1, RERE, RFX1, RING1, SAE1, SAP130, SAP30, SAP30L, SIN3A, SIRT1, SMAD4, SMARCAD1, TBL1XR1, TERF2, TLK2, TRIM24, TRIM28, TRIM33, UBE2I, UBE2S, VIM, WIZ, XPO5, XRCC5, XRCC6, ZBTB33, ZEB1, ZEB2, ZMYM2, ZNF451 | CHD5, EPCAM, NNMT |
| BB | Monoamine Oxidation and Neurotransmitter Catabolism | ADH5 | ACHE, MAOA, MAOB |
| C | SWI/SNF Chromatin Remodeling and Transcriptional Regulation | ACTL6A, ANP32E, ARID1A, ARID2, BANF1, BCL11A, BCL11B, BRD7, BRD8, CXXC1, DMAP1, DPF2, EP400, EPC1, HCFC1, HCFC2, ING3, INO80C, KANSL1, KANSL3, KDM6A, KMT2A, KMT2C, MBTD1, MCRS1, MGA, MORF4L2, MRGBP, PBRM1, PHF8, RBBP5, SETD1A, SETD1B, SGF29, SMARCB1, SMARCC1, SMARCD1, SMARCE1, SOX5, SOX6, SRCAP, SS18, SUPT20H, TADA3, TFPT, THAP11, TRRAP, VRK1, WDR5, YEATS2 | ATG9A, SOD2 |
| CC | Apoptotic Signaling and GABP Transcriptional Regulation | BID, CASP2, GABPA, GABPB1 | |
| D | Mediator Complex and RNA Polymerase II Transcription | CDC73, CDK19, CTDP1, EIF4EBP1, ERCC3, GTF2B, GTF2H1, INTS1, INTS11, INTS13, IWS1, KPNA2, MED1, MED10, MED11, MED12, MED13L, MED15, MED17, MED18, MED19, MED20, MED22, MED27, MED4, MED8, MLLT3, NABP2, NELFB, PAF1, POLR2H, POLR2L, RPS6KB1, SUPT5H, TAF8, ZNF687 | SEC61A2 |
| DD | mRNA Nuclear Export and TREX Complex Function | DDX39A, THOC1, THOC2, THOC6 | |
| E | Synaptic Vesicle Fusion and Presynaptic Exocytosis | STX2 | GABBR1, GABBR2, GABRA1, GABRA3, GABRB1, GABRB2, GABRG2, KCNJ6, KCTD16, NAPB, NAPG, NSF, RAB3A, RAB3C, RIMS1, RIMS2, RPH3A, SLC17A6, SLC32A1, SNAP91, STON2, STX16, STX1B, STXBP1, STXBP5, STXBP6, SV2A, SV2B, SYN1, SYP, SYT2, UNC13A, VAMP1, VAMP2 |
| EE | Phosphoinositide Signaling and PI(4,5)$P_2$ Metabolism | PIP5K1A, ROCK1 | PIP4P2, PIP5KL1 |
| F | DNA Replication Licensing and Replication Fork Progression | APEX1, FEN1, GINS3, LIG1, MCM2, MCM3, MCM4, MCM5, MCM6, MCM7, MCMBP, ORC2, ORC3, PARP1, PCNA, POLA1, POLA2, POLD1, POLD2, POLD3, PRIM1, RFC2, RFC3, RFC4, RFC5, RNASEH2A, RNASEH2B, RNASEH2C, RPA1, RPA2, RPA3, RRM1, UNG | |
| FF | MEIS Homeobox Transcription Factors and Developmental Patterning | MEIS1, MEIS2, PBX1, PBX2 | |
| G | Pre-mRNA Splicing and Spliceosome Assembly | AQR, BUD31, CCAR1, CLNS1A, CRIPT, CRNKL1, CWF19L2, DBR1, DDX46, DHX15, DHX38, EAPP, FRG1, ISY1, PPIE, PRPF18, PRPF39, PRPF8, RBM22, RBM5, SF3A3, SF3B2, SF3B4, SF3B6, SFSWAP, SNRPD1, SNRPD3, TFIP11, USP39, WDR70, XAB2, ZNF830 | |
| GG | Ketone Body Metabolism and Mitochondrial Energy Utilization | | AUH, BDH1, HMGCLL1, OXCT1 |

| | | | |
|---|---|---|---|
| H | Central Carbon Metabolism and Amino Acid Biosynthesis | CAD, GNPDA1, PAICS, PPAT, SORD | ABAT, ALDH5A1, ALDOC, CS, DLAT, GAD1, GGT7, GLS, GLUD1, GLUL, GOT1, GOT2, HK1, IDH3A, IDH3B, IDH3G, ME1, ME3, OAT, OGDHL, PDHA1, PDHB, PDHX, PDK1, PFKP, SUCLA2, SUCLG1 |
| HH | AMPK Energy Sensing and Metabolic Stress Signaling | | MLYCD, PRKAA2, STRADB |
| I | Mitochondrial Respiratory Chain and Oxidative Phosphorylation | TRIP13 | AFG3L2, ATP5ME, ATP5MF, ATP5MG, ATP5PB, ATP5PO, COA3, COX15, COX7A2L, COX7B, CYC1, MT-CO3, MT-CYB, MT-ND4, MT-ND5, NDUFA10, NDUFA11, NDUFA13, NDUFA9, NDUFAF3, NDUFAF6, NDUFB3, NDUFB4, NDUFB9, NDUFC2, NDUFS8, SCO2, SURF1 |
| II | Phospholipid Flippase Activity and Membrane Asymmetry | | ATP8A1, ATP8A2, TMEM30A |
| J | Cell Cycle Progression and Mitotic Chromosome Segregation | CDK1, CKS1B, DCTPP1, DTYMK, KIF11, KIF4A, MKI67, NCAPD2, NCAPD3, NCAPG, NCAPH, NME4, PBK, POU3F3, RAVER1, RRM2, SMC2, SMC4, SPAG5, UMPS | CPEB2, CPEB3, ENTPD3, ENTPD6, GJA1, KIFC2, PCDH7 |
| JJ | Mitochondrial Dynamics and Outer Membrane Remodeling | | GDAP1, MFN2, OPA1 |
| K | Neurexin Adhesion Complex and Synaptic Organization | | APBA1, CASKIN1, CNTNAP2, GRID2, LHFPL4, LIN7A, LIN7B, LRRTM1, MPP2, NLGN2, NLGN3, NLGN4X, NRXN1, NRXN2, NRXN3 |
| KK | $Na^{+}/K^{+}$-ATPase Ion Transport and Neuronal Membrane Potential | | ATP1A3, ATP1B1, ATP1B2 |
| L | Phospholipid Remodeling and Membrane Lipid Metabolism | GPD1 | AGPAT3, CDS1, GBA2, GPAT4, LPCAT1, LPCAT2, LPCAT4, LPGAT1, MBOAT7, PLD3, PLPP3, PNPLA6, PTDSS1 |
| LL | Actin-Based Vesicle Transport and Synaptic Trafficking | MYO6 | PAK3, TOM1 |
| M | RNA Exosome and Nuclear RNA Surveillance | C1D, EXOSC1, EXOSC10, EXOSC2, EXOSC4, EXOSC5, EXOSC6, EXOSC7, EXOSC8, FCF1, MPHOSPH6, RBM7, ZCCHC8 | |
| MM | Glutathione Conjugation and Xenobiotic Detoxification | | EPHX1, MGST1, MGST3 |
| N | Cohesin Complex and Sister Chromatid Cohesion | MAU2, NIPBL, PDS5A, PDS5B, RAD21, SMC1A, SMC1B, SMC3, SMC5, STAG1, STAG2, WAPL | |
| NN | ESCRT-I Complex and Endosomal Cargo Sorting | CHMP1B, MVB12A, VPS37B | |
| O | LDB1 Transcriptional Complex and Developmental Gene Regulation | BCL9L, LDB1, LMO3, LMO4, PYGO2, SSBP2, SSBP3, TCF12, TCF4, TLE3, TLE4, TLE5 | |
| OO | Calcium-Dependent Phosphodiesterases and cAMP/cGMP Signaling | PDE1A, PDE1C | PDE1B |
| P | Ribosome Biogenesis and Nucleolar rRNA Processing | EBNA1BP2, GTPBP4, NIFK, NOP9, NSUN6, RBM34, RIOK1, RRP1B, TSR1 | |
| PP | Endoplasmic Reticulum Membrane Complex and Protein Insertion | | EMC1, EMC3, MMGT1 |
| Q | RNA Polymerase III Transcription and mRNA 3′ End Processing | CPSF2, CPSF3, CPSF4, GTF3C1, GTF3C2, GTF3C3, GTF3C5, PAPOLA | |
| QQ | Pterin Metabolism and Tyrosine Hydroxylase Cofactor Pathways | PTPN12 | ALPL, PTS |
| R | Mitotic Spindle Assembly and Microtubule Organization | HAUS1, HAUS3, HAUS4, HAUS5, HAUS6, HAUS7, NUMA1 | |
| RR | Synaptic Endocytosis and Membrane Curvature Machinery | | AMPH, BIN1, DNM1 |
| S | Latrophilin Adhesion Receptors and Synapse Formation | UNC5D | ADGRL1, ADGRL2, ADGRL3, TENM1, TENM2, TENM3 |
| T | Voltage-Gated Potassium Channels and Neuronal Excitability | | DPP10, DPP6, KCNC1, KCND2, KCND3, KCNIP1 |
| U | Voltage-Gated Sodium Channels and Neuronal Action Potentials | | PNKD, PRRT2, SCN2A, SCN3A, SCN3B, SCN9A |
| V | One-Carbon Metabolism and Folate Cycle Enzymes | CBS, DHFR, SHMT1, SHMT2 | PCBD1, TPH2 |
| W | Iron Transport and Lipocalin-Associated Immune Response | | CST3, LCN2, SLC22A17, SLC3A2, SLC7A5 |
| X | Nuclear Pore Transport and Ran-Dependent Nuclear Trafficking | CETN3, NUP42, RANBP2, RCC1 | ADGRB1 |
| Y | Nuclear Lamina Structure and Nucleocytoskeletal Coupling | LMNB1, SUN1, SUN2, SYNE2 | SYNE1 |
| Z | m6A RNA Methylation and Post-Transcriptional Regulation | CBLL1, FTO, IGF2BP2, RBM15, WTAP | |

**Figure S7. STRING-based proteomics interaction networks in male T2 Cases vs. Controls.** (A–RR) STRING-derived proteomic interaction subnetworks constructed from significantly altered proteins in male T2 Cases vs. Controls. Blue nodes indicate proteins downregulated in T2 Cases, whereas red nodes indicate proteins upregulated in T2 Cases. The table summarizes the major functional modules represented in each interaction cluster.

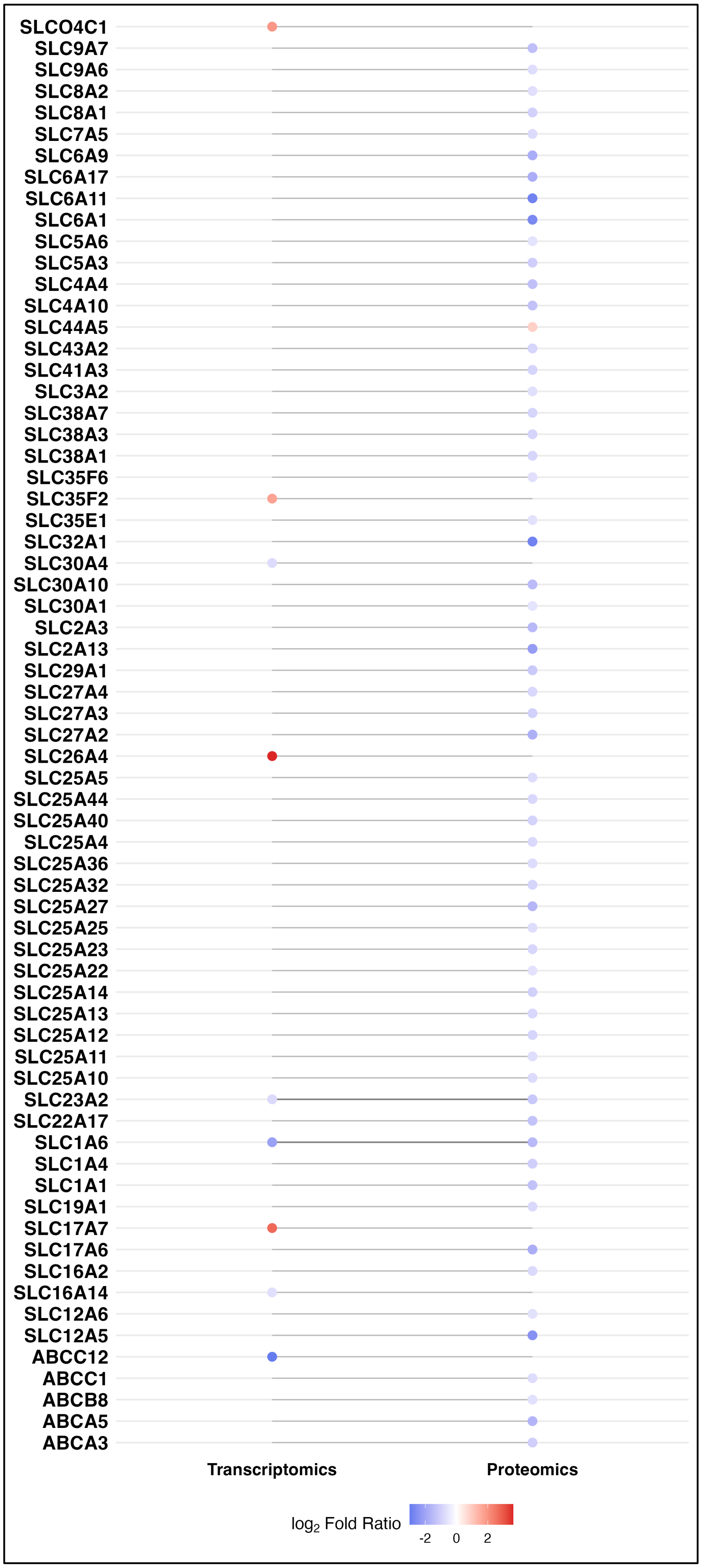
SLCO4C1
SLC9A7
SLC9A6
SLC8A2
SLC8A1
SLC7A5
SLC6A9
SLC6A17
SLC6A11
SLC6A1
SLC5A6
SLC5A3
SLC4A4
SLC4A10
SLC44A5
SLC43A2
SLC41A3
SLC3A2
SLC38A7
SLC38A3
SLC38A1
SLC35F6
SLC35F2
SLC35E1
SLC32A1
SLC30A4
SLC30A10
SLC30A1
SLC2A3
SLC2A13
SLC29A1
SLC27A4
SLC27A3
SLC27A2
SLC26A4
SLC25A5
SLC25A44
SLC25A40
SLC25A4
SLC25A36
SLC25A32
SLC25A27
SLC25A25
SLC25A23
SLC25A22
SLC25A14
SLC25A13
SLC25A12
SLC25A11
SLC25A10
SLC23A2
SLC22A17
SLC1A6
SLC1A4
SLC1A1
SLC19A1
SLC17A7
SLC17A6
SLC16A2
SLC16A14
SLC12A6
SLC12A5
ABCC12
ABCC1
ABCB8
ABCA5
ABCA3
Transcriptomics
Proteomics
$\log_2$ Fold Ratio
-2
0
2

**Figure S8. Coordinated transporter dysregulation in T2 cannabis-exposed male fetal brains.** Transcriptomic and proteomic $\log_2$ fold ratios (T2 Cases vs. Controls) are shown for all significantly altered solute carrier (SLC) and ATP-binding cassette (ABC) transporters identified (adjusted $P \leq 0.05$, $|\log_2 FR| \geq 0.58$). Each gene is displayed across molecular layers when detected, with lines connecting transcriptomic and proteomic measurements where both were observed. Proteomic data reveal a broad and consistent reduction in transporter abundance, whereas transcriptomic changes were fewer and more heterogeneous. Together, these data indicate a global decrease in membrane transport capacity at the protein level in T2 cannabis-exposed male fetal brains.

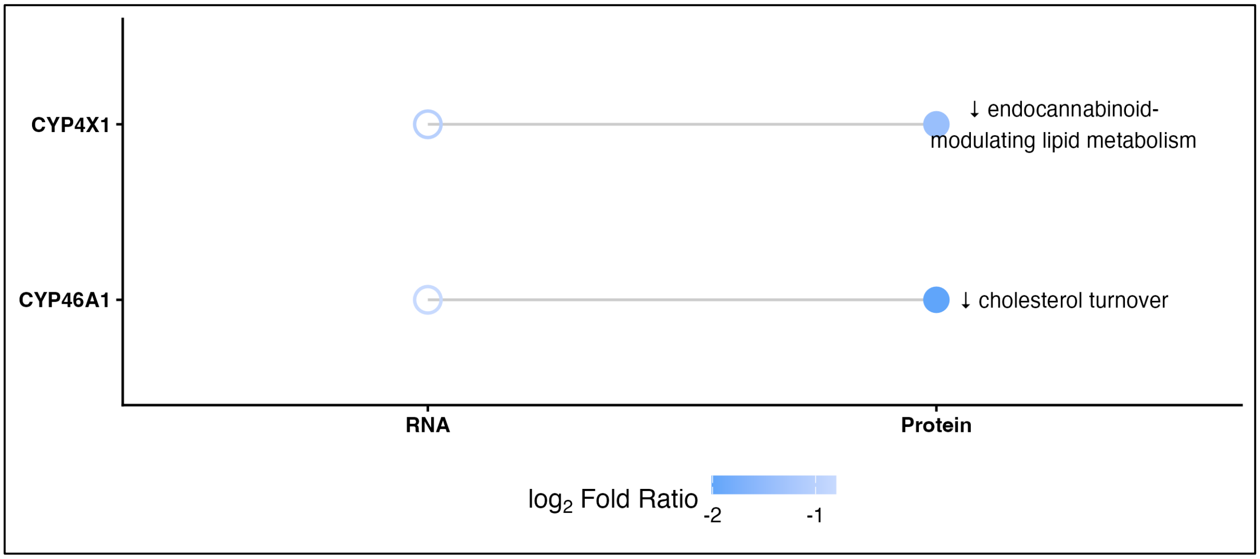


**Figure S9. Disruption of cholesterol turnover and endocannabinoid-related lipid metabolism in second-trimester male fetal brains.** Transcriptomic and proteomic $\log_2$ fold ratios (T2 Cases vs. Controls) are shown for CYP46A1 and CYP4X1 in male samples. Points represent $\log_2$ fold ratios, with color indicating the magnitude of the ratio. Opaque points denote statistically significant ratios (adjusted $P \leq 0.05$), whereas semi-transparent points indicate non-significant ratios.

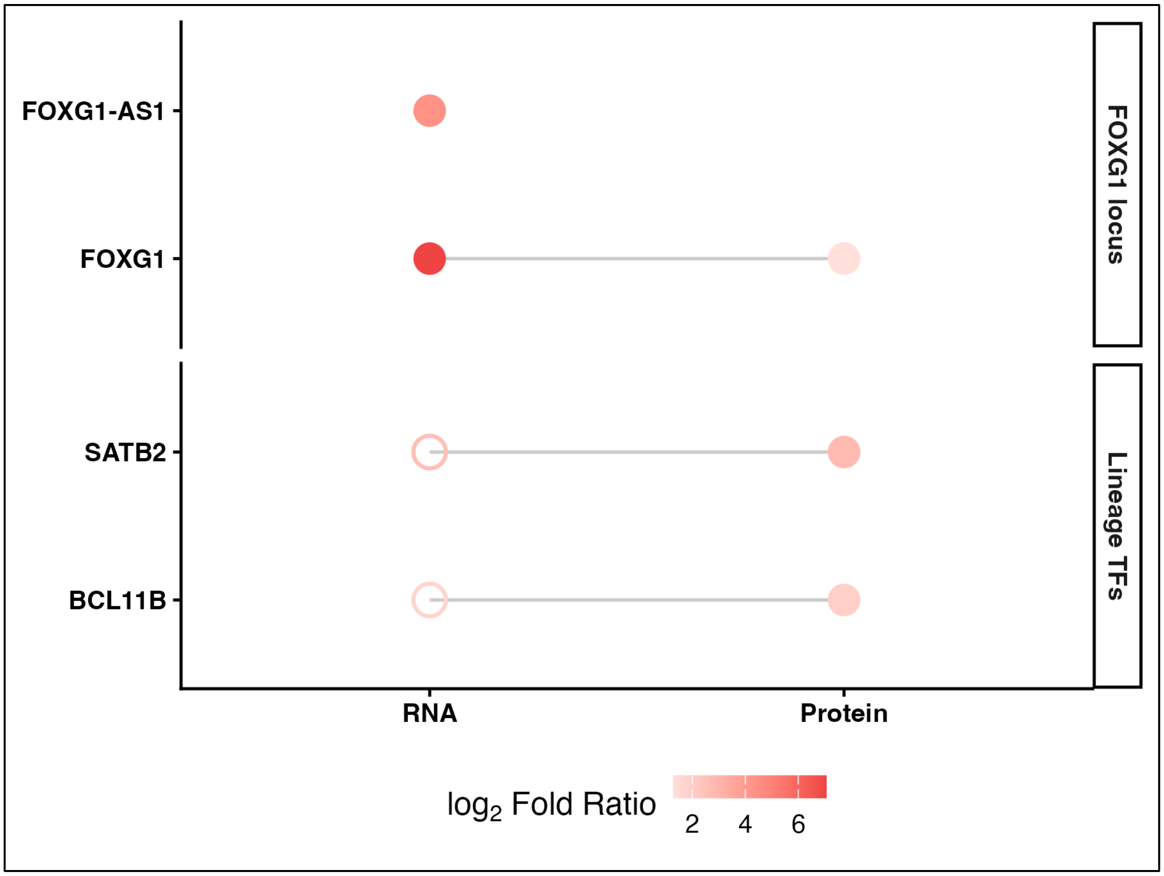


**Figure S10. Activation of developmental transcriptional programs in T2 male fetal brains, T2 Cases vs. Controls.** RNA and protein abundance changes for FOXG1, FOXG1-AS1, BCL11B, and SATB2 in T2 Cases vs. Controls. Points represent $\log_2$ fold ratio, with color indicating magnitude of the ratio. Opaque points denote statistically significant ratios (adjusted P $\leq 0.05$), whereas semi-transparent points indicate non-significant ratios. FOXG1-AS1 is represented at the RNA level only. Genes are grouped by functional category (FOXG1 locus and lineage transcription factors, TFs).

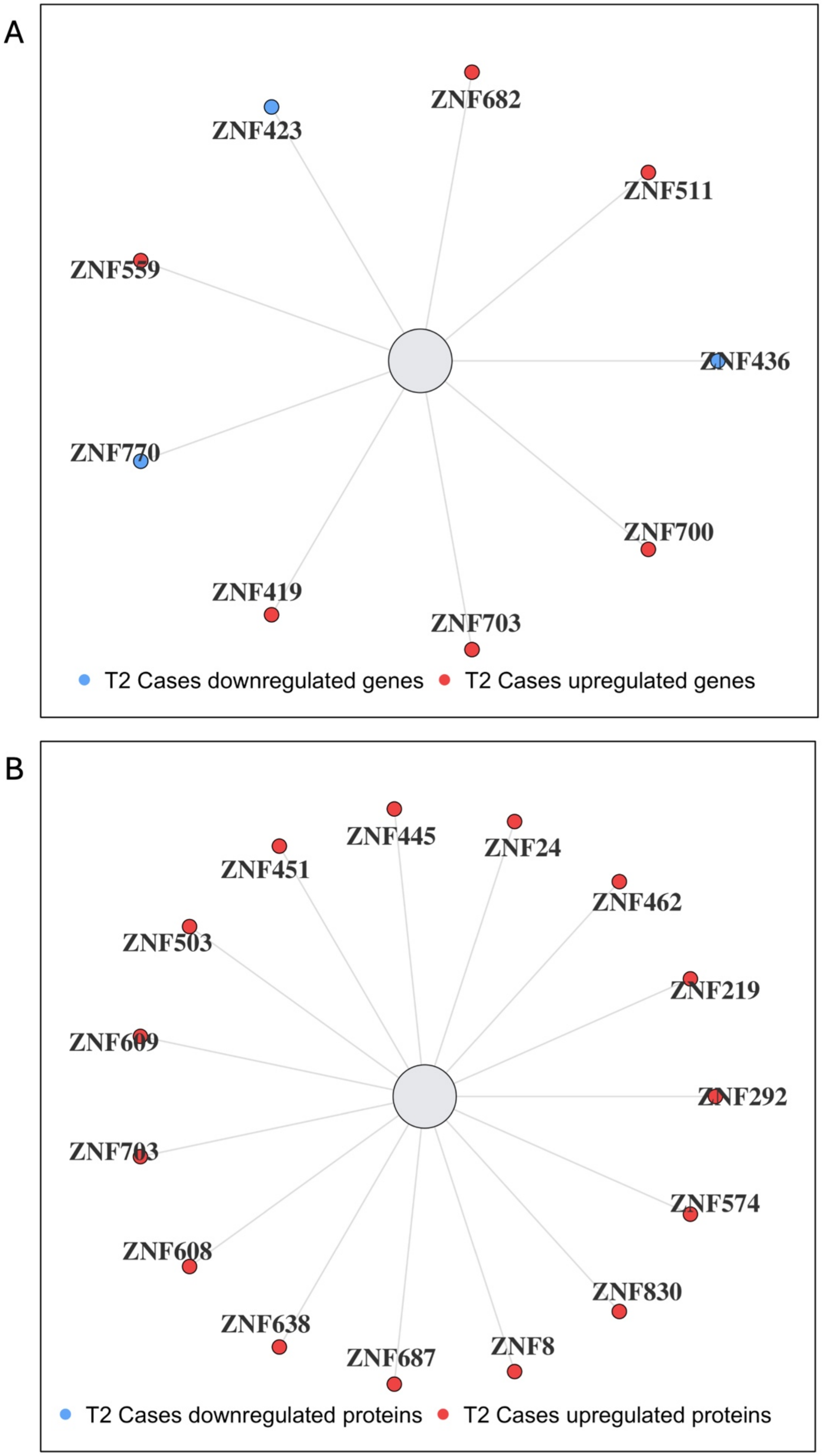
A
ZNF682
ZNF423
ZNF511
ZNF559
ZNF436
ZNF770
ZNF700
ZNF419
ZNF703
T2 Cases downregulated genes
T2 Cases upregulated genes
B
ZNF445
ZNF451
ZNF24
ZNF462
ZNF503
ZNF219
ZNF609
ZNF292
ZNF703
ZNF574
ZNF608
ZNF830
ZNF638
ZNF687
ZNF8
T2 Cases downregulated proteins
T2 Cases upregulated proteins

**Figure S11. Zinc finger transcription factor expression in male fetal brains, T2 Cases vs. Controls. (A)** Transcriptomic and **(B)** proteomic changes in zinc finger transcription factors in T2 Cases versus Controls. Nodes represent individual genes or proteins, colored by direction of ratio (red, upregulated; blue, downregulated) based on $\log_2$ fold ratio. Only statistically significant features (adjusted $P \leq 0.05$) are shown.

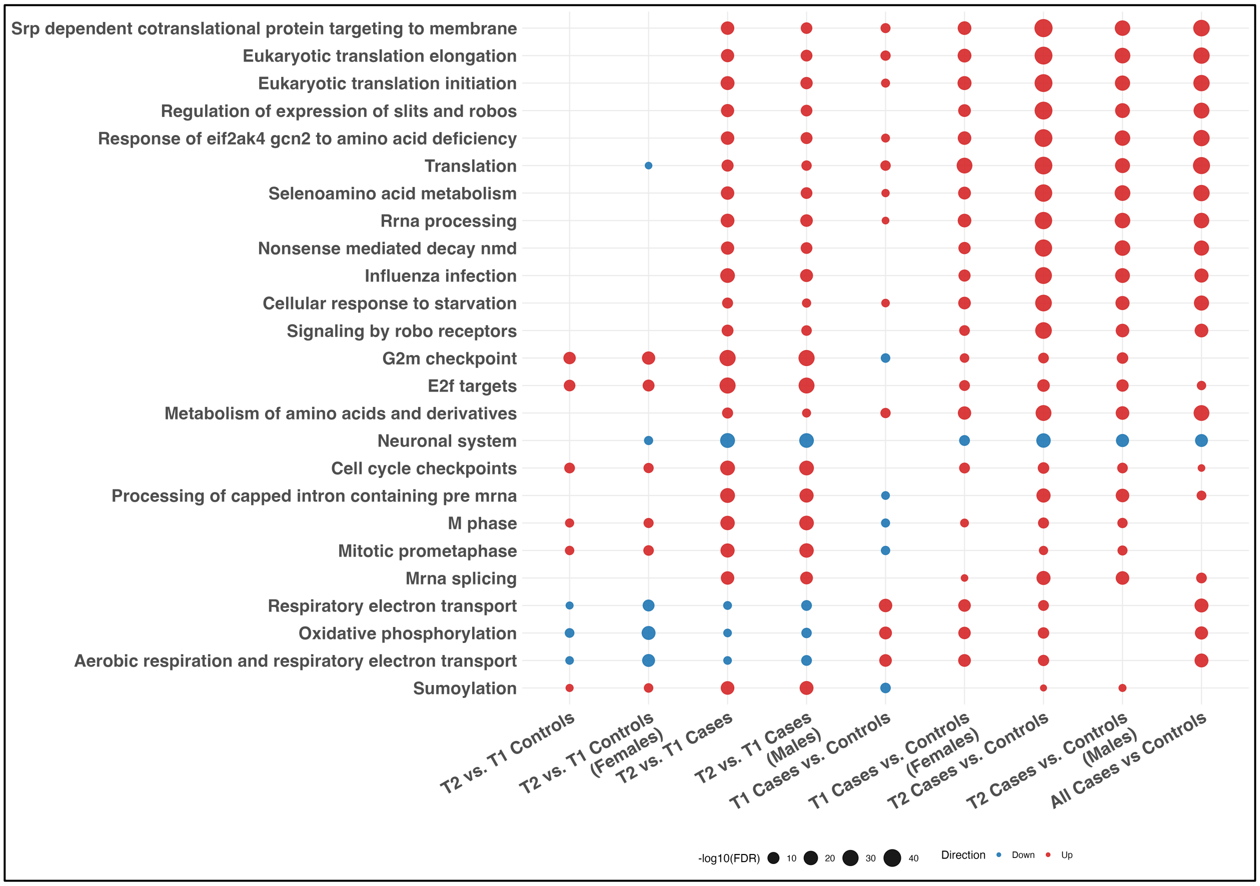


**Figure S12. Gene set enrichment analysis (GSEA) of transcriptomic changes in fetal brains across multiple comparisons.** Dot plot showing enriched pathways across multiple transcriptomic contrasts, with pathways ranked on the y-axis and comparisons on the x-axis. Point size reflects enrichment significance ($-\log_{10}$ FDR), and color indicates direction of enrichment (red, upregulated; blue, downregulated). Neuronal signaling pathways are consistently downregulated in cannabis exposure comparisons, whereas ribosomal, translational, and SRP-dependent targeting pathways, along with amino acid stress response programs, are enriched, indicating global reweighting of the transcriptome.

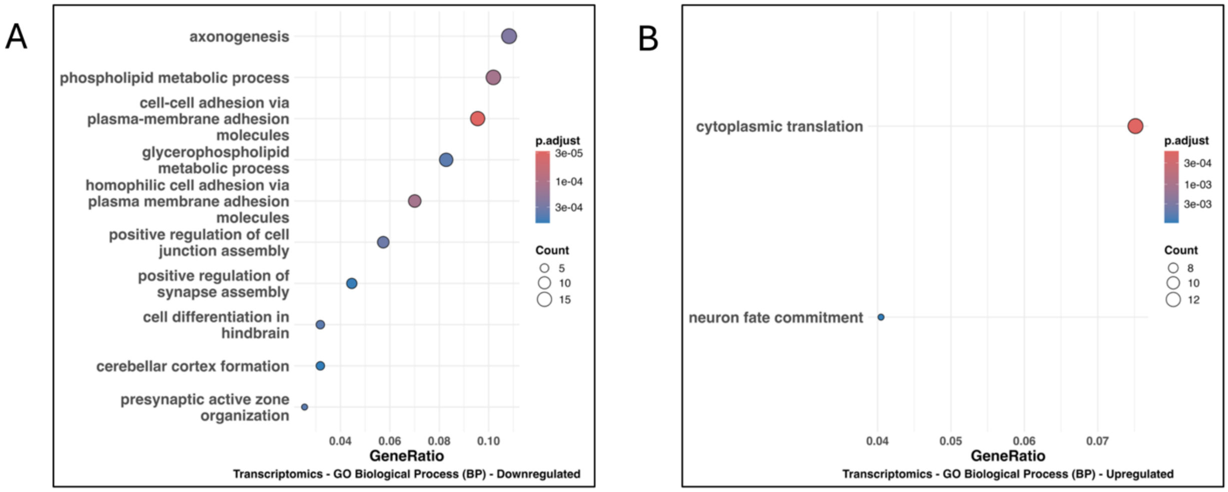


**Figure S13. GO Biological Process enrichment analysis of transcriptomic changes in male T2 Cases vs. Controls.** Gene Ontology (Biological Process) enrichment analysis of downregulated **(A)** and upregulated **(B)** genes. Dot plots show enriched terms ranked by gene ratio, with point size indicating gene count and color representing adjusted P value. Downregulated genes were enriched for axonogenesis, cell-cell adhesion, and phospholipid metabolic processes, whereas upregulated genes were enriched for cytoplasmic translation.

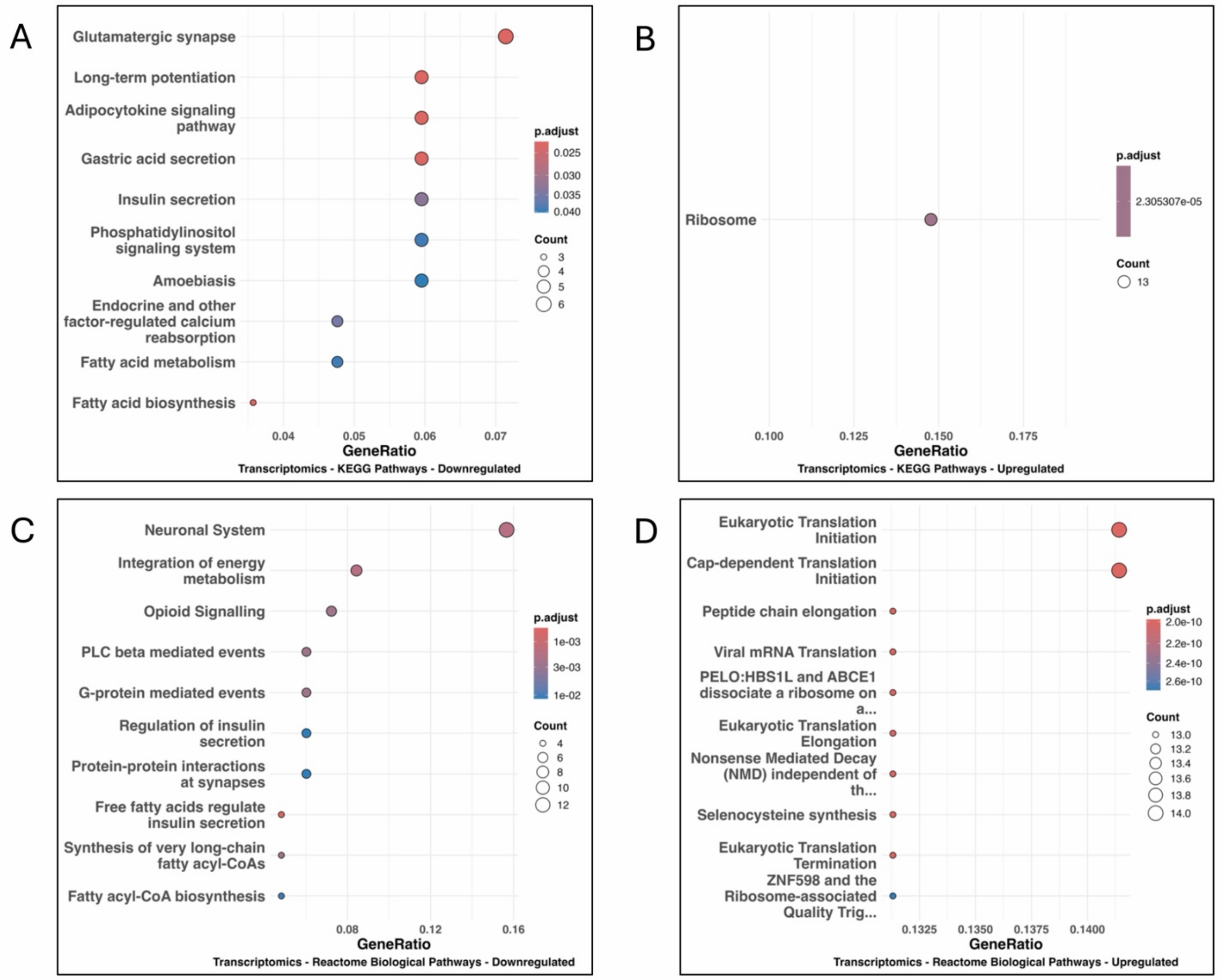


**Figure S14. KEGG and Reactome pathway enrichment analysis of transcriptomic changes in male T2 Cases vs. Controls.** KEGG pathway enrichment of downregulated (**A**) and upregulated (**B**) genes. Downregulated genes were enriched for neuronal and membrane-associated pathways, including glutamatergic synapse, long-term potentiation, phosphatidylinositol signaling, and lipid metabolism. In contrast, upregulated genes showed enrichment primarily for the ribosome pathway. Reactome pathway enrichment of downregulated (**C**) and upregulated (**D**) genes. Downregulated genes were enriched for neuronal system and G protein–mediated signaling pathways, whereas upregulated genes were enriched for eukaryotic translation, including initiation and elongation processes. Dot plots show enriched terms ranked by gene ratio, with point size indicating gene count and color representing adjusted P value.

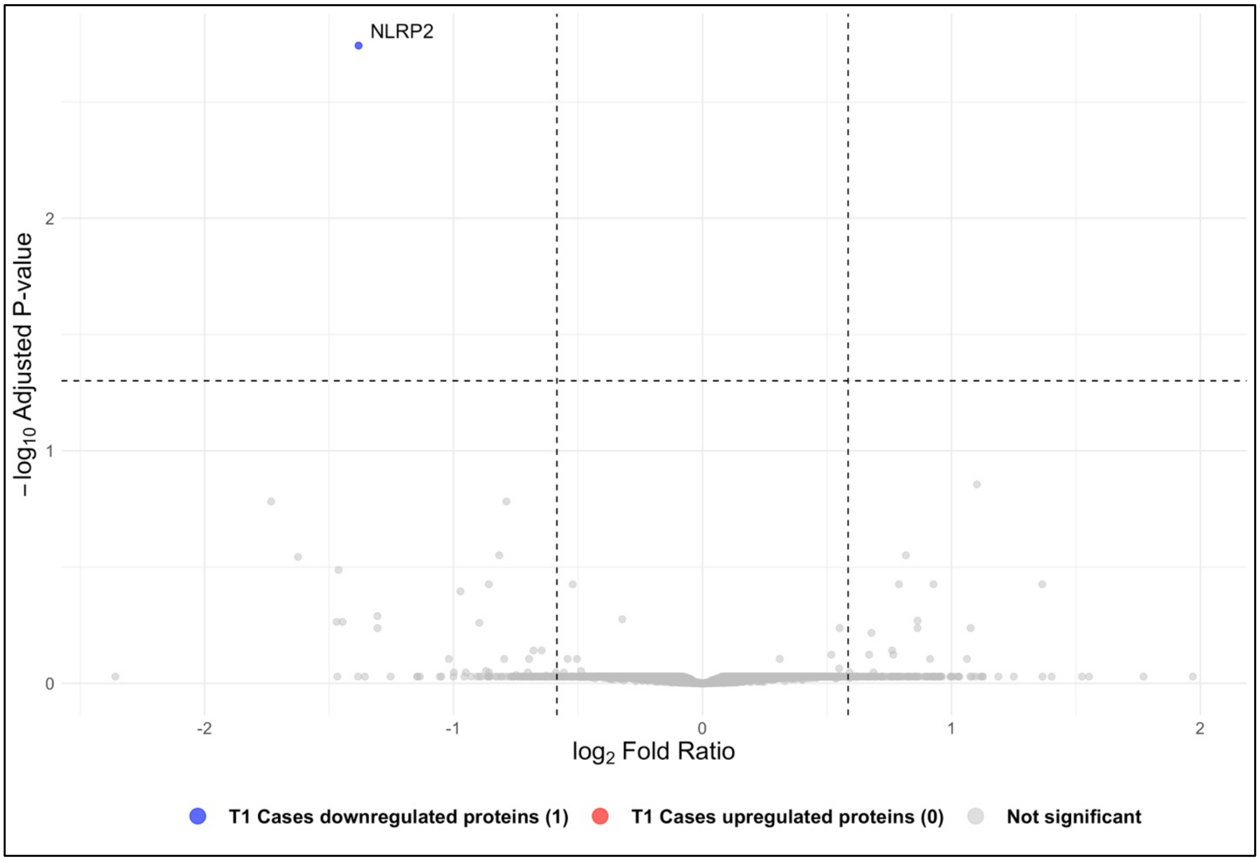


**Figure S15. Proteomic comparison of prenatal cannabis exposure of female T1 Cases vs. Controls.** Proteomic comparison of T1 Cases vs. Controls showing proteins associated with prenatal cannabis exposure. Volcano plot showing quantified proteins plotted by $\log_2$ fold ratio and Benjamini-Hochberg adjusted P value. Dashed vertical lines indicate the fold-ratio threshold ($|\log_2 FR| \geq 0.58$), and the horizontal dashed line indicates the adjusted P-value cutoff (adjusted P $\leq 0.05$). A single DEP, NLRP2, met the predefined significance criteria and was downregulated in T1 Cases relative to T1 Controls.

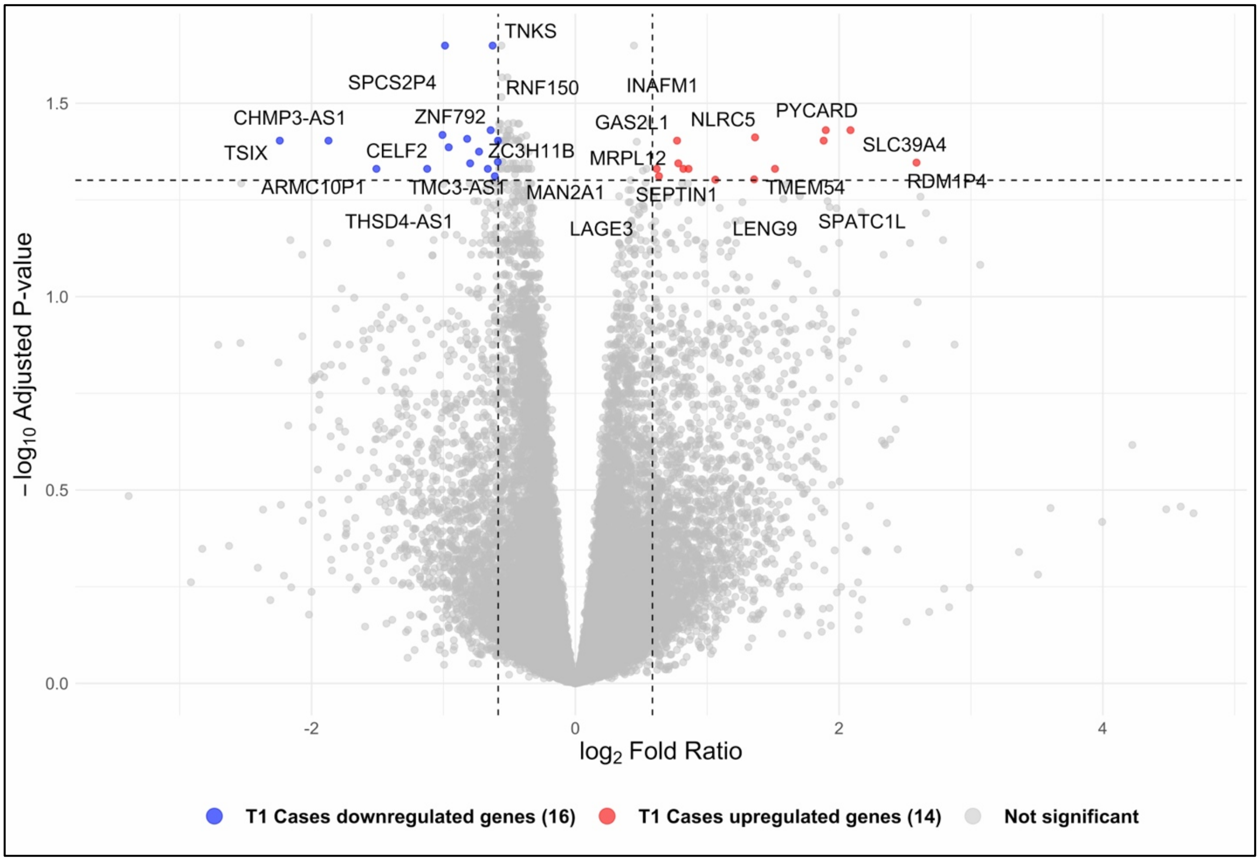


**Figure S16. Transcriptomic comparison of prenatal cannabis exposure in T1 fetal brains Cases vs. Controls (males and females).** Transcriptomic comparison of T1 Cases vs. Controls showing genes associated with prenatal cannabis exposure. Volcano plot showing quantified genes plotted by $log_2$ fold ratio and Benjamini-Hochberg adjusted P value. Dashed vertical lines indicate the fold-ratio threshold ($|log_2FR| \geq 0.58$), and the horizontal dashed line indicates the adjusted P-value cutoff (adjusted $P \leq 0.05$). A total of 30 DEGs met the predefined significance criteria, with both upregulated and downregulated genes observed in T1 Cases relative to T1 Controls.

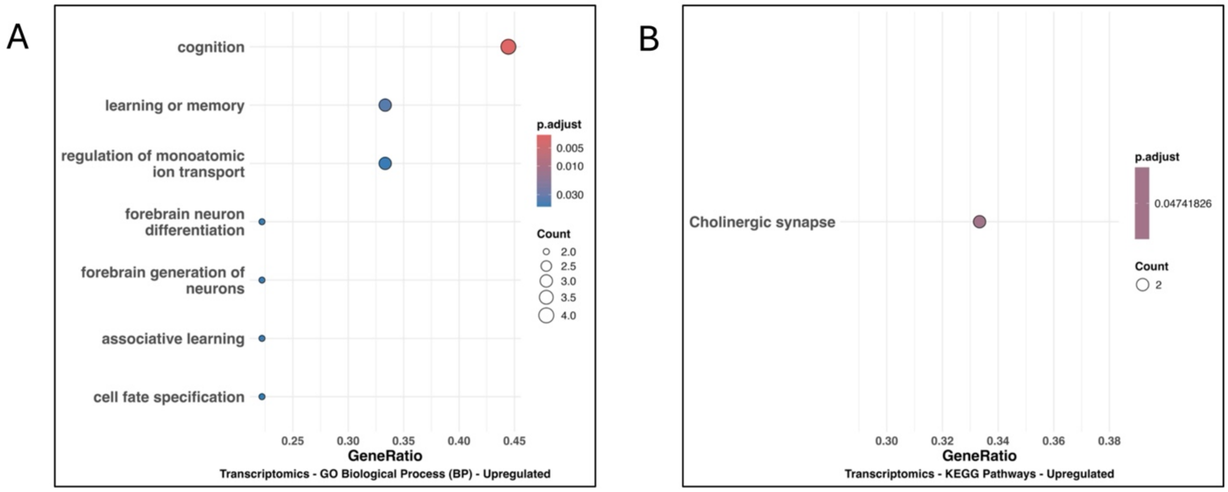


**Figure S17. Transcriptomic enrichment associated with development of control female fetal brains (T2 vs. T1 Controls). (A)** Gene Ontology (Biological Process) enrichment analysis of upregulated genes. Dot plot shows enriched terms ranked by gene ratio, with point size indicating gene count and color representing adjusted P value. Upregulated genes were enriched for processes related to cognition, learning and memory, ion transport regulation, and neuronal differentiation. **(B)** KEGG pathway enrichment analysis of upregulated genes. The cholinergic synapse pathway was enriched, indicating increased neurotransmission-related signaling during development.  Dot plots show enriched terms ranked by gene ratio, with point size indicating gene count and color representing adjusted P value.

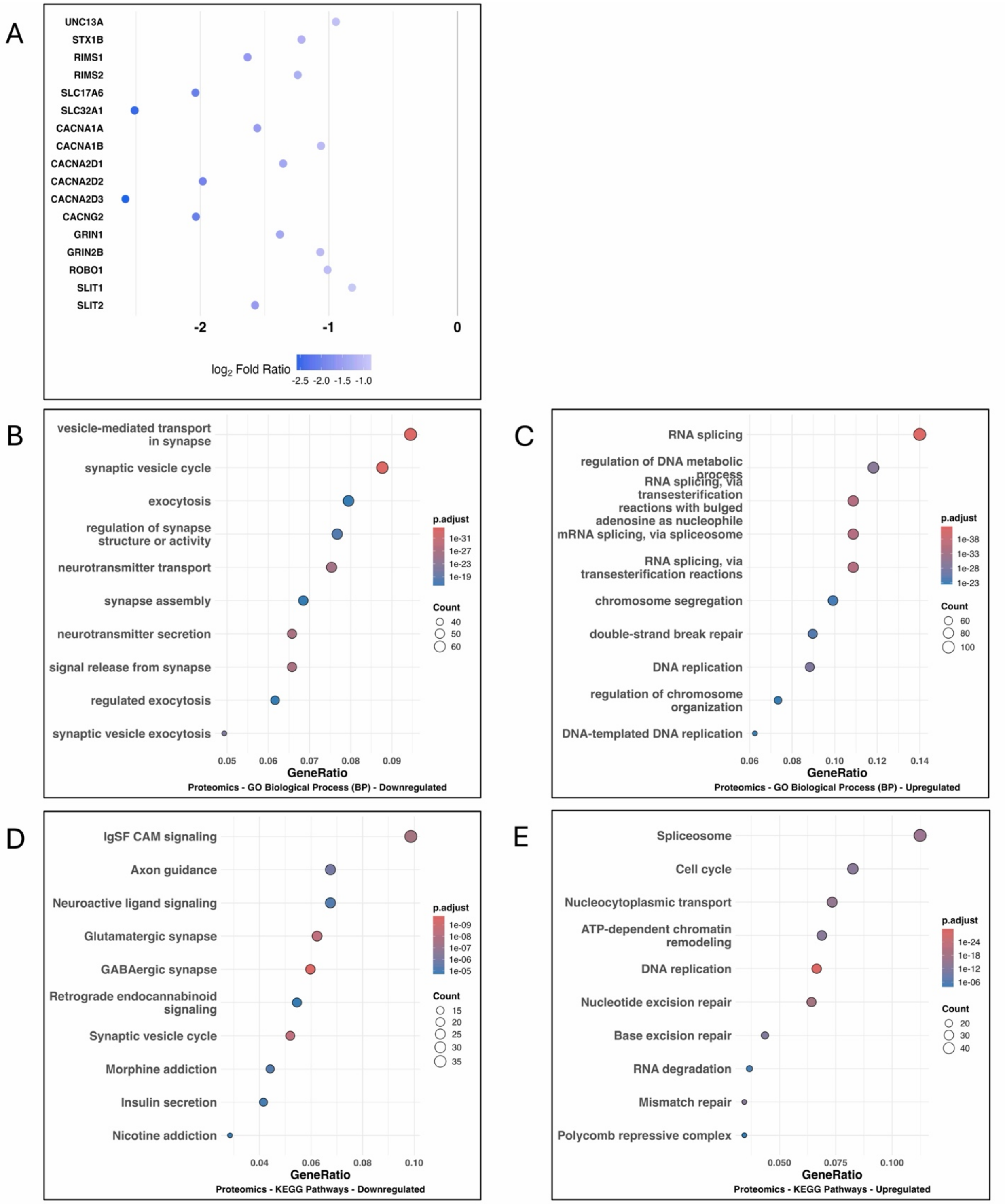


**Figure S18. Proteomic remodeling associated with control female fetal brain development (T2 vs. T1 Controls).** (**A**) Selected synaptic and excitability-related proteins showing reduced abundance in T2 relative to T1. Points represent $log_2$ fold ratio for individual proteins. Gene Ontology (Biological Process) enrichment analysis of downregulated (**B**) and upregulated (**C**)

proteins. Dot plots show enriched terms ranked by gene ratio, with point size indicating protein count and color representing adjusted P value. Downregulated proteins were enriched for synapse-related processes, whereas upregulated proteins were enriched for RNA processing, transcriptional regulation, and cell cycle activity. KEGG pathway enrichment analysis of downregulated (**D**) and upregulated (**E**) proteins. Downregulated pathways include neuronal signaling and synaptic transmission, whereas upregulated pathways were enriched for spliceosome, chromatin remodeling, and cell cycle–associated programs.

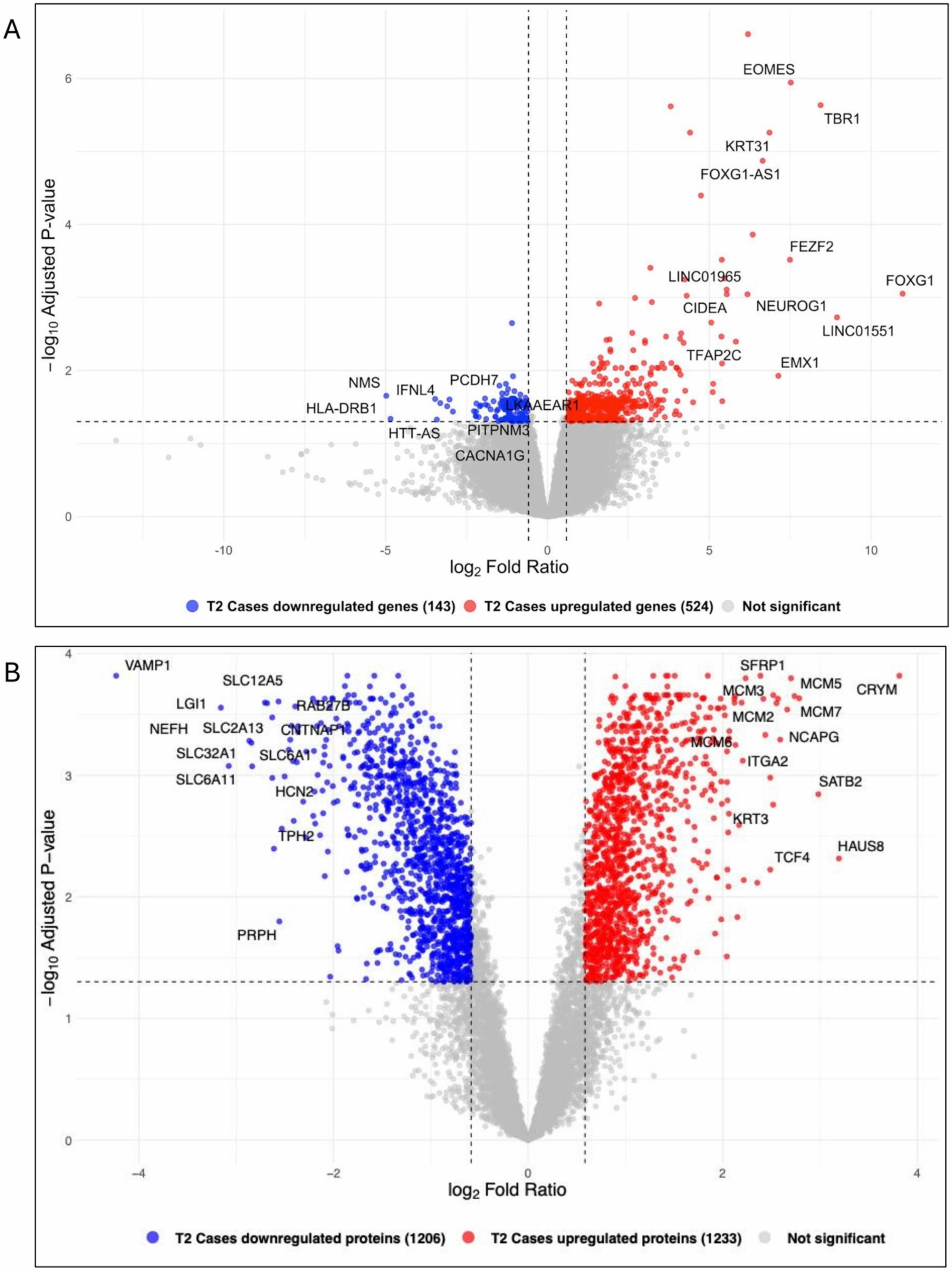

A
EOMES
TBR1
KRT31
FOXG1-AS1
FEZF2
LINC01965
FOXG1
CIDEA
NEUROG1
LINC01551
TFAP2C
EMX1
NMS
IFNL4
PCDH7
HLA-DRB1
LKAAEAR1
HTT-AS
PITPNM3
CACNA1G
−log10 Adjusted P-value
log2 Fold Ratio
T2 Cases downregulated genes (143)
T2 Cases upregulated genes (524)
Not significant
B
VAMP1
SLC12A5
LGI1
RAB27B
NEFH
SLC2A13
CNTNAP1
SLC32A1
SLC6A1
SLC6A11
HCN2
TPH2
PRPH
SFRP1
MCM5
CRYM
MCM3
MCM7
MCM2
NCAPG
MCM6
ITGA2
SATB2
KRT3
HAUS8
TCF4
−log10 Adjusted P-value
log2 Fold Ratio
T2 Cases downregulated proteins (1206)
T2 Cases upregulated proteins (1233)
Not significant

**Figure S19. Transcriptomic and proteomic association with brain development in male fetuses (T2 vs. T1 Cases).** Transcriptomic (**A**) and proteomic (**B**) comparisons of T2 versus T1 case fetal brains showing gene products associated with development. Volcano plots showing quantified genes or proteins plotted by $\log_2$ fold ratio and Benjamini-Hochberg adjusted P value. Dashed vertical lines indicate the fold-ratio threshold ($|\log_2 FR| \geq 0.58$), and the horizontal dashed line indicates the adjusted P-value cutoff (adjusted $P \leq 0.05$). In **A**, 667 DEGs were identified, with prominent upregulation of neurodevelopmental regulators including TBR1, EOMES, and FOXG1 in T2 relative to T1. In **B**, 2,439 DEPs were identified, indicating extensive proteomic remodeling during development in Cases.

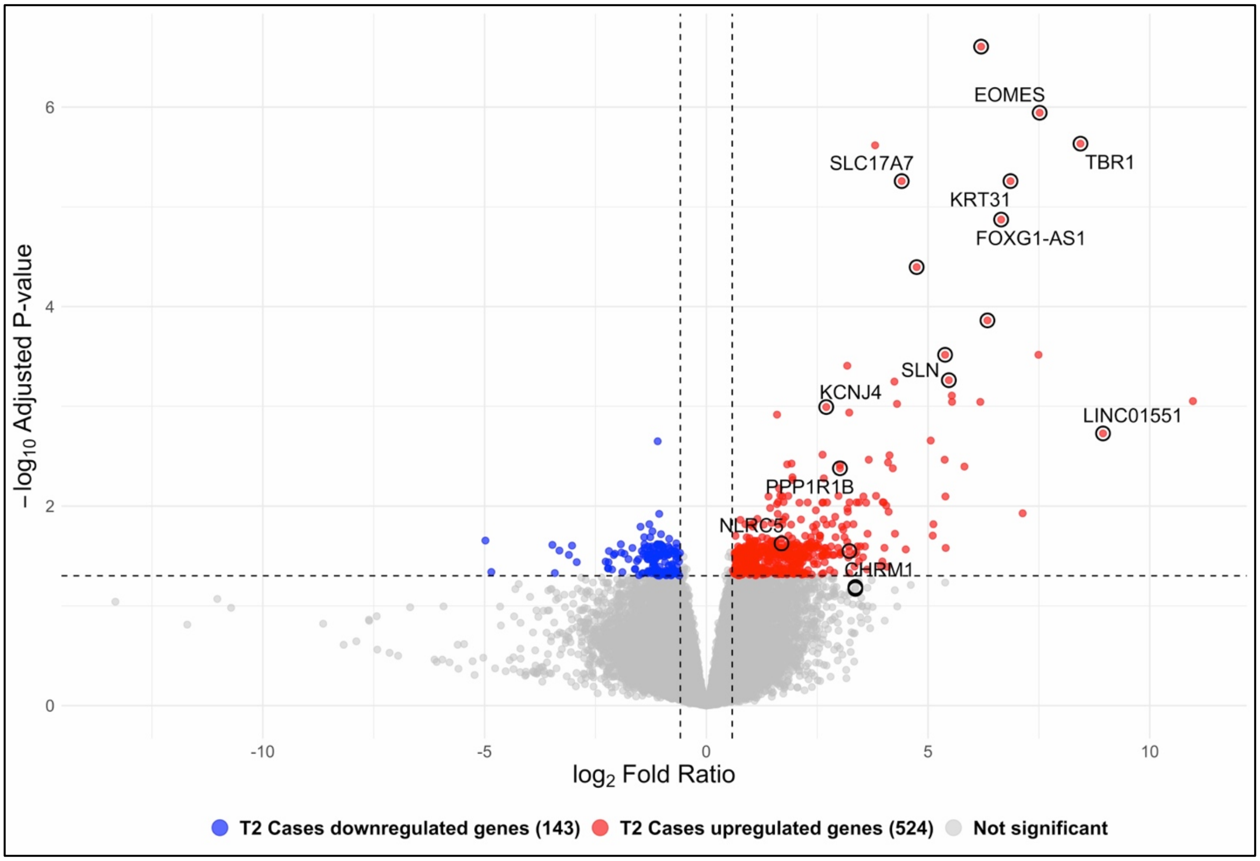


**Figure S20. Transcriptomic association with brain development in male fetuses (T2 vs. T1 Cases) with female control development overlaid.** Transcriptomic comparison of T2 versus T1 case brains showing genes associated with developmental progression. Volcano plot showing quantified genes plotted by $\log_2$ fold ratio and Benjamini-Hochberg adjusted P value. Dashed vertical lines indicate the fold-ratio threshold ($|\log_2 FR| \geq 0.58$), and the horizontal dashed line indicates the adjusted P-value cutoff (adjusted $P \leq 0.05$). Genes defining female control development (T2 Controls versus T1 Controls) are overlaid and highlighted by black circles. Red and blue points indicate genes significantly upregulated in T2 and T1 case samples, respectively.

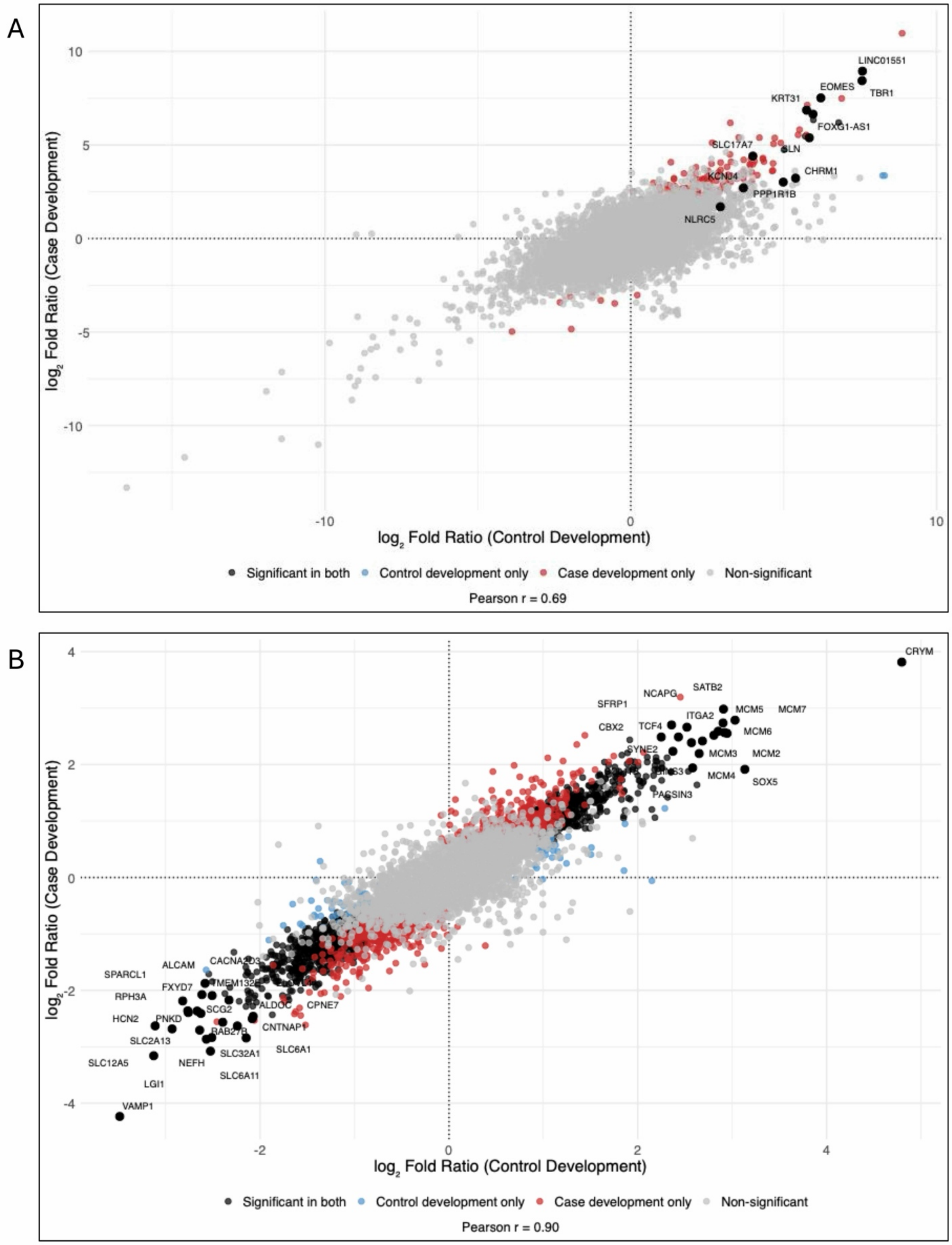

A
LINC01551
EOMES
TBR1
KRT31
FOXG1-AS1
SLC17A7
SLN
KCNJ4
CHRM1
PPP1R1B
NLRC5
10
5
0
-5
-10
-10
0
10
log2 Fold Ratio (Case Development)
log2 Fold Ratio (Control Development)
Significant in both
Control development only
Case development only
Non-significant
Pearson r = 0.69
B
CRYM
NCAPG
SATB2
SFRP1
MCM5
MCM7
ITGA2
CBX2
TCF4
MCM6
SYNE2
MCM3
MCM2
MCM4
SOX5
PACSIN3
ALCAM
CACNA2D3
SPARCL1
FXYD7
TMEM132E
RPH3A
ALDOC
CPNE7
SCG2
HCN2
PNKD
CNTNAP1
RAB27B
SLC2A13
SLC6A1
SLC32A1
SLC12A5
NEFH
SLC6A11
LGI1
VAMP1
4
2
0
-2
-4
-2
0
2
4
log2 Fold Ratio (Case Development)
log2 Fold Ratio (Control Development)
Significant in both
Control development only
Case development only
Non-significant
Pearson r = 0.90

**Figure S21. Cross-development concordance of transcriptomic and proteomic data between control females and case males.** (**A**) Transcriptomic concordance plot comparing $log_2$ fold ratio between control female development (T2 versus T1) and case male development (T2 versus T1). Each point represents a gene quantified in both datasets and is classified as significant in both conditions, control-only, case-only, or non-significant. A positive correlation (Pearson r = 0.69) indicates shared directionality of transcriptional changes across conditions. (**B**) Proteomic concordance plot comparing $log_2$ fold ratios between control female development (T2 versus T1) and case male development (T2 versus T1). Each point represents a protein quantified in both datasets and is classified as above. A stronger positive correlation (Pearson r = 0.90) indicated highly consistent directional change at the proteome level.

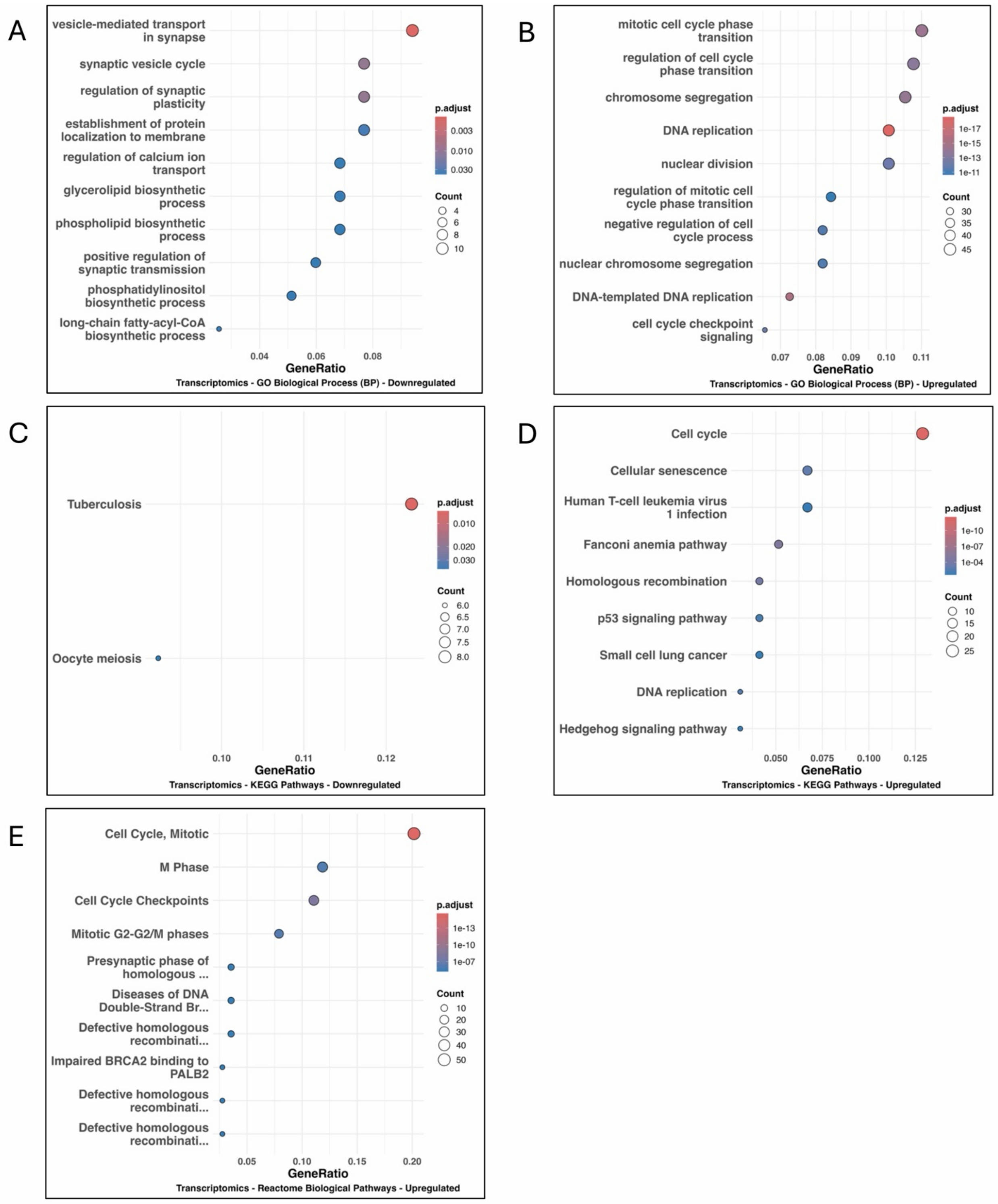


**Figure S22. Transcriptomic enrichment associated with development in male T2 Cases vs. T1 Cases.** Gene Ontology (Biological Process) enrichment analysis of downregulated (**A**) and upregulated (**B**) genes. Dot plots show enriched terms ranked by gene ratio, with point size indicating gene count and color representing adjusted P value. Downregulated genes were enriched for synaptic vesicle cycling, synaptic transmission, and lipid metabolic processes, whereas upregulated genes were enriched for cell cycle progression, DNA replication, and nuclear division. KEGG pathway enrichment analysis of downregulated (**C**) and upregulated (**D**) genes. Downregulated pathways were limited and do not form a coherent program, whereas upregulated pathways were dominated by cell cycle and DNA replication processes. Reactome pathway enrichment analysis of upregulated (**E**) genes. Upregulated pathways were enriched for mitotic cell cycle, DNA repair, and homologous recombination processes.

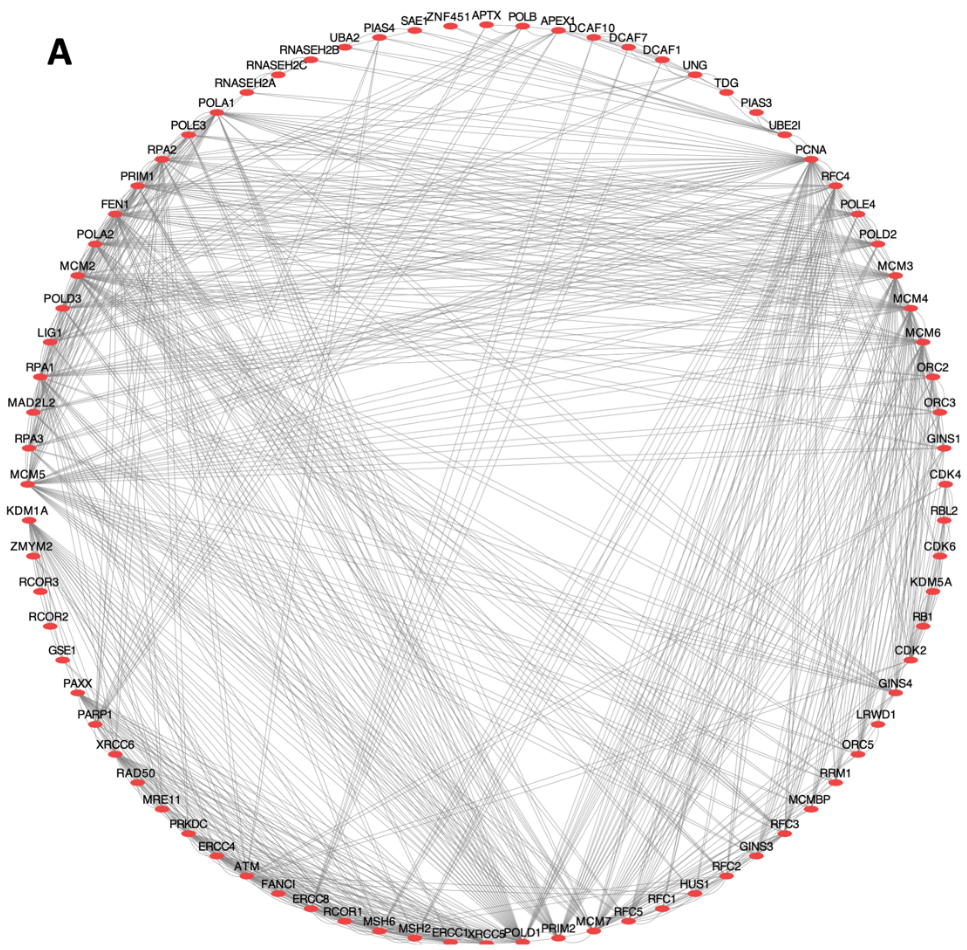
A
SAE1
ZNF451
APTX
POLB
APEX1
DCAF10
DCAF7
DCAF1
UNG
TDG
PIAS3
UBE2I
PCNA
RFC4
POLE4
POLD2
MCM3
MCM4
MCM6
ORC2
ORC3
GINS1
CDK4
RBL2
CDK6
KDM5A
RB1
CDK2
GINS4
LRWD1
ORC5
RRM1
MCMBP
RFC3
GINS3
RFC2
HUS1
RFC1
RFC5
MCM7
PRIM2
POLD1
XRCC5
ERCC1
MSH2
MSH6
RCOR1
ERCC8
FANCI
ATM
ERCC4
PRKDC
MRE11
RAD50
XRCC6
PARP1
PAXX
GSE1
RCOR2
RCOR3
ZMYM2
KDM1A
MCM5
RPA3
MAD2L2
RPA1
LIG1
POLD3
MCM2
POLA2
FEN1
PRIM1
RPA2
POLE3
POLA1
RNASEH2A
RNASEH2C
RNASEH2B
UBA2
PIAS4

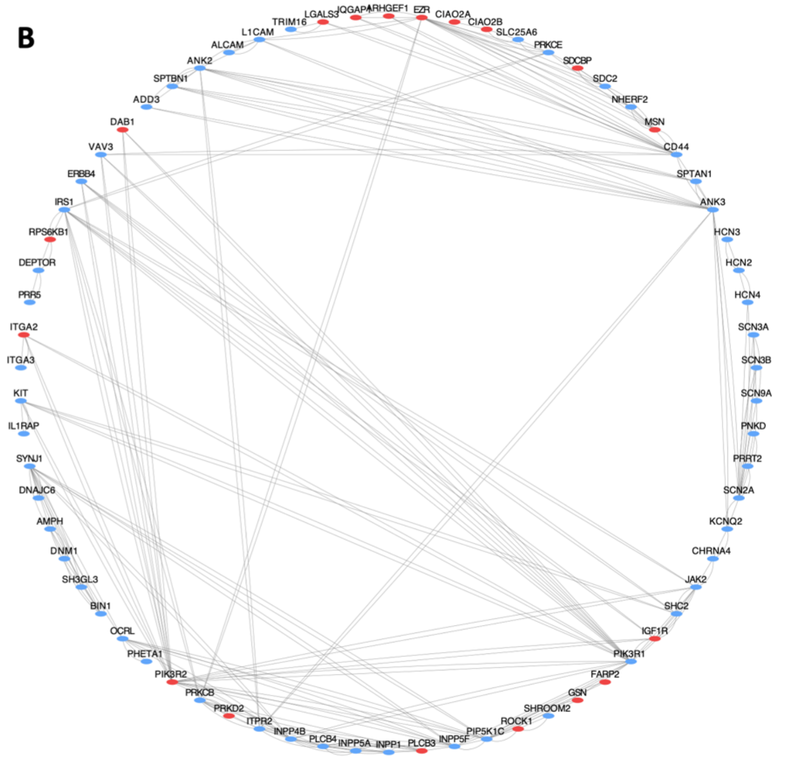
B
TRIM16
LGALS3
IQGAP1
ARHGEF1
EZR
CIAO2A
CIAO2B
SLC25A6
PRKCE
SDCBP
SDC2
NHERF2
MSN
CD44
SPTAN1
ANK3
HCN3
HCN2
HCN4
SCN3A
SCN3B
SCN9A
PNKD
PRRT2
SCN2A
KCNQ2
CHRNA4
JAK2
SHC2
IGF1R
PIK3R1
FARP2
GSN
SHROOM2
ROCK1
PIP5K1C
INPP5F
PLCB3
INPP1
INPP5A
PLCB4
INPP4B
ITPR2
PRKD2
PRKCB
PIK3R2
PHETA1
OCRL
BIN1
SH3GL3
DNM1
AMPH
DNAJC6
SYNJ1
IL1RAP
KIT
ITGA3
ITGA2
PRR5
DEPTOR
RPS6KB1
IRS1
ERBB4
VAV3
DAB1
ADD3
SPTBN1
ANK2
ALCAM
L1CAM

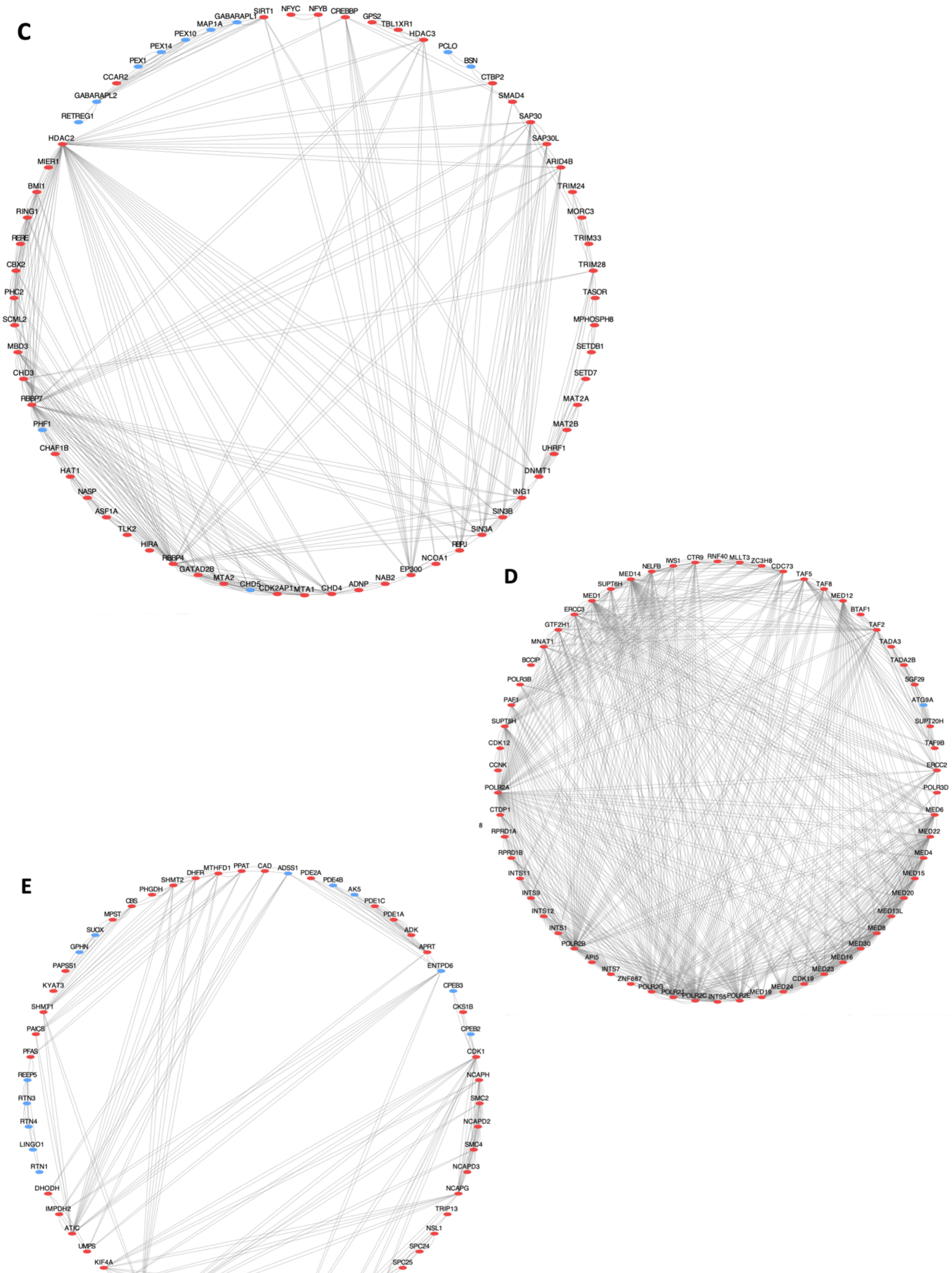
C
SIRT1
NFYC
NFYB
CREBBP
GPS2
TBL1XR1
HDAC3
PCLO
BSN
CTBP2
SMAD4
SAP30
SAP30L
ARID4B
TRIM24
MORC3
TRIM33
TRIM28
TASOR
MPHOSPH8
SETDB1
SETD7
MAT2A
MAT2B
UHRF1
DNMT1
ING1
SIN3B
SIN3A
RBPJ
NCOA1
EP300
NAB2
ADNP
CHD4
MTA1
CDK2AP1
CHD5
MTA2
GATAD2B
RBBP4
HIRA
TLK2
ASF1A
NASP
HAT1
CHAF1B
PHF1
RBBP7
CHD3
MBD3
SCML2
PHC2
CBX2
RERE
RING1
BMI1
MIER1
HDAC2
RETREG1
GABARAPL2
CCAR2
PEX1
PEX14
PEX10
MAP1A
GABARAPL1
D
IWS1
CTR9
RNF40
MLLT3
ZC3H8
CDC73
TAF5
TAF8
MED12
BTAF1
TAF2
TADA3
TADA2B
SGF29
ATG9A
SUPT20H
TAF9B
ERCC2
POLR3D
MED6
MED22
MED4
MED15
MED20
MED13L
MED8
MED30
MED16
MED23
CDK19
MED24
MED19
POLR2E
INTS5
POLR2C
POLR2J
POLR2G
ZNF687
INTS7
API5
POLR2B
INTS1
INTS12
INTS9
INTS11
RPRD1B
RPRD1A
8
CTDP1
POLR2A
CCNK
CDK12
SUPT5H
PAF1
POLR3B
BCCIP
MNAT1
GTF2H1
ERCC3
MED1
SUPT6H
MED14
NELFB
E
CAD
ADSS1
PDE2A
PDE4B
AK5
PDE1C
PDE1A
ADK
APRT
ENTPD6
CPEB3
CKS1B
CPEB2
CDK1
NCAPH
SMC2
NCAPD2
SMC4
NCAPD3
NCAPG
TRIP13
NSL1
SPC24
SPC25
BUB3
MAD2L1
PBK
RAVER1
RRM2
DCTPP1
NME7
NME4
DTYMK
DCTD
KIF11
TYMS
MKI67
KIF4A
UMPS
ATIC
IMPDH2
DHODH
RTN1
LINGO1
RTN4
RTN3
REEP5
PFAS
PAICS
SHMT1
KYAT3
PAPSS1
GPHN
SUOX
MPST
CBS
PHGDH
SHMT2
DHFR
MTHFD1
PPAT

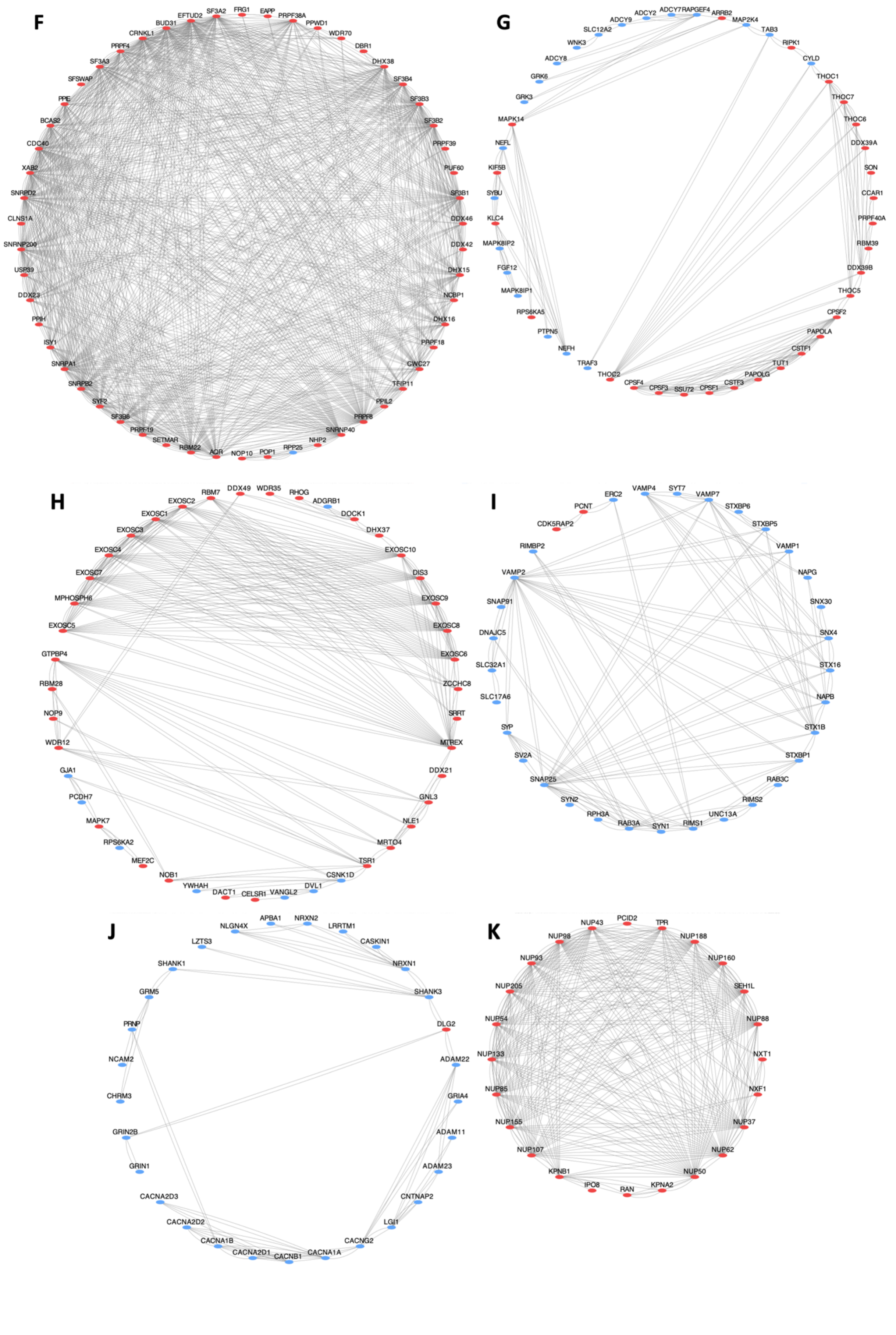
F
EFTUD2
SF3A2
FRG1
EAPP
PRPF38A
PPWD1
BUD31
WDR70
CRNKL1
DBR1
PRPF4
DHX38
SF3A3
SF3B4
SFSWAP
SF3B3
PPIE
SF3B2
BCAS2
PRPF39
CDC40
PUF60
XAB2
SF3B1
SNRPD2
DDX46
CLNS1A
DDX42
SNRNP200
DHX15
USP39
NCBP1
DDX23
DHX16
PPIH
PRPF18
ISY1
CWC27
SNRPA1
TFIP11
SNRPB2
PPIL2
SYF2
PRPF8
SF3B6
SNRNP40
PRPF19
NHP2
SETMAR
RPP25
RBM22
POP1
AQR
NOP10
G
ADCY2
ADCY7
RAPGEF4
ARRB2
ADCY9
MAP2K4
SLC12A2
TAB3
WNK3
RIPK1
ADCY8
CYLD
GRK6
THOC1
GRK3
THOC7
MAPK14
THOC6
NEFL
DDX39A
KIF5B
SON
SYBU
CCAR1
KLC4
PRPF40A
MAPK8IP2
RBM39
FGF12
DDX39B
MAPK8IP1
THOC5
RPS6KA5
CPSF2
PTPN5
PAPOLA
NEFH
CSTF1
TRAF3
TUT1
THOC2
PAPOLG
CPSF4
CSTF3
CPSF3
SSU72
CPSF1
H
RBM7
DDX49
WDR35
RHOG
EXOSC2
ADGRB1
EXOSC1
DOCK1
EXOSC3
DHX37
EXOSC4
EXOSC10
EXOSC7
DIS3
MPHOSPH6
EXOSC9
EXOSC5
EXOSC8
GTPBP4
EXOSC6
RBM28
ZCCHC8
NOP9
SRRT
WDR12
MTREX
GJA1
DDX21
PCDH7
GNL3
MAPK7
NLE1
RPS6KA2
MRTO4
MEF2C
TSR1
NOB1
CSNK1D
YWHAH
DVL1
DACT1
VANGL2
CELSR1
I
VAMP4
SYT7
ERC2
VAMP7
PCNT
STXBP6
CDK5RAP2
STXBP5
RIMBP2
VAMP1
VAMP2
NAPG
SNAP91
SNX30
DNAJC5
SNX4
SLC32A1
STX16
SLC17A6
NAPB
SYP
STX1B
SV2A
STXBP1
SNAP25
RAB3C
SYN2
RIMS2
RPH3A
UNC13A
RAB3A
SYN1
RIMS1
J
APBA1
NRXN2
NLGN4X
LRRTM1
LZTS3
CASKIN1
SHANK1
NRXN1
GRM5
SHANK3
PRNP
DLG2
NCAM2
ADAM22
CHRM3
GRIA4
GRIN2B
ADAM11
GRIN1
ADAM23
CACNA2D3
CNTNAP2
CACNA2D2
LGI1
CACNA1B
CACNG2
CACNA2D1
CACNA1A
CACNB1
K
NUP43
PCID2
TPR
NUP98
NUP188
NUP93
NUP160
NUP205
SEH1L
NUP54
NUP88
NUP133
NXT1
NUP85
NXF1
NUP155
NUP37
NUP107
NUP62
KPNB1
NUP50
IPO8
KPNA2
RAN

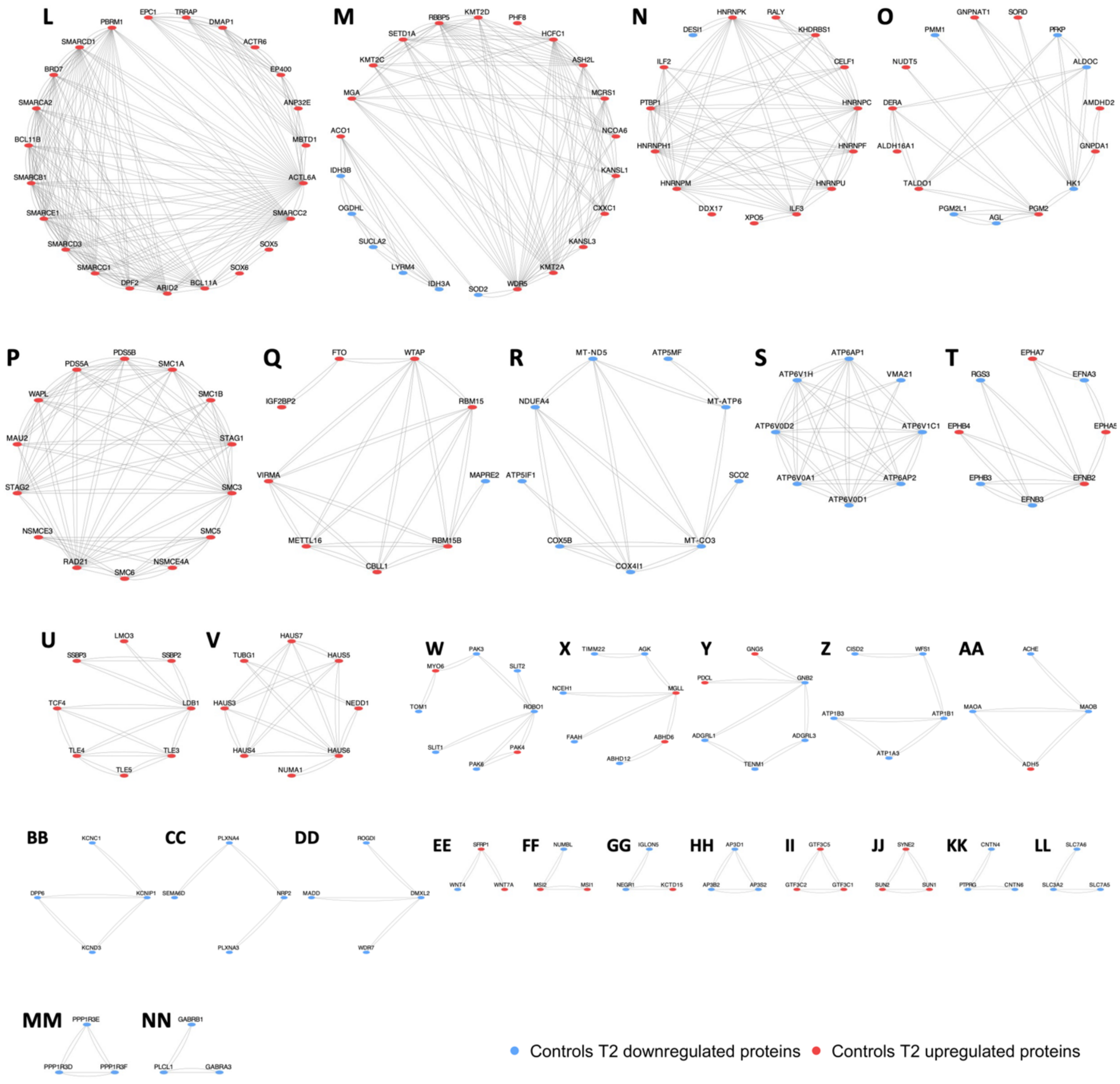


| Cluster | STRING Interaction Module Enriched For | Upregulated Proteins | Downregulated Proteins |
|---|---|---|---|
| A | DNA Replication Machinery and Genome Maintenance | APEX1, APTX, ATM, CDK2, CDK4, CDK6, DCAF1, DCAF10, DCAF7, ERCC1, ERCC4, ERCC8, FANCI, FEN1, GINS1, GINS3, GINS4, GSE1, HUS1, KDM1A, KDM5A, LIG1, LRWD1, MAD2L2, MCM2, MCM3, MCM4, MCM5, MCM6, MCM7, MCMBP, MRE11, MSH2, MSH6, ORC2, ORC3, ORC5, PARP1, PAXX, PCNA, PIAS3, PIAS4, POLA1, POLA2, POLB, POLD1, POLD2, POLD3, POLE3, POLE4, PRIM1, PRIM2, PRKDC, RAD50, RB1, RBL2, RCOR1, RCOR2, RCOR3, RFC1, RFC2, RFC3, RFC4, RFC5, RNASEH2A, RNASEH2B, RNASEH2C, RPA1, RPA2, RPA3, RRM1, SAE1, TDG, UBA2, UBE2I, UNG, XRCC5, XRCC6, ZMYM2, ZNF451 | |
| AA | Monoamine Oxidation and Neurotransmitter Catabolism | ADH5 | ACHE, MAOA, MAOB |
| B | PI3K Signaling and Neuronal Ion Channel Regulation | ARHGEF1, CIAO2A, CIAO2B, DAB1, EZR, FARP2, GSN, IGF1R, IQGAP1, ITGA2, LGALS3, MSN, PIK3R2, PLCB3, PRKD2, ROCK1, RPS6KB1, SDCBP | ADD3, ALCAM, AMPH, ANK2, ANK3, BIN1, CD44, CHRNA4, DEPTOR, DNAJC6, DNM1, ERBB4, HCN2, HCN3, HCN4, IL1RAP, INPP1, INPP4B, INPP5A, INPP5F, IRS1, ITGA3, ITPR2, JAK2, KCNQ2, KIT, L1CAM, NHERF2, OCRL, PHETA1, PIK3R1, PIP5K1C, PLCB4, PNKD, PRKCB, PRKCE, PRR5, PRRT2, SCN2A, SCN3A, SCN3B, SCN9A, SDC2, SH3GL3, SHC2, SHROOM2, SLC25A6, SPTAN1, SPTBN1, SYNJ1, TRIM16, VAV3 |
| BB | Kv4 Potassium Channel Complex and Neuronal Excitability | | DPP6, KCNC1, KCND3, KCNIP1 |

| | | | |
|---|---|---|---|
| C | Chromatin Remodeling Complexes and Histone Deacetylation | ADNP, ARID4B, ASF1A, BMI1, CBX2, CCAR2, CDK2AP1, CHAF1B, CHD3, CHD4, CREBBP, CTBP2, DNMT1, EP300, GATAD2B, GPS2, HAT1, HDAC2, HDAC3, HIRA, ING1, MAT2A, MAT2B, MBD3, MIER1, MORC3, MPHOSPH8, MTA1, MTA2, NAB2, NASP, NCOA1, NFYB, NFYC, PHC2, RBBP4, RBBP7, RBPJ, RERE, RING1, SAP30, SAP30L, SCML2, SETD7, SETDB1, SIN3A, SIN3B, SIRT1, SMAD4, TASOR, TBL1XR1, TLK2, TRIM24, TRIM28, TRIM33, UHRF1 | BSN, CHD5, GABARAPL1, GABARAPL2, MAP1A, PCLO, PEX1, PEX10, PEX14, PHF1, RETREG1 |
| CC | Semaphorin–Plexin Signaling and Axon Guidance | | NRP2, PLXNA3, PLXNA4, SEMA6D |
| D | RNA Polymerase II Transcription Machinery and Mediator Regulation | API5, BCCIP, BTAF1, CCNK, CDC73, CDK12, CDK19, CTDP1, CTR9, ERCC2, ERCC3, GTF2H1, INTS1, INTS11, INTS12, INTS5, INTS7, INTS9, IWS1, MED1, MED12, MED13L, MED14, MED15, MED16, MED19, MED20, MED22, MED23, MED24, MED30, MED4, MED6, MED8, MLLT3, MNAT1, NELFB, PAF1, POLR2A, POLR2B, POLR2C, POLR2E, POLR2G, POLR2J, POLR3B, POLR3D, RNF40, RPRD1A, RPRD1B, SGF29, SUPT20H, SUPT5H, SUPT6H, TADA2B, TADA3, TAF2, TAF5, TAF8, TAF9B, ZC3H8, ZNF687 | ATG9A |
| DD | Rab3/Rabconnectin Vesicle Trafficking and Synaptic Regulation | | DMXL2, MADD, ROGDI, WDR7 |
| E | Cell Cycle Progression and Nucleotide Biosynthesis | ADK, APRT, ATIC, BUB3, CAD, CBS, CDK1, CKS1B, DCTD, DCTPP1, DHFR, DHODH, DTYMK, IMPDH2, KIF11, KIF4A, KYAT3, MAD2L1, MKI67, MPST, MTHFD1, NCAPD2, NCAPD3, NCAPG, NCAPH, NME4, NME7, NSL1, PAICS, PAPSS1, PBK, PDE1A, PDE1C, PDE2A, PFAS, PHGDH, PPAT, RAVER1, RRM2, SHMT1, SHMT2, SMC2, SMC4, SPC24, SPC25, TRIP13, TYMS, UMPS | ADSS1, AK5, CPEB2, CPEB3, ENTPD6, GPHN, LINGO1, PDE4B, REEP5, RTN1, RTN3, RTN4, SUOX |
| EE | WNT Ligand Signaling and Developmental Patterning | SFRP1, WNT7A | WNT4 |
| F | Spliceosomal snRNP Assembly and Pre-mRNA Splicing | AQR, BCAS2, BUD31, CDC40, CLNS1A, CRNKL1, CWC27, DBR1, DDX23, DDX42, DDX46, DHX15, DHX16, DHX38, EAPP, EFTUD2, FRG1, ISY1, NCBP1, NHP2, NOP10, POP1, PPIE, PPIH, PPIL2, PPWD1, PRPF18, PRPF19, PRPF38A, PRPF39, PRPF4, PRPF8, PUF60, RBM22, SETMAR, SF3A2, SF3A3, SF3B1, SF3B2, SF3B3, SF3B4, SF3B6, SFSWAP, SNRNP200, SNRNP40, SNRPA1, SNRPB2, SNRPD2, SYF2, TFIP11, USP39, WDR70, XAB2 | RPP25 |
| FF | Neural Stem Cell RNA Binding and Notch Fate Regulation | MSI1, MSI2 | NUMBL |
| G | mRNA Cleavage and Polyadenylation with MAPK Signaling | ARRB2, CCAR1, CPSF1, CPSF2, CPSF3, CPSF4, CSTF1, CSTF3, DDX39A, DDX39B, KIF5B, KLC4, MAPK14, PAPOLA, PAPOLG, PRPF40A, RBM39, RIPK1, RPS6KA5, SON, SSU72, THOC1, THOC2, THOC5, THOC6, THOC7, TUT1 | ADCY2, ADCY7, ADCY8, ADCY9, CYLD, FGF12, GRK3, GRK6, MAP2K4, MAPK8IP1, MAPK8IP2, NEFH, NEFL, PTPN5, RAPGEF4, SLC12A2, SYBU, TAB3, TRAF3, WNK3 |
| GG | IgLON Adhesion Molecules and Neuronal Connectivity | KCTD15 | IGLON5, NEGR1 |
| H | RNA Exosome Surveillance and Ribosomal Biogenesis | CELSR1, DACT1, DDX21, DDX49, DHX37, DIS3, DOCK1, EXOSC1, EXOSC10, EXOSC2, EXOSC3, EXOSC4, EXOSC5, EXOSC6, EXOSC7, EXOSC8, EXOSC9, GNL3, GTPBP4, MAPK7, MEF2C, MPHOSPH6, MRTO4, MTREX, NLE1, NOB1, NOP9, RBM28, RBM7, RHOG, SRRT, TSR1, WDR12, WDR35, ZCCHC8 | ADGRB1, CSNK1D, DVL1, GJA1, PCDH7, RPS6KA2, VANGL2, YWHAH |
| HH | AP-3 Vesicle Trafficking Complex and Synaptic Transport | | AP3B2, AP3D1, AP3S2 |
| I | Synaptic Vesicle Docking and Neurotransmitter Release | CDK5RAP2, PCNT | DNAJC5, ERC2, NAPB, NAPG, RAB3A, RAB3C, RIMBP2, RIMS1, RIMS2, RPH3A, SLC17A6, SLC32A1, SNAP25, SNAP91, SNX30, SNX4, STX16, STX1B, STXBP1, STXBP5, STXBP6, SV2A, SYN1, SYN2, SYP, SYT7, UNC13A, VAMP1, VAMP2, VAMP4, VAMP7 |
| II | RNA Polymerase III Transcription Complex Assembly | GTF3C1, GTF3C2, GTF3C5 | |
| J | Postsynaptic Receptor Scaffold and Glutamatergic Signaling | DLG2 | ADAM11, ADAM22, ADAM23, APBA1, CACNA1A, CACNA1B, CACNA2D1, CACNA2D2, CACNA2D3, CACNB1, CACNG2, CASKIN1, CHRM3, CNTNAP2, GRIA4, GRIN1, GRIN2B, GRM5, LGI1, LRRTM1, LZTS3, NCAM2, NLGN4X, NRXN1, NRXN2, PRNP, SHANK1, SHANK3 |
| JJ | LINC Nuclear Envelope Complex and Nuclear Positioning | SUN1, SUN2, SYNE2 | |
| K | Nuclear Pore Complex Assembly and Nucleocytoplasmic Transport | IPO8, KPNA2, KPNB1, NUP107, NUP133, NUP155, NUP160, NUP188, NUP205, NUP37, NUP43, NUP50, NUP54, NUP62, NUP85, NUP88, NUP93, NUP98, NXF1, NXT1, PCID2, RAN, SEH1L, TPR | |
| KK | Contactin Adhesion Complexes and Axon Guidance | | CNTN4, CNTN6, PTPRG |
| L | SWI/SNF Chromatin Remodeling and Transcriptional Regulation | ACTL6A, ACTR6, ANP32E, ARID2, BCL11A, BCL11B, BRD7, DMAP1, DPF2, EP400, EPC1, MBTD1, PBRM1, SMARCA2, SMARCB1, SMARCC1, SMARCC2, SMARCD1, SMARCD3, SMARCE1, SOX5, SOX6, TRRAP | |
| LL | Large Neutral Amino Acid Transport and mTOR Nutrient Signaling | | SLC3A2, SLC7A5, SLC7A6 |
| M | SET1/MLL Histone Methyltransferase Complex and Chromatin Activation | ACO1, ASH2L, CXXC1, HCFC1, KANSL1, KANSL3, KMT2A, KMT2C, KMT2D, MCRS1, MGA, NCOA6, PHF8, RBBP5, SETD1A, WDR5 | IDH3A, IDH3B, LYRM4, OGDHL, SOD2, SUCLA2 |
| MM | Protein Phosphatase-1 Glycogen Regulation and Metabolic Control | | PPP1R3D, PPP1R3E, PPP1R3F |
| N | Heterogeneous Nuclear RNP Complexes and RNA Processing | CELF1, DDX17, HNRNPC, HNRNPF, HNRNPH1, HNRNPK, HNRNPM, HNRNPU, ILF2, ILF3, KHDRBS1, PTBP1, RALY, XPO5 | DESI1 |
| NN | GABA Receptor Signaling and Inhibitory Synaptic Regulation | | GABRA3, GABRB1, PLCL1 |
| O | Glycolysis and Hexose Phosphate Metabolism | ALDH16A1, AMDHD2, DERA, GNPDA1, GNPNAT1, NUDT5, PGM2, SORD, TALDO1 | AGL, ALDOC, HK1, PFKP, PGM2L1, PMM1 |
| P | Cohesin Complex Assembly and Chromosome Segregation | MAU2, NSMCE3, NSMCE4A, PDS5A, PDS5B, RAD21, SMC1A, SMC1B, SMC3, SMC5, SMC6, STAG1, STAG2, WAPL | |
| Q | m6A RNA Methylation Machinery and Post-transcriptional Regulation | CBLL1, FTO, IGF2BP2, METTL16, RBM15, RBM15B, VIRMA, WTAP | MAPRE2 |
| R | Mitochondrial Electron Transport and Oxidative Phosphorylation | | ATP5IF1, ATP5MF, COX4I1, COX5B, MT-ATP6, MT-CO3, MT-ND5, NDUFA4, SCO2 |

| | | | |
|---|---|---|---|
| S | V-ATPase Proton Pump Assembly and Vesicular Acidification | | ATP6AP1, ATP6AP2, ATP6V0A1, ATP6V0D1, ATP6V0D2, ATP6V1C1, ATP6V1H, VMA21 |
| T | Ephrin–EPH Receptor Signaling and Axon Guidance | EFNB2, EPHA5, EPHA7, EPHB4 | EFNA3, EFNB3, EPHB3, RGS3 |
| U | TCF4 Transcriptional Regulation and Neuronal Differentiation | LDB1, LMO3, SSBP2, SSBP3, TCF4, TLE3, TLE4, TLE5 | |
| V | Augmin Microtubule Nucleation and Mitotic Spindle Assembly | HAUS3, HAUS4, HAUS5, HAUS6, HAUS7, NEDD1, NUMA1, TUBG1 | |
| W | SLIT–ROBO Signaling and Axon Guidance | MYO6, PAK4 | PAK3, PAK6, ROBO1, SLIT1, SLIT2, TOM1 |
| X | Endocannabinoid Hydrolysis and Lipid Metabolism | ABHD6, MGLL | ABHD12, AGK, FAAH, NCEH1, TIMM22 |
| Y | Latrophilin Adhesion Receptors and Transsynaptic Signaling | GNG5, PDCL | ADGRL1, ADGRL3, GNB2, TENM1 |
| Z | $Na^+$/$K^+$ ATPase Ion Transport and ER Stress Regulation | | ATP1A3, ATP1B1, ATP1B3, CISD2, WFS1 |

**Figure S23. STRING interaction network modules identified from proteomic analysis of female fetal brain development in the absence of prenatal cannabis exposure (Controls T2 vs. T1).** (A–NN) STRING-derived proteomic interaction subnetworks constructed from significantly altered proteins identified in female Controls T2 vs. T1. Blue nodes indicate proteins downregulated in T2 Controls, whereas red nodes indicate proteins upregulated in T2 Controls. The table summarizes the major functional modules represented within each interaction cluster.

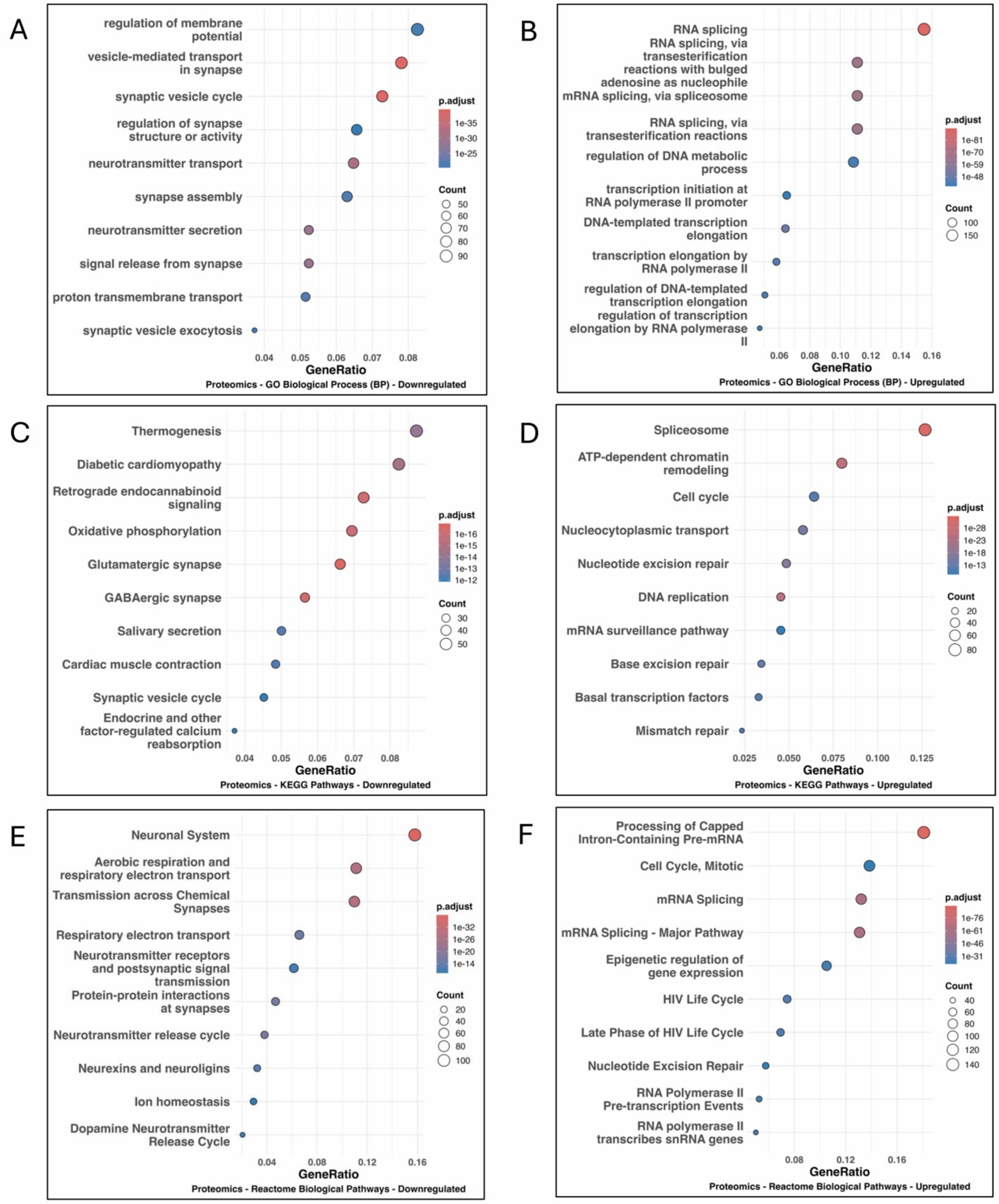


**Figure S24. Proteomic enrichment associated with development in male T2 Cases vs. T1 Cases.** Gene Ontology (Biological Process) enrichment analysis of downregulated (**A**) and upregulated (**B**) proteins. Dot plots show enriched terms ranked by gene ratio, with point size indicating protein count and color representing adjusted P value. Downregulated proteins were enriched for synaptic vesicle cycling, neurotransmitter transport, and synapse organization, whereas upregulated proteins were enriched for RNA splicing, transcriptional regulation, and DNA metabolic processes. KEGG pathway enrichment analysis of downregulated (**C**) and upregulated (**D**) proteins. Downregulated pathways include oxidative phosphorylation, retrograde endocannabinoid signaling, and synaptic transmission pathways, whereas upregulated pathways were enriched for spliceosome, chromatin remodeling, and cell cycle processes. Reactome pathway enrichment analysis of downregulated (**E**) and upregulated (**F**) proteins. Downregulated pathways were enriched for neuronal system, synaptic transmission, and respiratory electron transport, whereas upregulated pathways were enriched for cell cycle progression, RNA processing, and DNA repair pathways. Dot plots show enriched terms ranked by gene ratio, with point size indicating gene count and color representing adjusted P value.
